# Structural Basis of Lipid Membrane Binding by Human Ferlins

**DOI:** 10.1101/2025.01.25.634844

**Authors:** Constantin Cretu, Aleksandar Chernev, Csaba Zoltan Kibedi Szabo, Vladimir Pena, Henning Urlaub, Tobias Moser, Julia Preobraschenski

## Abstract

Ferlins, ancient membrane proteins with a unique architecture, are central to multiple essential, Ca^2+^-dependent vesicle fusion processes. Despite numerous functional studies and their link to burdening human diseases, a mechanistic understanding of how these multi-C_2_ domain proteins interact with lipid membranes to promote their remodeling and fusion is currently lacking. Here, we elucidate the near-complete cryo-electron microscopy structures of human myoferlin and dysferlin in their Ca^2+^ and lipid-bound states. We show that ferlins adopt compact, ring-like tertiary structures achieved upon membrane binding. The top arch of the ferlin ring, comprising the C_2_C-C_2_D region, is rigid and varies little across the observed functional states. In contrast, the N-terminal C_2_B and the C-terminal C_2_F-C_2_G domains cycle between alternative conformations and, in response to Ca^2+^, close the ferlin ring, promoting tight interaction with the target membrane. Probing key domain interfaces validates the observed architecture and informs a model of how ferlins engage lipid bilayers in a Ca^2+^- dependent manner. This work reveals the general principles of human ferlin structures and provides a framework for future analyses of ferlin-dependent cellular functions and disease mechanisms.

## Introduction

Excitable cells rely on precisely timed Ca^2+^ signals to trigger exocytosis in neural synapses and contraction in muscle cells (Clapham, 2007, Jahn & Fasshauer, 2012, Kuo & Ehrlich, 2015, Luan & Wang, 2021). Less is known how the uncontrolled influx of Ca^2+^ through a large membrane lesion promotes a rapid acute response (Cooper & McNeil, 2015). C_2_- domain proteins are key molecular players utilizing the incoming Ca^2+^ signal to tether intracellular vesicles to their target membranes and ultimately promote their fusion by coordinated binding to phospholipids and other factors (Rizo, 2022, Rizo & Sudhof, 1998). Beyond the well-studied synaptotagmins, which feature only two consecutive C_2_ domains (Rizo & Sudhof, 1998, Sudhof, 2013), ferlins, predicted to comprise up to eight C_2_ domains (Dominguez, McCord et al., 2022), play a pivotal role in mediating these processes (Bansal & Campbell, 2004, Cooper & Head, 2015, Cooper & McNeil, 2015, Pangrsic, Reisinger et al., 2012) and are critically needed at multiple steps (Chen, Monga et al., 2024, Cooper & McNeil, 2015, Pangrsic et al., 2012, Rizo, 2022).

Ferlins, such as dysferlin (FER1L1), myoferlin (FER1L3), and otoferlin (FER1L2), form an ancient group of C_2_ domain Ca^2+^-sensing proteins present in almost all eukaryotic lineages (Bansal & Campbell, 2004, Han & Campbell, 2007, Pangrsic et al., 2012, Petit, Bonnet et al., 2023). Dysferlin and myoferlin are highly expressed in skeletal and heart muscle cells, which are prone to membrane injuries during contractions (Bansal & Campbell, 2004, Cooper & McNeil, 2015, Pramono, Tan et al., 2009). Distributed at the sarcolemma, its specialized internal structures and various endomembrane vesicles (Paulke, Fleischhacker et al., 2024), dysferlin has been proposed to play major roles in Ca^2+^-dependent membrane resealing, in the biogenesis and maintenance of the transverse-tubules system (Kerr, Ziman et al., 2013, Paulke et al., 2024), and other trafficking pathways (Cooper & McNeil, 2015, Glover & Brown, 2007, Han & Campbell, 2007). The central physiological role of dysferlin in maintaining the integrity of muscle cell membranes is evident from over 400 disease-causing *DYSF* mutations identified in limb-girdle muscle dystrophy type 2B (LGMD2B) and Miyoshi myopathy (MM) – rare autosomal recessive muscle wasting disorders characterized by deficiencies in sarcolemma repair (Bansal, Miyake et al., 2003, Cooper & Head, 2015). Similar to dysferlin, myoferlin (Cooper & McNeil, 2015, Davis, Delmonte et al., 2000, de Morree, Hensbergen et al., 2010), has been linked to diverse membrane remodelling and organelle repair events, including in other cell types, as well as to myoblast membrane fusion during myogenesis (Davis et al., 2000, Doherty, Cave et al., 2005). However, unlike dysferlin, myoferlin has recently also been found to be overexpressed in various human cancers and the altered vesicular trafficking and function of myoferlin has been linked to cancer cell proliferation, metastasis, and resistance to chemotherapy (Cooper & McNeil, 2015, Gupta, Yano et al., 2021, Zhang, Li et al., 2018). The closely related otoferlin is mainly expressed in sensory hair cells of the inner ear, and several hundred pathogenic mutations cause the deafness DFNB9 (Moser & Starr, 2016, Petit et al., 2023, Santarelli, del Castillo et al., 2015, Vona, Rad et al., 2020, Yasunaga, Grati et al., 1999). In mechanistic terms, it has been suggested that otoferlin is involved in Ca^2+^-sensing for synaptic vesicle fusion (Johnson & Chapman, 2010, Michalski, Goutman et al., 2017, Roux, Safieddine et al., 2006) and replenishment (Pangrsic, Lasarow et al., 2010, Vogl, Panou et al., 2016), as well as exocytosis-endocytosis coupling (Duncker, Franz et al., 2013, Jung, Maritzen et al., 2015, Kroll, Jaime Tobon et al., 2019, Strenzke, Chakrabarti et al., 2016), and its mutations disrupt synaptic sound encoding, resulting in an auditory synaptopathy (Moser & Starr, 2016, Pangrsic et al., 2012). Despite the recent insights into the individual roles of ferlins and them representing targets for gene therapy or drug discovery (Al-Moyed, Cepeda et al., 2019, Gupta et al., 2021, Llanga, Nagy et al., 2017, Moser, Chen et al., 2024, Zhang et al., 2018), the underlying molecular mechanisms, especially concerning how they promote membrane remodelling and fusion through concerted Ca^2+^ and phospholipid binding, have remained largely unresolved, in part, due to limited structural information on their active states.

In structural terms, ferlins arguably possess the most unique and complex architecture among the known Ca^2+^-sensitive C_2_-domain proteins (Lek, Evesson et al., 2012, Pangrsic et al., 2012). The ferlin amino-terminal (N-terminal) cytoplasmic domain is predicted to contain up to eight β-sandwich C_2_ domains (Dominguez et al., 2022). The ferlin C_2_ domains are thought to be connected by unstructured linker regions and are followed by a carboxy-terminal (C-terminal), single-pass transmembrane region, anchoring the proteins to cellular membranes. In addition, type-I ferlins, such as dysferlin and myoferlin, have two accessory domains with an unknown function, DysF and FerA, inserted between the third and fourth C_2_ domains (Dominguez et al., 2022). Generally, similar to synaptotagmins (Chapman, 2008, Rizo, 2022, Rizo & Sudhof, 1998), the individual C_2_ domains of ferlins have variable Ca^2+^ and phospholipid binding activities (Abdullah, Padmanarayana et al., 2014, Marty, Holman et al., 2013), with few exceptions (otoferlin’s C_2_A domain (Helfmann, Neumann et al., 2011, Johnson & Chapman, 2010)). However, previous studies failed to clarify how full-length ferlins are precisely organized to interact with lipid membranes and which of their structural motifs are critical for membrane binding. Consequently, it remains poorly understood how ferlins act on lipid membranes to promote their remodelling and fusion, and how Ca^2+^- sensitive conformational rearrangements mediate these processes (Lek et al., 2012, Pangrsic et al., 2012, Xu, Pallikkuth et al., 2011).

Herein, we leverage structural biology approaches and functional analyses to obtain near complete and high-resolution cryo-EM models of the two largest human ferlins, myoferlin and dysferlin, in their Ca^2+^ and lipid-bound states. Besides revealing the intricate organization of these essential vesicle trafficking factors, our ferlin structures shed light on how the C_2_ and accessory motifs engage lipid bilayers in a coordinated manner. We further advance a model of how ferlins cycle between alternative conformational states to transiently bind lipid membranes and facilitate their remodelling and fusion.

## Results

### 2.4 Å cryo-EM structure of membrane-bound human myoferlin

To obtain the first complete structure of a human ferlin, we established the heterologous expression and purification of full-length myoferlin and dysferlin (**Fig 1A and Appendix Fig S1A**). In addition to the membrane-anchored constructs, we expressed the entire cytosolic region of the ferlins, comprising all C_2_ and accessory motifs (**Appendix Fig S1A**). Generally, the detergent or liposome reconstituted samples were homogenous and retained their ability to bind Ca^2+^ and negatively charged lipid membranes, as previously reported for the individual ferlin domains (Abdullah et al., 2014, Marty et al., 2013, Padmanarayana, Hams et al., 2014) (**Appendix Fig S1B and S1D-G**). However, in contrast to an early model of dysferlin (Xu et al., 2011) and a recent report (Huang, Grandinetti et al., 2024), we did not observe a significant tendency of the proteins to dimerize through the C_2_ domains, in mass photometry measurements and size-exclusion chromatography, suggestive of the type-I ferlins being organized as a monomers in solution (**Appendix Fig S1B-C and S1H**).

**Figure 1.**
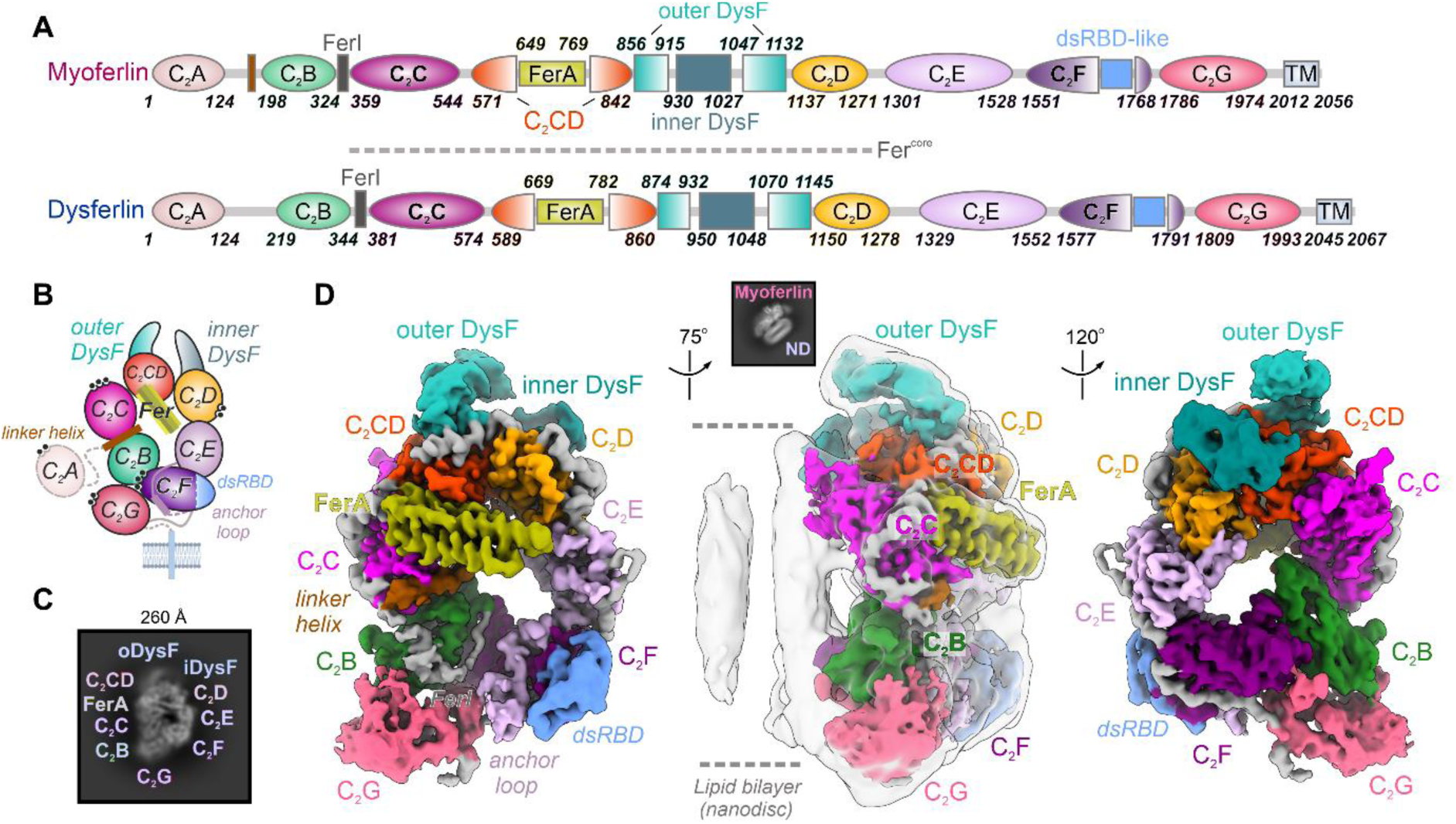
Cryo-EM structure of human myoferlin bound to a model lipid bilayer (nanodisc). **A** Schematic depicting the structure-based domain composition of human myoferlin and dysferlin. The C_2_ domains of myoferlin and dysferlin are shown as coloured ellipses, whereas the DysF, FerA and the transmembrane region (TM) are displayed as boxes. The linker helix between C_2_A and C_2_B is depicted as a brown box, whereas the remaining linker regions are coloured grey. dsRBD-like: the double-stranded RNA-binding-like subdomain of C_2_F. **B** Schematic representation of the soluble myoferlin-lipid nanodisc cryo-EM structure. The transmembrane helix is depicted for orientation purpose and is absent from the soluble expression construct. The modelled Ca^2+^ ions are indicated as black dots. **C** Typical top-view of lipid-bound soluble myoferlin. Myoferlin domains, which are visible in the 2D class average, are indicated. **D** The cryo-EM map of lipid-bound soluble myoferlin displayed in three different orientations. The individual ferlin domains are color-coded as in A. The soluble myoferlin map has been locally scaled and low-pass filtered to 2.8 Å, whereas the nanodisc density (middle panel, contoured in gray around the ferlin cytosolic region) has been low-passed to 8 Å resolution to allow the visualization of the ordered lipid regions (see also **Appendix Fig S2-S6**). The inset (middle panel) shows a typical side-view class average, consistent with an asymmetric membrane recognition mechanism.

Initial attempts at single-particle cryo-EM imaging of lipid-free ferlins revealed the intrinsic flexibility of the N-terminal (C_2_A-C_2_B) and C-terminal domains (C_2_F-C_2_G), which limited the resolution of the maps. We, therefore, hypothesized that these more dynamic motifs, all predicted to engage lipid bilayers (Abdullah et al., 2014, Johnson & Chapman, 2010, Kwok, Otto et al., 2023, Marty et al., 2013), would become organized upon membrane binding. In our attempts to identify an optimal membrane system, we observed that the cytosolic region of myoferlin (residues 1-1997) formed stable complexes with large MSP2N2 (membrane scaffold protein 2 N2)-based lipid nanodiscs, comprising anionic phospholipids and chosen to accommodate all interacting domains (Cannon, Sarsam et al., 2023) (**Fig 1D and Appendix Fig S1B-C**). We further stabilized the myoferlin-nanodisc complexes through glutaraldehyde crosslinking (Kastner, Fischer et al., 2008) and imaged four different protein-lipid samples, which were assembled on membranes containing PS (Phosphatidylserine) alone or combined PS and PI(4,5)P_2_ (Phosphatidylinositol-4,5-bisphosphate) (**Appendix Fig S2-S5**). Computational image sorting allowed us to identify intact particles and reconstruct near-complete cryo-EM maps of the lipid-bound myoferlin (**Fig EV1, Appendix Fig S2-S5 and Movie EV1**). These well-resolved maps refined in 3D to a 2.4-2.9 Å global resolution, allowing us to confidently assign and build nearly all C_2_ and accessory domains, apart from the flexible N-terminal C_2_A domain (**Fig 1D, Fig EV1, Appendix Fig S2-S5 and Appendix Tables S1-S4**). Myoferlin interacts with the MSP2N2 nanodisc through multiple binding motifs, all projecting on one side and forming well-defined protein-lipid contact sites (**Fig 1D, Fig EV1 and Fig EV2**). As expected, the remaining lipid membrane is largely disordered in these structures (**Fig 1D and Fig EV2**).

### Complex tertiary interfaces organize the lipid-bound state of myoferlin

Although predicted to adopt an extended, “beads on a string”-like topology (Dominguez et al., 2022, Leclère & Dulon, 2023), the overall cryo-EM map of the lipid-bound myoferlin revealed a surprisingly compact domain architecture (**Fig 1B and 1D**). As observed in 2D class averages (**Fig 1C, Appendix Fig S2B, Fig S4C, Fig S5B and S5E**), the individual structural motifs of myoferlin are distributed in an almost coplanar manner around a ∼30 Å central cavity, describing an elliptic ring that spans ∼150 Å and ∼90 Å along its long and short axes, respectively. One side of the composite ring engages the membrane bilayer at four distinct contact sites, covering the entire nanodisc perimeter (**Fig 1D and Fig EV2**). In contrast, the solvent-facing side of myoferlin harbors no lipid recognition motifs, consistent with the presence of a single and nearly planar membrane-binding surface (**Fig 1D and Fig EV2 A-F**).

The top region of the ferlin ring (**Fig 1A-B and 1D**) is formed by the structurally more rigid C_2_C (residues 358-544), C_2_CD-FerA (residues 571-842) and C_2_D (residues 1137-1271) domains, which we refer to as the ferlin core module (Fer^core^, **Fig EV1D-E and Movie EV2**). The inner and outer DysF domains (**Fig 1A and 1D**), each comprising two long β-strands, are closely linked to the Fer^core^, through C_2_D and C_2_CD, respectively, and separated by ∼14 Å (**Fig 1A**, **Fig 1D**, **Fig 2D and Fig EV1F**). Opposite of the Fer^core^, the N-terminal C_2_B (residues 198-324), the C-terminal C_2_F (residues 1551-1768), and the membrane-proximal C_2_G (residues 1786-1974) approach each other in 3D and pack closely, despite being separated by more than 1200 residues (**Fig 1D**, **Fig 2A-B, Fig EV1 and Movie EV2**). Unlike the Fer^core^ motifs, these C_2_ domains are engaged in fewer and transient interdomain contacts, in part mediated by loop regions or unstructured elements (**Fig 2B-C and Movie EV2**). C_2_B contributes to the largest number of contacts and is surrounded from three sides by the top loops of C_2_F (L1, L3 and L4), the long β6-β7 subdomain of C_2_G (residues 1906-1944, **Fig EV1J-K**) and the C_2_C domain of Fer^core^ (**Fig 2B-C**). Additional contacts are provided by the membrane-binding β-hairpin of C_2_G (residues 1859-1879), inserted between the β4-β5 strands, and the upstream linker helix (residues 175-190), stacked between C_2_B and C_2_C (**Fig 2B-C, Fig EV1A and EV1J-L**). The conserved but largely unstructured FerI motif (residues 325-357) appears to fold between C_2_F, C_2_G and C_2_B, before reaching C_2_C, likely also stabilizing the composite interface (**Fig 1A**, **Fig 2A-B and Fig EV1A**). Finally, the top and bottom arches of the ferlin ring are connected through the C_2_E domain, which interacts with both C_2_F and C_2_G through its extended anchor loop (residues 1447-1498), as well as with the upstream C_2_D through its loop 4 (L4, **Fig 2A**, **Fig 2D-E and Fig EV1H**).

**Figure 2.**
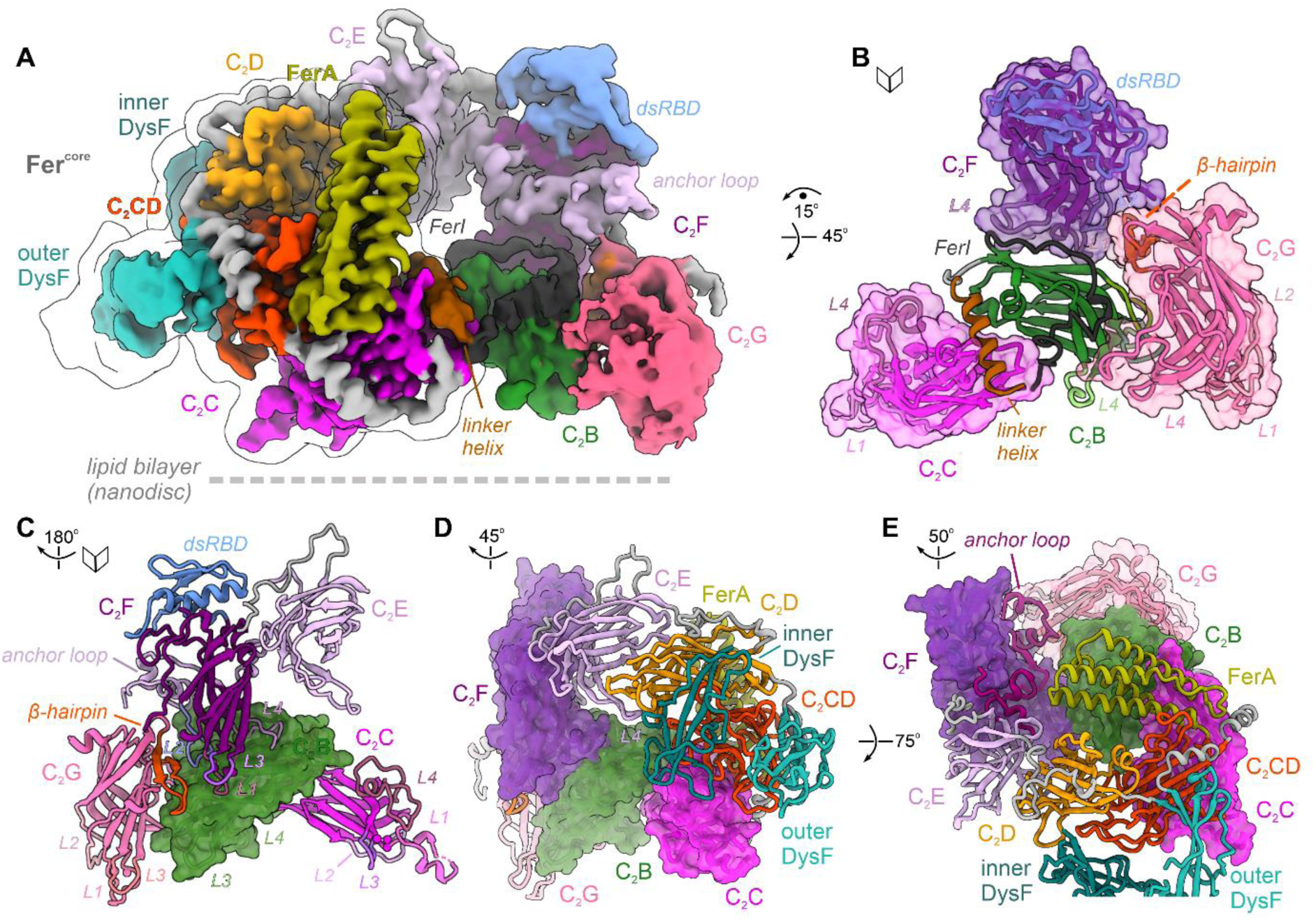
The multipartite structural organization of lipid-bound myoferlin. **A** Overall cryo-EM map of lipid-bound soluble myoferlin (residues 1-1997), viewed from its N-terminal side. The map has been color-coded after the modelled subunits and the key structural motifs are indicated. The Fer^core^ module of myoferlin (the C_2_C-C_2_D region) is contoured in light gray. **B** The multiple tertiary interfaces between the N-terminal C_2_B-C_2_C and the C-terminal C_2_F-C_2_G, observed in the lipid-bound myoferlin structure. Additional structural elements, such as the linker helix and the FerI motif, bridging the C_2_B and C_2_C domains, and the β-hairpin subdomain of C_2_G, are indicated. The C_2_ domains, except C_2_B, are depicted as transparent solvent-excluded surfaces with the domain model fitted inside. **C** The C_2_B domain orientation in the lipid-bound state appears to be induced by its shared interfaces with C_2_C, C_2_F and C_2_G. C_2_B is shown in a surface representation. The top loops (L1-L4) of the interacting domains are indicated. **D** The peripheral inner and outer DysF motifs of myoferlin, viewed from the membrane-interacting side. The C_2_C, C_2_F and C_2_G domains are depicted as surfaces. Note that the two DysF motifs are connected to the Fer^core^ module through C_2_D (orange) and C_2_CD (red). **E** C_2_E connects the Fer^core^ module to the C-terminal C_2_F domain through a long insertion loop (the “anchor loop”). C_2_E is depicted as cartoon, whereas the C_2_B-C_2_C and C_2_F-C_2_G are shown as surfaces. The anchor loop inserts between the β6 and β7 strands of C_2_E (see also **Appendix Fig S4h**). At the same time (**Fig 2D**), its L4 loop is oriented towards C_2_D.

### The rigid ferlin core module is stabilized by a new C_2_-like accessory domain

Consistent with limited proteolysis experiments in cultured cells (Woolger, Bournazos et al., 2017), the multipartite organization of the myoferlin ring appears to be centered around the tightly packed Fer^core^ module (**Fig 3A-B and Movie EV2**). Fer^core^ covers almost one half of the ferlin ring and is distributed symmetrically at the two ends of the arch (**Fig 3A-B**). In between C_2_C and C_2_D, four distinctive motifs are observed (**Fig 3A-B**): (i) the FerA domain (residues 649-769), located in the proximity of C_2_C and folded as a four-helix bundle (Harsini, Chebrolu et al., 2018); (ii) the outer DysF domain (also known as N-DysF, residues 856-915 and 1047-1132), occupying a more central location; (iii) the inner DysF domain (or C-DysF, residues 930-1027), inserted between the two β-strands of the outer DysF (Patel, Harris et al., 2008, Sula, Cole et al., 2014) and oriented towards the membrane surface; (iv) a previously undetected C_2_-like motif, which we denoted as the C_2_CD domain (residues 571- 648 and 770-842).

**Figure 3.**
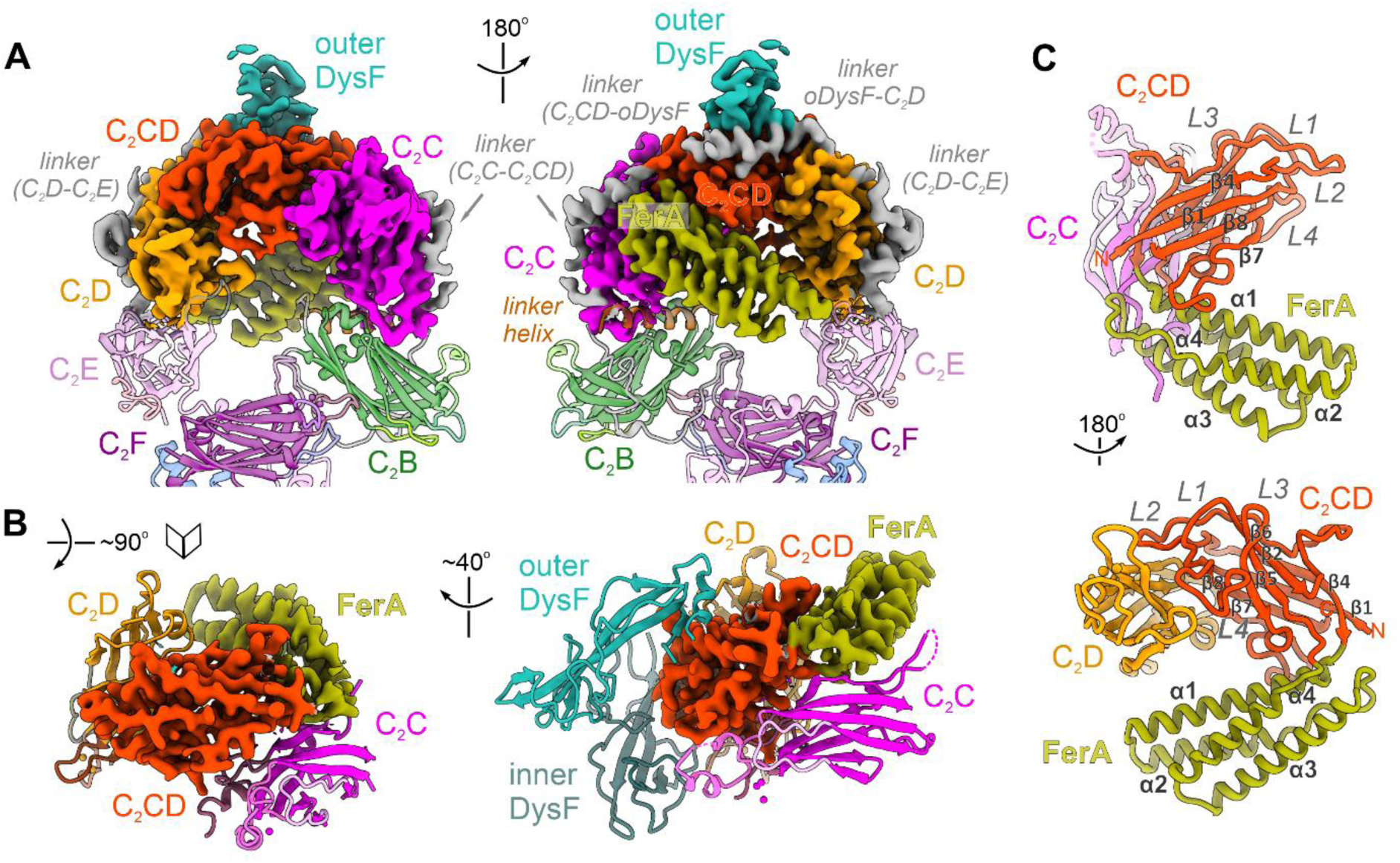
Structure of the rigid Fer^core^ module. **A** Structure of the Fer^core^ module, spanning the top ferlin arch. Fer^core^ comprises the C_2_C-C_2_D region (color-coded cryo-EM map after the modelled subunits) and includes a new C_2_-like structural domain, the C_2_CD domain (red). The neighbouring domains, C_2_B and C_2_E-C_2_F are shown in cartoon representation. The modelled linker regions, connecting the C_2_ domains, are coloured grey. **B** Interaction interfaces between C_2_CD and its neighbouring FerA, C_2_C and C_2_D domains. C_2_CD and FerA are depicted as cryo-EM maps, whereas C_2_C, C_2_D, outer and inner DysF are shown as cartoons. **C** Structure of myoferlin’s C_2_CD and FerA domains. The seven β-strands of C_2_CD and the four α-helices of FerA are indicated.

The newly identified C_2_CD domain is present in all ferlins, including dysferlin and otoferlin (**Fig 3C**, **Fig 6C-D, Appendix Fig S9A-D and S10A-D**), and spans the distance between C_2_C and C_2_D, thereby appearing to “glue” the Fer^core^ module together. Like all other C_2_ domains of myoferlin, C_2_CD consists of two β-sheets and has a type-II topology (Dominguez et al., 2022). However, unlike typical C_2_ domains, its second β-sheet comprises only three β- strands (**Fig 3C and Appendix S10B**), the top loops (L1-L4) are extended, lack conservation and engage in tertiary contacts with C_2_C, inner DysF and C_2_D, instead of being available for Ca^2+^ and/or phospholipid binding (see also **Appendix Fig S9E-G**). C_2_CD is closely associated with the four-helix bundle of FerA, inserted between its β4-β5 strands (**Fig 3C**). Interestingly, FerA does not engage the lipid nanodisc and exits the C_2_CD domain opposite from the membrane-binding surface (**Fig 3B-C**, see also **Appendix Fig S9E-G and Fig S11**), partially closing the central cavity of myoferlin as it aligns diagonally along the C_2_C-C_2_E axis, without reaching C_2_E. We suggest that the large contact interfaces of C_2_CD with the remaining Fer^core^ motifs (∼4000 Å^2^) indicate a primary role as a repurposed C_2_ domain packing platform (**Appendix Fig S11A-C**), contrasting the Ca^2+^-sensing and lipid binding functions of other myoferlin C_2_ domains (see also **Fig 4A and Fig. 5**).

**Figure 4.**
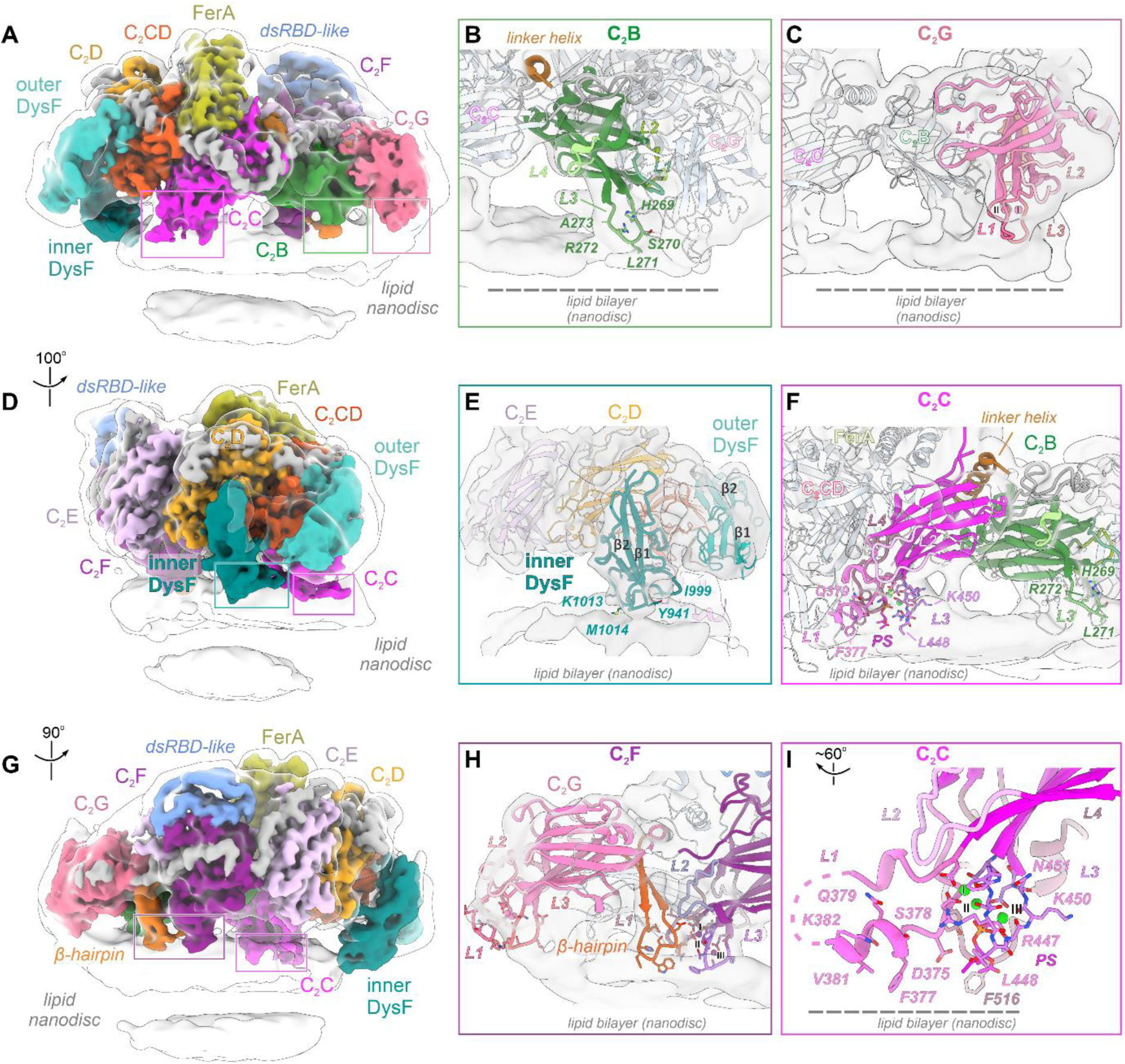
The composite lipid membrane recognition interface of human myoferlin. **A** The lipid nanodisc contact interfaces of the C_2_B and C_2_C domains. The focused myoferlin (1-1997) cryo-EM map (color-coded after the modelled subunits) is shown together with the 8 Å low-passed overall map (**Appendix Fig S2C**). The ordered lipid nanodisc regions interact with myoferlin at multiple contact sites, defined by C_2_ (C_2_B, C_2_C, C_2_F, C_2_G) and accessory (inner DysF, β-hairpin of C_2_G) domains. **B** C_2_B interacts with lipid bilayers independent of Ca^2+^ binding. Although no bound Ca^2+^ ions were observed and C_2_B lacks conservation of the typical aspartate residues involved in divalent cation coordination, the domain engages the lipid bilayer (nanodisc) through its L3 top loop. C_2_B residues directly interacting or located close to the lipid bilayer are shown as sticks. **C** Close-up of C_2_G’s lipid nanodisc contact interface. The top loops of C_2_G (L1 and L3) project close to C_2_B’s binding site. The two modelled Ca^2+^ ions are indicated. **D** The peripheral inner DysF motif engages the lipid bilayer close to C_2_C’s binding site. The lipid nanodisc density is coloured in grey, whereas the cryo-EM map of myoferlin is coloured-coded as in **A. E** Close-up of the inner DysF domain interacting with the lipid nanodisc. Inner DysF’s loop residues located close to the nanodisc surface are shown as sticks, for the sake of orientation. Note that the outer DysF and Ca^2+^-bound C_2_D, flanking the inner DysF, do not bind the lipid nanodisc. **F** C_2_C establishes extensive, both Ca^2+^-dependent and independent, contacts with lipid bilayers. C_2_C’s membrane binding involves hydrophobic and basic residues of the L1, L3 and L4 loops, as well as direct Ca^2+^-mediated phospholipid headgroup coordination. The bound phosphatidylserine (PS) moiety and key interface residues are depicted as sticks. **G-H** The C-terminal C_2_F and C_2_G bind lipid bilayers through a composite interface. The key lipid-binding motifs (the L3 loops of C_2_F and C_2_G, the β-hairpin of C_2_G) are indicated and the membrane-facing residues are shown as sticks. **I** The top L1 loop of C_2_C interacts with the lipid bilayer through an amphipathic insertion helix. Note the virtually parallel orientation of L1’s amphipathic helix, with its hydrophobic residues engaging the lipid bilayer (see also **Fig 4F**). The Ca^2+^-coordinating residues and the bound PS headgroup are shown as sticks.

**Figure 5.**
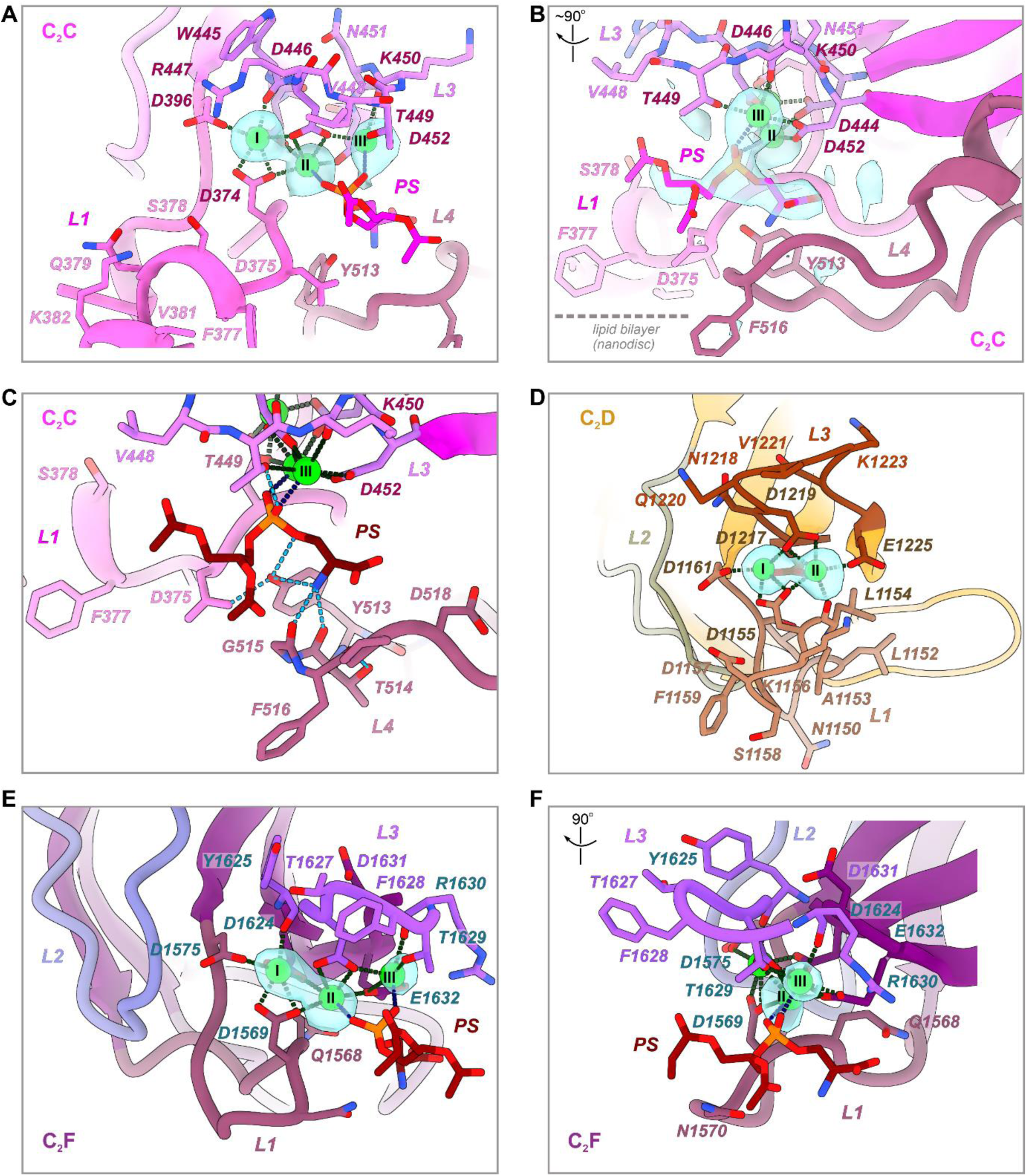
Ca^2+^ and phospholipid-binding sites modelled in the lipid-bound structure of myoferlin. **A** Three Ca^2+^ ions and a phosphatidylserine (PS) headgroup bind the acidic pocket of C_2_C. The normalized cryo-EM difference map (cyan, 6.5σ) is contoured around the modelled Ca^2+^ ions. Ca1(I) and Ca2(II) have a hexadentate coordination. Ca3(III) is coordinated by both L3 loop residues and a PS coordinating oxygen. The coordination bonds are depicted as dashed lines. **B** The modelled PS headgroup contributes to C_2_C Ca^2+^ ion coordination. The cryo-EM difference map (cyan, 6.5σ) is contoured around PS and two Ca^2+^ ions (Ca2 and Ca3). See also **Fig EV1M**. **C** Molecular recognition of the PS headgroup by myoferlin’s C_2_C domain. The modelled PS is shown as sticks (dark red). Ca^2+^ coordination bonds are depicted as black dashed lines, whereas the polar interactions of PS with the L3 and L4 loops are coloured cyan. **D** The Ca^2+^-binding sites of the C_2_D domain. The cryo-EM difference map (cyan, 6.5σ) is contoured around the modelled Ca^2+^ ions. The two Ca^2+^ ions of C_2_D (Ca1, Ca2) have a hexadentate coordination achieved through interaction with both L1 and L3 loop residues. **E** Ca^2+^ and phospholipid binding sites of the C_2_F domain. The cryo-EM difference map (cyan, normalized, 5σ) is contoured around the three modelled Ca^2+^ ions (Ca1-Ca3). Two Ca^2+^ ions (Ca1 and Ca2), bound to C_2_F, have a hexadentate coordination. The Ca3 ion is coordinated by the L3 loop and PS oxygen atoms. C_2_F residues contributing to Ca^2+^ coordination are depicted as sticks and coloured in teal. **F** Several hydrophobic (F1628, Y1625) and basic (R1630) residues, located at the tip of the L3 loop, engage the lipid bilayer, possibly independent of Ca^2+^ coordination. The PS headgroup and the modelled Ca^2+^ ions are represented as in **E.**

**Figure 6.**
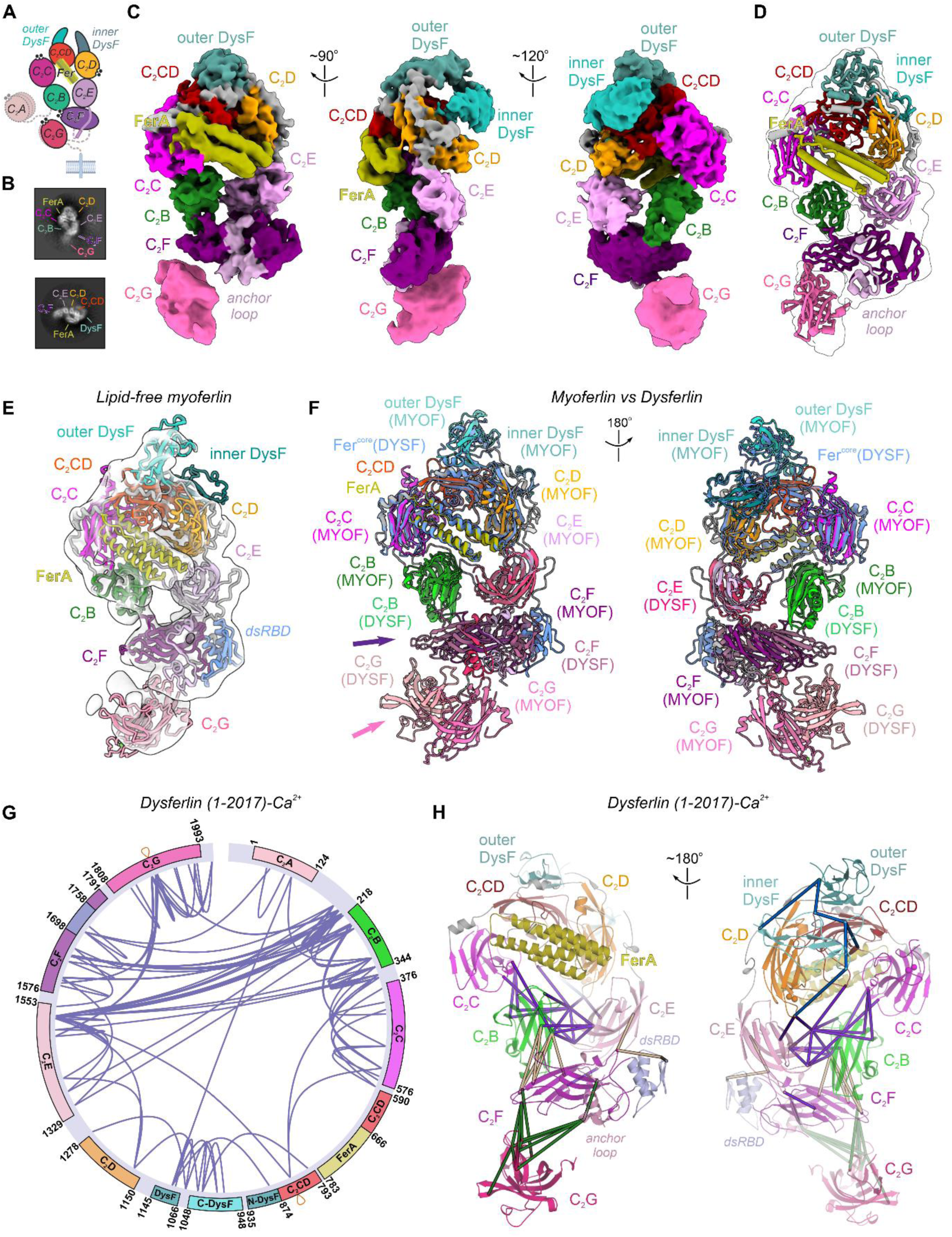
Cryo-EM structures of human dysferlin and myoferlin in the lipid-free states. **A** Schematic representation of dysferlin’s cryo-EM structure. The transmembrane region, absent from the expressed cytosolic domain of dysferlin (residues 1-2017), is shown for the sake of orientation. **B** Two representative, reference-free 2D class averages of dysferlin. The visible dysferlin domains are indicated. **C** The cryo-EM map of the lipid-free and Ca^2+^-bound dysferlin (1-2017), rendered in three different orientations. The individual ferlin domains are colour-coded as in **A**. The map has been locally scaled and low-pass filtered to 5 Å (see also **Appendix Fig S6-S8**). **D** The overall model of dysferlin (1-2017) showing the observed tertiary interfaces between C_2_B, C_2_E, C_2_F, and C_2_G. The final model of dysferlin (1-2017) is fitted within the overall map. **E** The cryo-EM structure of lipid-free and Ca^2+^-bound myoferlin (1-1997). The individual ferlin domains are colour-coded and the overall map has been low-pass filtered to 10 Å (see also **Fig EV3E**) to better visualize the dynamic C_2_G domain. **F.** Structural superposition between dysferlin (1-2017) and myoferlin (1-1997) in the lipid- free states. Consistent with it being rigid, the Fer^core^ module is virtually identical in the two lipid-free ferlin structures. Note that, compared to the C-terminal C_2_F-C_2_G, C_2_B and C_2_E adopt similar poses in the lipid-free structures (see also **Fig 6C**). **G** Circular plot displaying all the observed intramolecular crosslinks between dysferlin’s structural motifs. The BS3 chemical crosslinking data at FDR (false discovery rate) of 1% were not filtered by score or spectral count (see also **Appendix Fig. S14A-C and Dataset EV1**). **H** Selected interdomain BS3 crosslinks mapped onto the lipid-free state of dysferlin. The displayed Euclidean distances (coloured lines) were calculated between the Cα atoms of the crosslinked lysine residues and are all below the 35 Å theoretical distance threshold (see also **Appendix Fig S14A-B and Dataset EV1**).

### The composite membrane-binding interface of human myoferlin

Despite numerous structural studies of other C_2_-domain proteins, such as synaptotagmins (Rizo, 2022, Rizo, Sari et al., 2022, Schauder, Wu et al., 2014, Seven, Brewer et al., 2013), there has not been direct experimental evidence for how a multi-C_2_ domain protein interacts with lipid bilayers through the combined binding of Ca^2+^ and phospholipids. In particular, the relative orientation of all the membrane-binding domains, their lipid-interacting motifs, and their sensitivity to Ca^2+^ (or lack thereof), remained speculative (Arac, Chen et al., 2006, Honigmann, van den Bogaart et al., 2013, Rizo et al., 2022, Seven et al., 2013). The cryo-EM structure of human myoferlin elucidates the organization of its seven C_2_ and accessory domains upon concerted membrane-binding, enabling us to identify the precise motifs involved in lipid recognition.

In the nanodisc-bound structure (**Fig 4A, 4D and 4G**), the cytosolic region of myoferlin engages the lipid surface primarily through the C_2_B-C_2_G, C_2_C, the inner DysF and C_2_F-C_2_G domains, establishing a total of four membrane anchoring points. At the first binding site, we observe that the N-terminal C_2_B (**Fig 4B**) faces the lipid membrane with its concave surface at a ∼28° tilt angle. In this side-orientation, C_2_B projects the loop 3 (L3) nearly perpendicular to the membrane plane, binding the bilayer through several hydrophobic (L271, A273) and polar (S270, R272) residues located at the L3 tip. The remaining residues of the L1-L3 loops do not participate in Ca^2+^-coordination, as no ion densities were observed in the cryo-EM maps. Instead, they establish contacts with C_2_G, which contributes to the same contact site, suggestive of cooperative membrane binding (**Fig 4B-C**). Like C_2_B, the C-terminal C_2_G faces the nanodisc with its concave side tilted by ∼70° (**Fig 4C and Fig 4H**), directly interacting with a small membrane patch through polar and hydrophobic residues of L3 (K1892, F1893, L1895). However, in contrast to C_2_B, the acidic residues in L1 and L3 of C_2_G are conserved and, likely, involved in the coordination of two Ca^2+^ ions (**Fig 4C and Fig EV1J-L**). Surprisingly, our structures revealed that an additional membrane anchoring point (**Fig 4D-E**) is provided by the inner DysF motif. Located between C_2_CD and C_2_D, the inner DysF forms direct membrane contacts (**Fig 4D-E and Fig EV1F**), likely, through hydrophobic (I997, I999, P1000, P1001), basic (K1013, K1016), and aromatic residues (Y1015), providing first evidence, to our knowledge, of its role as a generic class of lipid-binding motifs (Kaur, de la Ballina et al., 2023). The positioning of the inner DysF at the membrane surface appears to be influenced by its interactions with the L2 of C_2_CD and the L1 of the neighbouring C_2_D (**Fig 2E and Fig 4D**). While engaging the inner DysF, C_2_D is positioned ∼20 Å away from the lipid nanodisc, where it still coordinates two Ca^2+^ ions through its L1 and L3 (**Fig 4E**, **Fig 5D and Fig EV1G**) and interacts through the L3 loop with the L4 of the downstream C_2_E (**Fig 2D-E**). Overall, these observations suggest that the C_2_D domain might guide the inner DysF to the membrane and couple this binding event to the recruitment of C_2_F-C_2_G via C_2_E, possibly in a Ca^2+^-dependent manner (see also **Fig 7E**).

**Figure 7.**
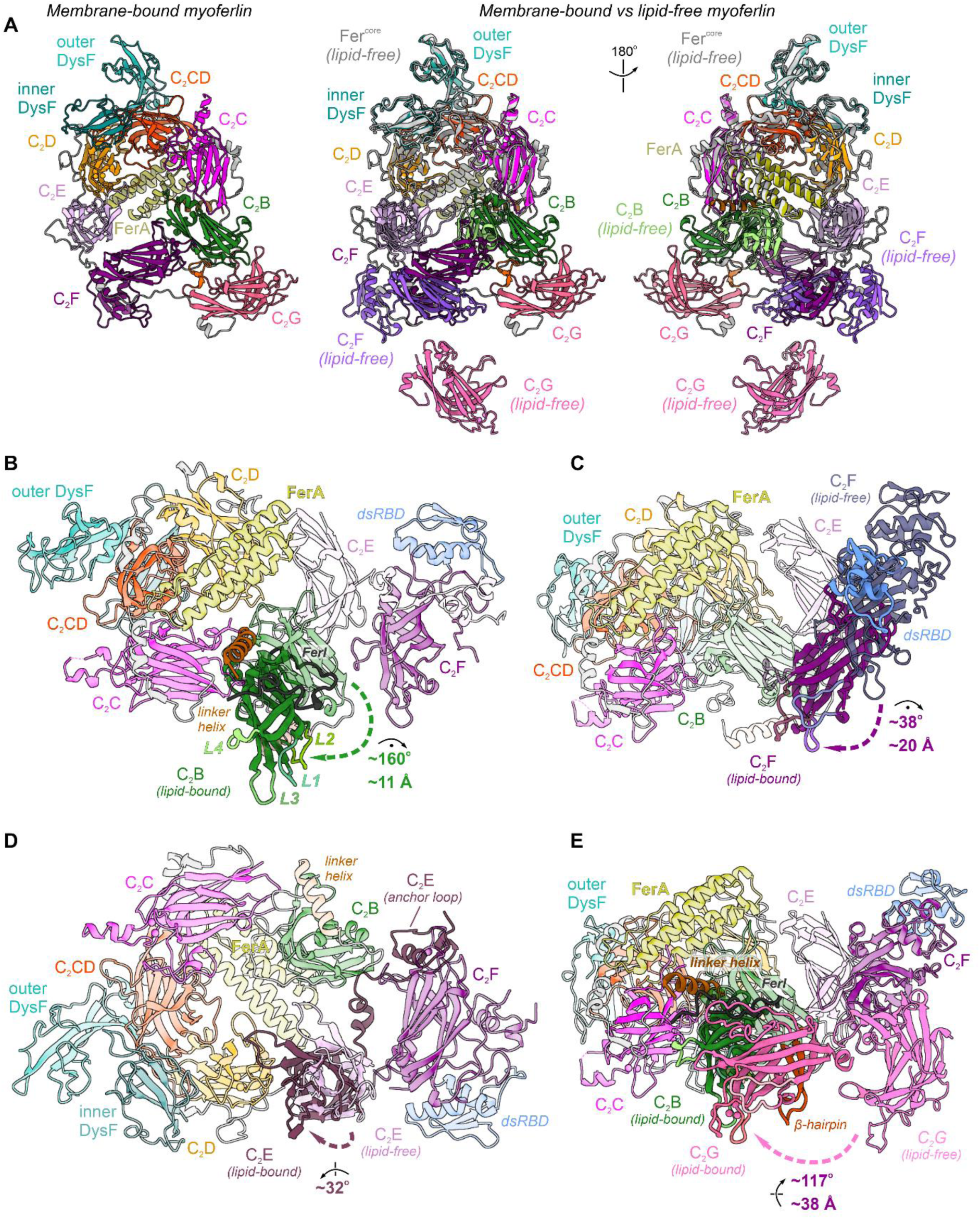
Large-scale conformational transition of ferlins upon lipid membrane binding. **A** Side-by-side comparison of lipid-free and membrane-bound myoferlin. The structures were superimposed based on the Fer^core^ module and the individual domains are colour coded. **B** The large-scale conformational rearrangement of C_2_B upon lipid membrane binding. Myoferlin in the lipid-free state is shown as a transparent model. As a result of a ∼160° out- of-plane rotation and ∼11 Å translation, the top L3 loop of C_2_B projects towards the nanodisc surface in the lipid-bound state. Concomitantly, the linker helix and FerI motif are positioned between C_2_C and C_2_F, respectively, likely stabilizing the new pose of C_2_B. **C** The large-scale displacement of the C_2_F domain upon lipid bilayer binding. Compared to the lipid-free states of myoferlin and dysferlin, where the top loops are oriented towards C_2_B, C_2_F moves by ∼20 Å in the direction of Fer^core^ and towards the membrane plane. **D** In-plane rotation of C_2_E which accompanies the lipid membrane recognition by human myoferlin. Because of the observed rearrangement, the long L4 loop of C_2_E establishes contacts with the Ca^2+^-bound L1 loop of C_2_D, likely fixing the new pose. As the anchor loop wraps around C_2_G and is displaced together with C_2_E, it is possible that C_2_E and C_2_G domain movements are coupled. **E** C_2_G translocates by ∼40 Å to engage the lipid bilayer together with C_2_B and C_2_F in the myoferlin structure. The β-hairpin subdomain of C_2_G is coloured in orange. Because in the lipid-free states of myoferlin and dysferlin C_2_G is dynamic and shares no interfaces with C_2_B and C_2_F, it is likely that the lipid-dependent movement of C_2_B and C_2_F precedes the recruitment of the membrane proximal C_2_ domain. Myoferlin model in the lipid-free state is depicted as transparent cartoons.

Therefore, consistent with our assays (**Appendix Fig S1D-G**), the established membrane contacts of C_2_B, C_2_G and inner DysF have a mixed nature: they are neither exclusively electrostatic nor hydrophobic, nor do they appear to confer strict specificity for acidic phospholipids (such as PS and PI(4,5)P_2_) via Ca^2+^-binding. Such an interaction mechanism suggests that the positioning of C_2_B, C_2_G, and possibly, inner DysF might be modulated by neighbouring C_2_ domains, rather than by bona fide Ca^2+^-driven membrane recruitment.

### Myoferlin’s Ca^2+^-and phospholipid binding sites

Compared to the side-oriented C_2_B and C_2_G, the C_2_C domain establishes a major membrane contact site by engaging the lipid bilayer through its Ca^2+^-coordinating top loops (**Fig 4F**, **Fig 4I and Fig 5A-C**). The local resolution of the myoferlin map enabled accurate modelling of three bound Ca^2+^ ions and a recruited PS lipid headgroup, revealing an intricate membrane recognition mechanism (**Fig 4F and Fig 5A-C**). The largest contact interface between C_2_C and the nanodisc is provided by the long L1. While coordinating two Ca^2+^ ions (Ca1 and Ca2) through D374 and D396 at its base (**Fig 5A-C and Fig EV1B-C**), the distal end of C_2_C’s L1 comprises a conserved helical insertion (residues 377-382, **Fig 4I**) of amphipathic nature that interacts with the membrane mainly with its hydrophobic side (F377, V381, K382). The L3 of C_2_C (**Fig 5A-C**) coordinates all three Ca^2+^ ions (Ca1-Ca3) through D444, W445 (backbone carbonyl), D446, T449, D452, and simultaneously contacts the lipid bilayer (**Fig 4F**) through both hydrophobic and polar residues at its tip (R447, L448, T449, K450). Intriguingly, the modelled PS headgroup (**Fig 5B-C and Fig EV1M**), originating from the lipid nanodisc, binds at the interface between the L3 and the L4 loops of C_2_C. As observed in other C_2_ domain structures (Honigmann et al., 2013, Rizo & Sudhof, 1998, Verdaguer, Corbalan-Garcia et al., 1999), the bound PS headgroup completes the coordination shells of the second and third Ca^2+^ sites (Ca2 and Ca3). At the same time, its serine moiety (**Fig 5B-C**) is recognized by L4 residues (Y513, G515), while L4 itself forms several additional membrane contacts (via F516 and P517).

Like C_2_C, the C_2_F domain binds the nanodisc surface using its Ca^2+^-coordinating loops, while also recruiting the β-hairpin subdomain of C_2_G to the same site (**Fig 4G-H and Fig 5E-F**). In such a configuration, the L3 tip of C_2_F inserts deeply into the lipid bilayer (residues F1628, T1629, R1630), contributing, along with L1, to the coordination of three Ca^2+^ ions (**Fig 5E- F**). The first Ca^2+^ ion (Ca1) is coordinated by typical aspartate residues in L1 (D1575, D1569) and L3 (D1624, Y1625), while the second and third Ca^2+^ (Ca2 and Ca3) involve a glutamate residue in L3 (E1632), a backbone carbonyl (R1630), and a recruited PS headgroup to complete their coordination shells. Interestingly, although not directly binding Ca^2+^ or phospholipids, the L2 of C_2_F forms close contacts with the β-hairpin motif of C_2_G at the same membrane contact site (**Fig 4H**). The β-hairpin subdomain (residues 1860-1879), inserted between β4-β5 strands of C_2_G, is conserved among ferlins and comprises two antiparallel β-strands connected by a loop. Consistent with it representing a lipid-binding accessory motif (**Fig 4H and Fig EV1J**), the β-hairpin loop penetrates the membrane surface and interacts with the bilayer through hydrophobic (F1869, W1870, I1872) and basic residues (K1866, H1868), possibly alongside the L3 of C_2_F and the nearby C_2_G.

In conclusion, membrane recognition by human myoferlin is achieved through both Ca^2+^- dependent and Ca^2+^-independent mechanisms, involving four C_2_ domains (C_2_B, C_2_C, C_2_F, and C_2_G) and two accessory motifs (inner DysF and the β-hairpin of C_2_G). The lipid-interacting groups are arranged asymmetrically to one side of the ferlin ring and form multiple composite interfaces (C_2_B-C_2_G, C_2_F-β-hairpin), highlighting a unique binding mode by this large multi-C_2_ domain protein.

#### Lipid-free structures of human myoferlin and dysferlin

To understand how ferlins are organized prior to their membrane recruitment and to clarify which conformational rearrangements accompany their membrane binding, we obtained additional cryo-EM structures of Ca^2+^-bound soluble myoferlin (residues 1-1997) and of the closely related dysferlin (residues 1-2017) at overall resolutions of ∼3.4 Å and ∼3.5 Å, respectively (**Fig 6A-F, Appendix Fig S6-S8 and Fig EV3**). In the lipid-free state of both ferlins, the C_2_ domains also form a generally similar ring-like structure consisting of only six C_2_ motifs (**Fig 6A-F and Movie EV3**). The rigid Fer^core^ module is well-resolved (**Appendix Fig S9A-C and Appendix Fig S10-S11**) and has a similar organization in both ferlin structures, including the C_2_C, C_2_CD-FerA, and C_2_D domains (**Appendix Fig S8A-G and Fig S9D-G**). Importantly, in both structures (**Fig EV3F and Fig S12A-C**), C_2_C and C_2_D coordinate two divalent cations each, and alanine substitution of seven key aspartates in their top loops completely abolished the Ca^2+^ and liposome-binding activity of the Fer^core^ module (**Appendix Fig S12D-G and Fig S13A-C**). The inner and outer DysF motifs are linked to the Fer^core^ and have a generally similar architecture as observed in the lipid-bound myoferlin structures (**Fig 6C**, **Fig 6F, Fig EV3C-E and Appendix Fig S8**). In contrast, the membrane-interacting C_2_B and C_2_F-C_2_G are more dynamic and adopt different poses in the lipid-free state of ferlins **(Fig 6E-F, Fig EV3A, Appendix Fig S6 and Fig S7G-J**).

Compared to the nanodisc-bound structures (**Fig 1D and Fig EV2 A-F**), the lipid-free cryo-EM models of both dysferlin (**Fig 6C-E and Appendix Fig S7-S9**) and myoferlin (**Fig 6E, Fig EV3A-E**) revealed that the N-terminal C_2_B and the C-terminal C_2_E domains have a virtually antiparallel orientation, with the top loops of C_2_B projecting away from the membrane-facing side of the ferlin ring. The C_2_B and C_2_E domains also establish several small tertiary interfaces with C_2_C and C_2_D of the Fer^core^ module, respectively, as well as with the C_2_F domain (both C_2_B and C_2_E), aligned along the short axis of the ring (**Fig 6F and Appendix Fig S7G-J**). At the same time, like in the lipid-bound state, the anchor loop of C_2_E wraps around C_2_F, ∼40 Å away from C_2_E’s β-sheets (**Fig 6C and 6F**). As a result of these multiple contacts, the C_2_F domain appears surrounded on three sides by C_2_B, C_2_E, and the poorly resolved and dynamic C_2_G domain (**Fig 6D-F, Fig EV3 D-E and Appendix Fig S7G- J**). This conformation of C_2_F significantly differs from the membrane-bound state of myoferlin, where the domain engages the lipid bilayer through its Ca^2+^-binding loops (**Fig 4G and Fig 6F**). Consequently, in the lipid-free states of myoferlin and dysferlin (**Fig 6C-F and Fig EV3 D-E**), while still available for Ca^2+^-binding, the orientation of C_2_F’s L1-L3 loops seems incompatible with efficient membrane insertion, partly due to steric hindrance from the interacting C_2_B domain.

To confirm the physiological relevance of the lipid-free ferlin state, resolved here at lower-resolution for myoferlin and dysferlin, we subjected soluble dysferlin (residues 1-2017) to chemical crosslinking in the presence of Ca^2+^ and identified the crosslinked residues by tandem mass-spectrometry (Cretu, Schmitzova et al., 2016). Mapping the crosslinked lysines onto the cryo-EM structure of dysferlin (**Fig 6G-H, Appendix Fig S14A-C and Dataset EV1**) indicated that 66 out of 71 observed unique crosslinks (∼92.96%) occurred between residues less than 35 Å apart (48/52, ∼92.31%, when omitting the more dynamic C_2_G domain). Importantly, the existence of characteristic interfaces between C_2_B, C_2_E, C_2_F- dsRBD and C_2_G, as observed in the lipid-free dysferlin model (**Fig 6H**), was strongly supported by the crosslinking data, with only five outlier interdomain crosslinks being observed, all of them involving the flexible C_2_G and the anchor loop of C_2_E. Consistent with the dysferlin cryo-EM map (**Fig 6G-H, Appendix Fig S14B and S14D**), our crosslinking data placed the C_2_E domain between C_2_B, C_2_D and the C_2_F-dsRBD module. Furthermore, the location of C_2_F-dsRBD at the base of the dysferlin ring was supported by its crosslinks to both C_2_B and C_2_E, and to the neighbouring C_2_G, flanking C_2_F from both sides (**Fig 6H and Appendix Fig S14B**). From these analyses, we conclude that the lipid-free conformation is indeed populated in solution, possibly representing the default state of the ferlins in cells.

#### The different conformational states of human ferlins

Despite being comparable in general terms, the lipid-free and membrane-bound ferlin structures represent two distinct conformations (**Fig 7A and Movie EV4**). Structure-based superposition and modelling of the transition between the two resolved states of myoferlin allowed us to decipher the possible sequence of events (**Fig 7B-E, Movies EV4 and EV5**). We delineated major structural rearrangements occurring at several N-terminal (C_2_B) and C-terminal (C_2_E-C_2_G) sites of the myoferlin ring, indicating a multistep transition (**Fig 7B-E and Movie EV5**). Compared to the rigid Fer^core^, our analyses show that the N-terminal C_2_B appears to rotate by ∼160° upon membrane binding and moves by ∼11 Å towards the membrane plane (**Fig 7B-E and Movie EV5**). This dramatic change in the domain’s orientation entails both disruption (C_2_E), reconfiguration (C_2_C and C_2_F), and formation (with C_2_G and the linker helix) of tertiary interfaces and membrane contacts (through C_2_B’s L3). Concomitantly, our superposition indicates that the C_2_F-dsRBD module, which, in the lipid-free state, interacts with C_2_B, appears to translocate by ∼20 Å and rotate by an additional ∼38° from the ring’s periphery towards the Fer^core^ (**Fig 7C**). The large-scale displacement of the C_2_F domain would enable the previously hindered Ca^2+^-binding loops to directly engage the lipid bilayer (**Fig 7C**). Accompanying this C_2_F repositioning (**Fig 7D and Movie EV5**), the upstream C_2_E, interacting with C_2_F through the anchor loop, also rotates by ∼32° in the direction of the Fer^core^, while its long L4 contacts the Ca^2+^-bound L3 of C_2_D. Notably, the new poses of C_2_E and C_2_F appear to be stabilized initially by the repositioned C_2_B (**Fig 7C- D**) and the unstructured FerI motif, which covers the distance between C_2_B and C_2_C. Finally, the C_2_G domain, flexible in the lipid-free states, binds the reconfigured C_2_B-C_2_F interface at both of its sides to close the myoferlin ring. Besides establishing new interdomains contacts, the rearranged C_2_G simultaneously engages the lipid membrane with its Ca^2+^-binding loops and its β-harpin subdomain, inserting into the bilayer in the vicinity of C_2_B and C_2_F, respectively (**Fig 7E**). Interestingly, our structures also show that C_2_G binds more tightly phosphoinositide-containing bilayers, possibly because of additional interactions between its polybasic patch and exposed PI(4,5)P_2_ headgroups, as previously suggested and/or observed in other C_2_ domain structures (Carpenter, Khuu et al., 2023, Guerrero-Valero, Ferrer-Orta et al., 2009, Guillen, Ferrer-Orta et al., 2013, Kwok et al., 2023, Padmanarayana et al., 2014) (**Fig EV4 A-C**).

The conformational rearrangement of myoferlin (and possibly dysferlin) to its lipid-bound state is possible because the C_2_ domains undergoing displacement share relatively small contact interfaces in the lipid-free conformation, and the connecting linker regions are long and partially unstructured (**Fig 1A, Fig EV2 B-C, Appendix Fig S13D**). For example, C_2_E’s contact interfaces with C_2_B (∼252 Å^2^), C_2_D (∼347 Å^2^) and C_2_F (∼235 Å^2^, excluding the anchor loop) are equally small. Likewise, in the lipid-free state, the C_2_B-C_2_F interface involves a relatively small number of residues (∼193 Å^2^) and is not present in all the imaged particles (**Fig 6D-E, Fig EV3A and Appendix S6C**). Moreover, calpain-cleavage of myoferlin and alternative dysferlin isoforms appears to release of the C-terminal C_2_F-C_2_G domains, which indicates that the C_2_B-C_2_F and C_2_E-C_2_F interfaces are also dynamic in cells (Piper, Ross et al., 2017, Redpath, Woolger et al., 2014). In this respect, the accessory ferlin structural motifs – the β-hairpin of C_2_G, the linker helix and FerI – resolved exclusively in the lipid-bound state of myoferlin, are likely required to stabilize the reconfigured C_2_ domain interfaces.

Therefore, the tertiary interactions between the Fer^core^, C_2_B, C_2_E and C_2_F, observed in the lipid-free structures, seem to be generally weak, having possibly evolved to facilitate efficient membrane sampling on fast timescales and in a complex cellular environment. Formation of new and extended contacts between C_2_B, C_2_F, and the repositioned C_2_G not only increases the structure’s stability, but also allows their concomitant, in-plane binding as part of a composite and asymmetric interface. Because the C-terminal C_2_G is connected to the transmembrane helix through a relatively short linker (approximately 20 residues), our structures indicate that the large-scale movement of the domain towards the Fer^core^ would shorten the distance between the ferlin vesicle and the target membrane, likely facilitating their close apposition. Indeed, supporting such a scenario, we observe that both myoferlin and dysferlin engage lipid bilayers in a Ca^2+^-dependent manner and promote tight vesicle-vesicle interaction (*i.e.*, docking) *in vitro,* when both PS and PI(4,5)P_2_ are present on the target membrane (**Fig EV4 D-E)**. Given the high degree of conservation of their lipid-free structures, it is likely that dysferlin and, possibly, otoferlin transition through a similar sequence of domain rearrangements and conformational states upon membrane binding (**Fig 8**). Such a role of ferlins in progressing from loose vesicle tethering to tight docking of membranes has been suggested for otoferlin based on electron tomography (Vogl, Cooper et al., 2015) and, like myoferlin (**Fig EV4 D-E**) and dysferlin (Codding, Marty et al., 2016), otoferlin’s C_2_ domains accelerate SNARE-dependent membrane fusion *in vitro* (Johnson & Chapman, 2010). These functional similarities indicate that ferlins may indeed share common organization principles and a conserved membrane remodelling mechanism.

**Figure 8.**
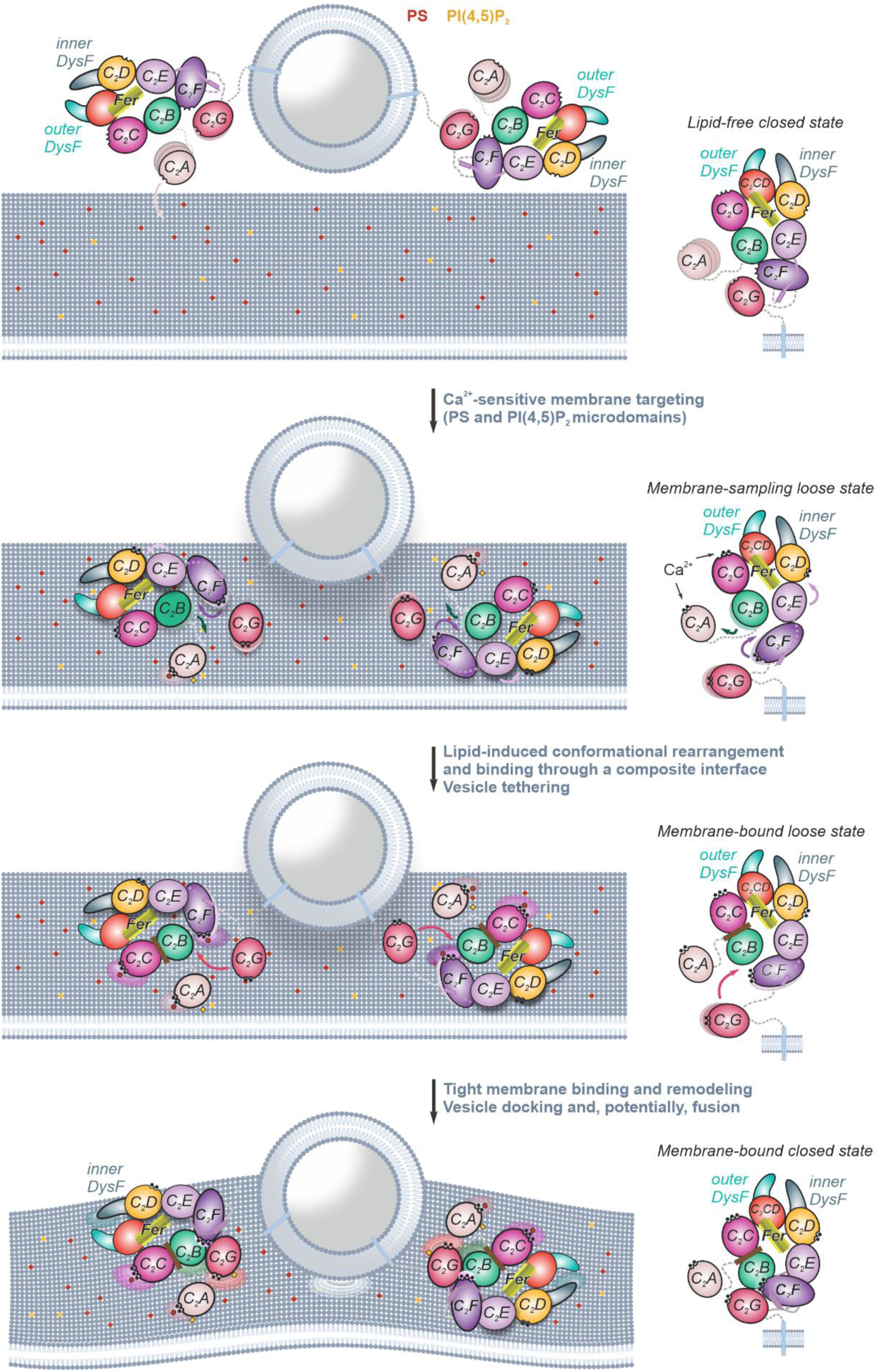
Model of ferlins’ Ca^2+^-sensitive recruitment and binding to lipid membranes. Schematic depicting a structure-guided model for how ferlins facilitate Ca^2+^-dependent vesicle targeting, docking, and local membrane remodelling to modulate and possibly promote fusion of two lipid bilayers. The ferlin domains are color-coded and those interacting with the lipid membranes are outlined in black. PS (phosphatidylserine) and PI(4,5)P_2_ (phosphatidylinositol-4,5-bisphosphate) are indicated as red and orange, respectively. The membrane-sampling loose state has been observed in the lipid-free dysferlin cryo-EM data (**Appendix Fig S6A and Fig S14A**). The hypothetical membrane-bound loose state, where the C-terminal C_2_G domain is yet to engage the lipid bilayer, is based on Alphafold2 predictions of myoferlin and dysferlin (**Appendix Fig S15B-D**). The lipid-free and membrane-bound closed states have been depicted based on the cryo-EM structures of lipid- free myoferlin/dysferlin and membrane-bound myoferlin, respectively.

## Discussion

Ferlins are the largest and the most complex multi-C_2_ domain Ca^2+^-sensitive factors (Lek et al., 2012). In this study, we report the near-atomic cryo-EM structures of human myoferlin in its Ca^2+^ and lipid-bound state. We also obtained cryo-EM models of both myoferlin and the related dysferlin in their lipid-free states and carried out supporting experiments to validate our structural models. These analyses provide first insights into how the unique structural features of ferlins are adapted to their many ascribed roles in cellular trafficking pathways, underlying the acute response to membrane and organelles injuries, muscle biogenesis and metabolism, or fast transmitter exocytosis at hair cell ribbon synapses.

We propose that the multifaceted roles of ferlins in diverse Ca^2+^-dependent pathways in cells could stem from their ability to sample alternative conformational states (**Fig. 8**), achieved by exploiting a unique structural organization. Contrary to previous models (Dominguez et al., 2022, Woolger et al., 2017, Xu et al., 2011), but in agreement with Alphafold2 predictions, nanoscopic imaging (Shaib, Chouaib et al., 2024) and a recent lipid-free dysferlin structure (Huang et al., 2024), the ferlin C_2_ domains are not trivially organized as “beads on a string” or as in other, well-studied multi-C_2_ domain factors (Schauder et al., 2014, Shin, Han et al., 2005). Instead, the conserved structural motifs pack uniquely in 3D and form state-defining interfaces, bridging both neighbouring and sequence-distant domains. The central ferlin core module (Fer^core^) comprises the region spanning the C_2_C and C_2_D domains. Fer^core^, likely, maintains part of the, otherwise dynamic, ferlin structure rigid and might support the ordered and reversible Ca^2+^/lipid-driven conformational transitions of ferlins. In this respect, the newly identified ferlin C_2_-like domain, the C_2_CD domain, observed in our myoferlin and dysferlin structures, and the closely linked four-helix bundle of FerA might have evolved to promote the tight packing of the Fer^core^ module. Importantly, Alphafold predictions support the presence of a similarly organized Fer^core^ in all remaining paralogs, including otoferlin, whose mutations cause the nonsyndromic deafness DFNB9 and are a target of the first inner ear gene-therapy trials (Lv, Wang et al., 2024, Moser et al., 2024) (**Appendix Fig S15A and S15D**).

Although appearing defined and stable enough to be observed upon cryo-EM imaging, the interfaces between the membrane-proximal domains and the Fer^core^ module are more dynamic in the lipid-free states of myoferlin and dysferlin. This notable structural feature appears to be a direct consequence of the highly specialized roles served by ferlins in cells. Having low energy barriers between their discrete conformations could, possibly, explain the transiently formed contacts between state-defining C_2_ domains, such as C_2_B, C_2_F and C_2_G (**Fig. 8**). This may constitute a significant advantage in protein-rich microenvironments, such as the active zones of hair cell ribbon synapses (otoferlin) (Moser, Grabner et al., 2020) or caveolae-rich plasma membrane compartments (dysferlin) (Corrotte, Almeida et al., 2013), allowing them to sample conformations close to the ground state. As a result, switching between the different ferlin conformations could be accomplished by reversibly shifting the equilibrium towards a given, functionally relevant state without a significant free energy consumption (**Fig. 8**). Consistently, all ferlin-dependent cellular processes occur on fast timescales, depend on Ca^2+^, and are highly dynamic and reversible, with the ferlins appearing to be needed for both the forward and reverse reactions (*e.g.*, synaptic vesicle exocytosis and replenishment, vesicle docking and undocking).

An important question raised by our structural analyses pertains to the possible connections between the observed structural changes upon membrane binding *in vitro* (**Fig. 8**) and the documented roles of ferlins in promoting Ca^2+^-sensitive vesicle tethering and membrane fusion. As specialized trafficking factors highly expressed in muscle cells, the membrane-anchored myoferlin and dysferlin reside at the sarcolemma and shuttle to the endosomal compartment, without accumulating in a defined intracellular vesicle pool (Cooper & McNeil, 2015, Lek et al., 2012). The related otoferlin is enriched at the presynaptic active zone membranes and in the synaptic vesicles of inner hair cells, while also shuttling through the endosomal pathway during a synaptic vesicle release cycle (Jung et al., 2015, Moser et al., 2020, Pangrsic et al., 2012, Revelo, Kamin et al., 2014). In all cases, the ability of ferlins to react to an increase in intracellular Ca^2+^, following large sarcolemma injuries (dysferlin) or a sound-evoked receptor potential (otoferlin), appears to be determined by their cellular localization on vesicles, interactions with other factors, and the local phospholipid composition of the target membranes (Chen, Fang et al., 2023, Chen et al., 2024). Consistent with having such multimodal mechanisms of action, the cryo-EM models of myoferlin and dysferlin revealed a dynamic structural organization, apparently evolved to accommodate different membrane environments and binding modalities. Our structures show that the Ca^2+^- sensitive N-terminal C_2_A (in type-I ferlins) and the membrane proximal domains (C_2_F-C_2_G) are more dynamic in the lipid-free ferlin states, which could result in an increased membrane capture radius (**Fig. 8**). We, therefore, propose that by employing their mobile C_2_ domains as pioneer Ca^2+^ and lipid-sensitive motifs, ferlins would be able to efficiently sample and approach distant target membranes. Consequently, membrane attachment of these pioneer C_2_ domains would trigger, in the next step, the tight association of the remaining Fer^core^ core domains (C_2_C-C_2_D and the inner DysF), altogether assembling an asymmetric ring-like structure that could bridge two cellular membranes (tethering) and, at the same time, promote local membrane buckling and remodeling through deep insertion into the bilayer at multiple contact sites (tight vesicle docking, **Fig. 8**). In our model, as only one side of the ring would engage the lipid bilayer, the available ferlin domains, such as FerA, could simultaneously interact with other pathway factors, such as SNAREs (Codding et al., 2016) or other ferlin molecules (Pangrsic et al., 2012), further narrowing the gap between the two membranes to facilitate their fusion (**Fig. 8**). This structural adaptation could be particularly relevant for dysferlin, whose functions in plasma membrane repair have been linked to the Ca^2+^-sensitive clustering of highly heterogenous endomembrane vesicles to promote their homotypic and heterotypic fusion (Cooper & McNeil, 2015).

Future functional and structural studies are needed to probe the validity of these mechanisms, beyond simple nanodisc membranes, particularly within the context of pathway-defining, higher-order ferlin complexes, as pioneered for synaptotagmins (Zhou, Lai et al., 2015, Zhou, Zhou et al., 2017). These studies will likely entail the use of alternative imaging modalities (cryo-electron tomography and super-resolution imaging) and computational approaches (molecular dynamics simulations), applied to more complex membrane systems (such as liposomes or lipid nanotubes). Such analyses will help exclude improbable non-physiological myoferlin conformations induced by nanodiscs and further elucidate the roles of additional ferlin molecules and ferlin-SNAREs interactions in the vesicle-membrane fusion cycle.

## Materials and Methods

### Reagents and tools table

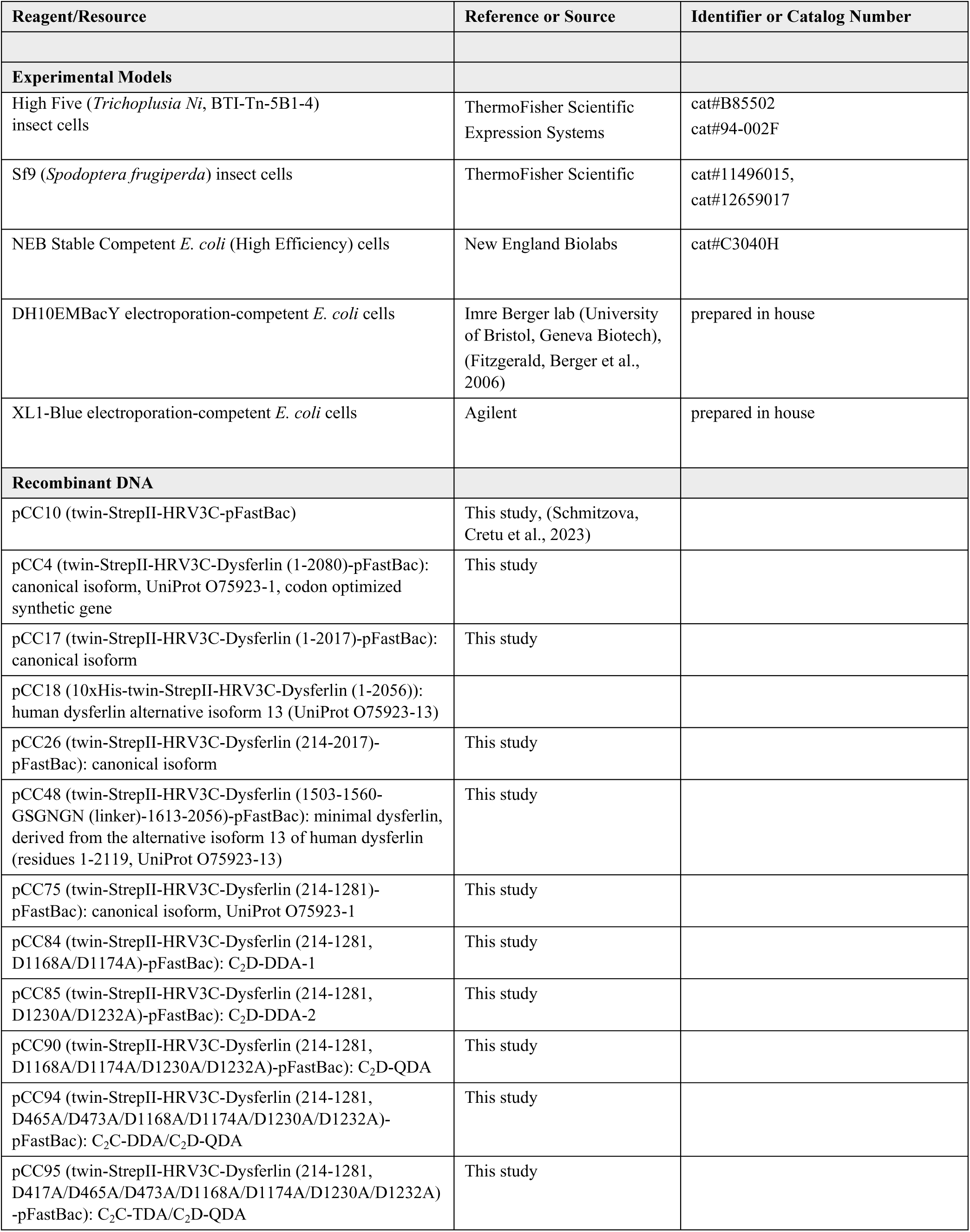
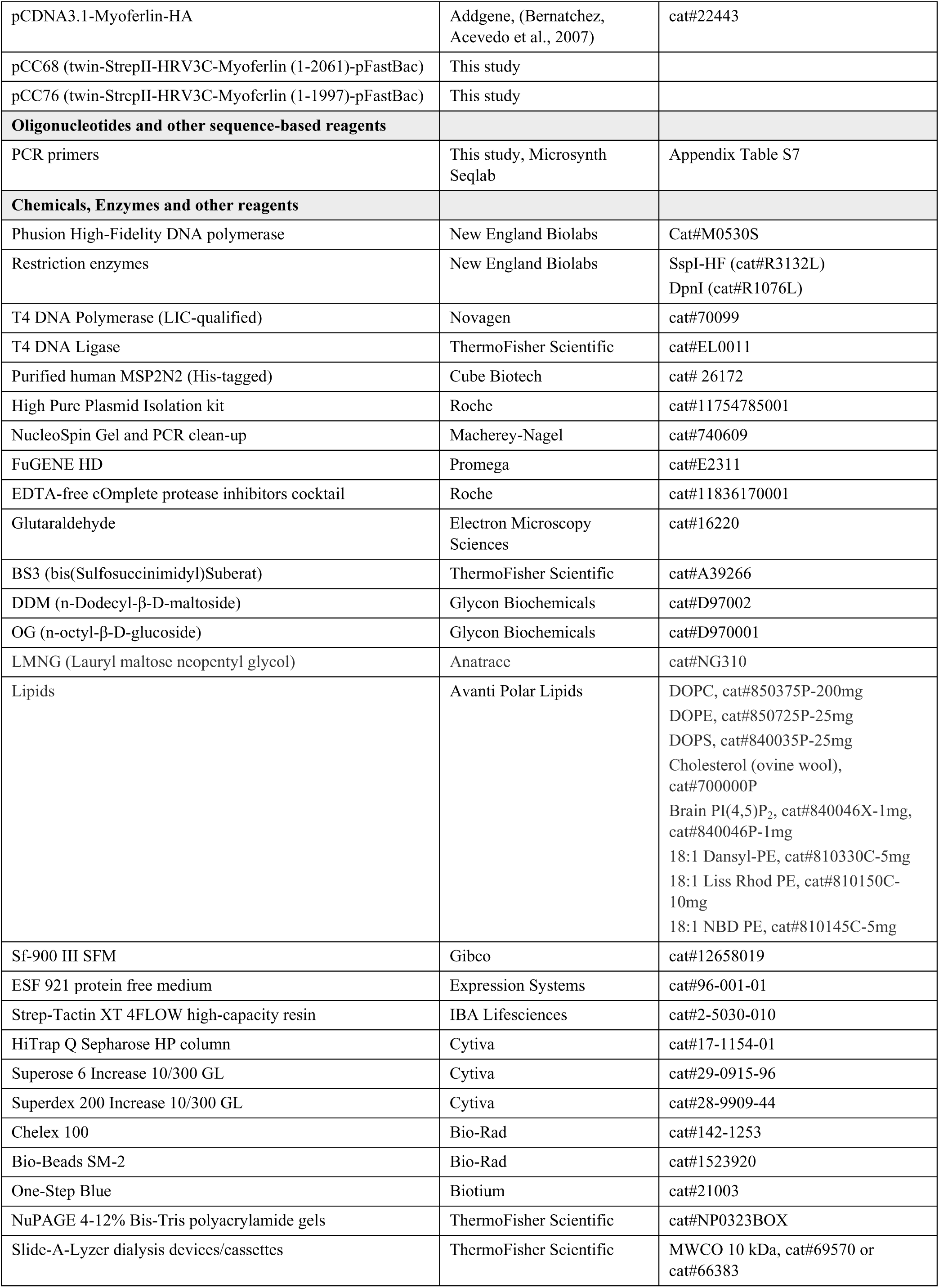
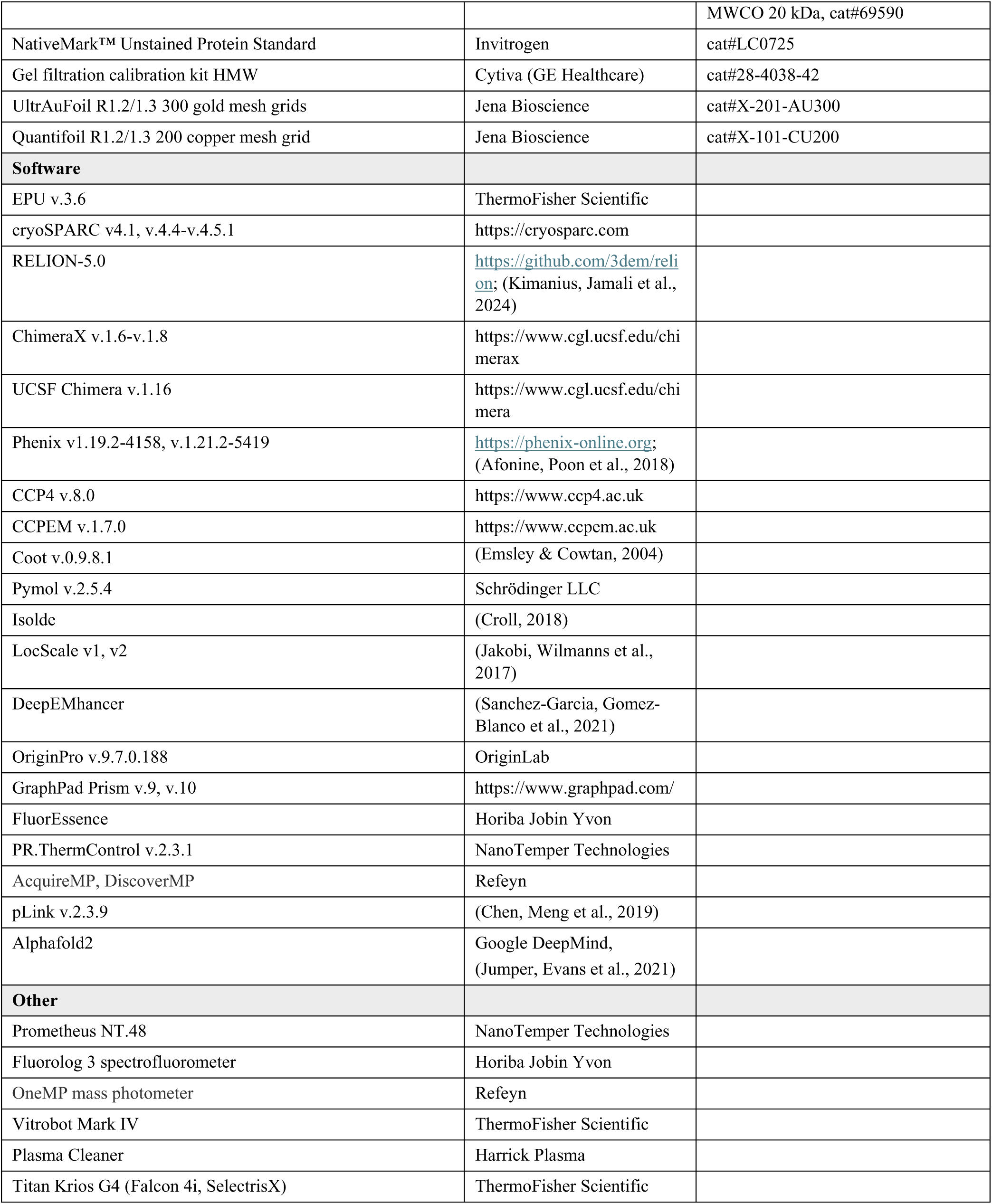

## Methods and protocols

### Plasmids and molecular cloning

The codon-optimized full-length canonical human dysferlin (DYSF, FER1L1) expression construct (NM_003494.4, transcript variant 8, isoform 1, UniProt O75923-1) was constructed by GenScript and cloned into the pFastBac1 vector backbone by using the EcoRI and KpnI restriction sites in frame with an N-terminal twin-StrepII affinity tag, which can be cleaved with the HRV-3C (Human Rhinovirus 3C) protease. The full-length human myoferlin (MYOF, FER1L2, Uniprot Q9NZM1-1) was obtained from Addgene (pCDNA3.1-Myoferlin- HA, cat#22443, (Bernatchez et al., 2007)) and subcloned by LIC (ligation independent cloning) into a modified pFastBac backbone (first described in (Schmitzova et al., 2023)), in frame with an HRV-3C protease cleavable N-terminal twin-StrepII-tag. The soluble dysferlin (residues 1-2017) and myoferlin (residues 1-1997) constructs and the domain truncation mutants of dysferlin (Fer^core^ (the C_2_B-C_2_D fragment, residues 214-1281) and minimal dysferlin (residues 1503-1560 and 1613-2056), encompassing the C_2_F-C_2_G domains) were obtained by “around-the-horn” PCR-based cloning using the full-length constructs as templates and retained the N-terminal twin-StrepII tag. Neutralizing substitutions in the Ca^2+^- binding motifs of the dysferlin core region (residues 214-1281) were introduced by sequential PCR-based site-directed mutagenesis (**Appendix Table S7**). All constructs were verified by Sanger sequencing (Microsynth Seqlab GmbH, Göttingen) of the open reading frames (ORFs). All cloning primers were synthesized by Microsynth Seqlab GmbH (Göttingen).

### Expression and purification of human myoferlin and dysferlin

To enable their expression in insect cells, all myoferlin and expression constructs were incorporated into bacmids through transformation of electro-competent DH10MultibacY cells, as previously described (Cretu, Agrawal et al., 2018, Cretu, Gee et al., 2021, Cretu et al., 2016, Schmitzova et al., 2023), extracted for transfection using the High Pure Plasmid Isolation kit (Roche, cat#11754785001), and precipitated with isopropanol. To produce V_0_ recombinant baculoviruses, adherent Sf9 (*Spodoptera frugiperda*) cells, cultured in Sf-900 III SFM (Gibco, cat#12658019), were transfected with the prepared bacmids with FuGENE HD (Promega, cat#E2311) in 6-well plates. The initial V_o_ viruses were amplified by infecting Sf9 cells at a 1:10-1:20 ratio to produce the V_1_ generation of baculoviruses. For large-scale protein production, the V1 baculoviruses were used to infect Sf9 or Hi5 (*Trichoplusia Ni*) suspension cultures, grown in the ESF 921 protein free medium (Expression Systems, cat#96- 001-01), in titers sufficient to induce cell proliferation arrest after 24 hours (Cretu et al., 2016, Schmitzova et al., 2023). Typically, Hi5 and Sf9 cells were infected at a density of ∼1.0x10^6^ cells/mL and harvested 60-72 hours after baculovirus infection, when cells’ viability dropped to ∼80-85%. Insect cell infection was monitored every 16-24 hours by observing the expression of the eYFP marker. Full-length and soluble dysferlin (1-2017) were overexpressed on a preparative scale in Hi5 cells. All myoferlin samples (full-length myoferlin (1-2061) and soluble myoferlin (1-1997)), the Fer^core^ of dysferlin and its mutant variants were expressed and purified from Sf9 cells.

All dysferlin and myoferlin constructs were purified by exploiting the highly specific twin-StrepII affinity tag and their binding to an anion-exchange resin (**Fig S1A**). Membrane-anchored full-length myoferlin and dysferlin were purified in the presence of DDM (n-Dodecyl-β-D-maltoside, Glycon Biochemicals, cat#D97002), whereas, after cell lysis, all soluble ferlin constructs were purified in the absence of detergent. Typically, insect cells (Sf9 or Hi5) from ∼1 L culture were resuspended in 10-15 mL of the Lysis buffer (50 mM HEPES-KOH, pH 7.5, 300 mM KCl, 10% (v/v) Glycerol, 4 mM DTT (Dithiothreitol), 2-2.5% (w/v) DDM (or 0.2% (v/v) Triton X-100 in the case of the soluble constructs), EDTA-free cOmplete protease inhibitors cocktail (Roche, cat#11836170001, 1 tablet per 50 mL buffer)) per gram of cell pellet and detergent-extracted by incubation for 90 min at 4°C while gently rotating on a roller mixer or lysed by sonication on ice (soluble constructs), using the Branson Ultrasonics Sonifier 250 (Duty cycle 30%, Output 3, for 2 min). The crude lysate was cleared by ultracentrifugation at 46,300 r.p.m. for 1 h at 4°C in a type 70Ti rotor or at 15,000 r.p.m. for 1 eh at 4°C in the JA-18 rotor (Beckman Coulter) and further filtered through a 0.8 μm Minisart membrane (Sartorius, cat#16592). The filtered lysate was incubated with 2-3 mL Strep-Tactin XT 4FLOW high-capacity resin (4-6 mL 50% slurry, IBA Lifesciences, cat#2- 5030-010) per liter of culture, for 1 h at 4-8°C on a roller mixer. The resin was pelleted by centrifugation at 2000 r.p.m., transferred to an Econo-Pac (Bio-Rad, cat#7321010) gravity-flow column, and thoroughly washed with the Wash buffer (50 mM HEPES-KOH, pH 7.5, 300-500 mM KCl, 5% (v/v) Glycerol, 2 mM DTT, 0.5% (w/v) DDM (membrane-anchored constructs)) and then with the Binding buffer (50 mM HEPES-KOH, pH 7.5, 150 mM KCl, 5% (v/v) Glycerol, 2 mM DTT, 0.1% (w/v) DDM (membrane-anchored constructs)). The bound proteins were eluted by competition with biotin added to the Elution buffer (50 mM HEPES-KOH, pH 7.5, 120-140 mM KCl, 5% (v/v) Glycerol, 2 mM DTT, 0.1% (w/v) DDM (in case of the membrane-anchored constructs), 1 mM EDTA (Ethylenediaminetetraacetic acid), 60 mM biotin). The affinity tag was not cleaved.

To remove nucleic acid contaminants and minor impurities, the Strep-Tactin eluates were further purified by anion-exchange chromatography on a 5 mL HiTrap Q Sepharose HP column (Cytiva, cat#17-1154-01), mounted on an Äkta go system. Prior to sample application, the Q Sepharose HP column was equilibrated with Buffer A (20 mM HEPES-KOH, pH 7.5, 150 mM KCl, 5% (v/v) Glycerol, 2 mM DTT (or 1 mM TCEP (Tris(2- carboxyethyl)phosphine)), 0.03% (w/v) DDM (membrane-anchored constructs)). In the next step, the column was washed extensively with Buffer A and the sample eluted off the column using a linear, 0-30% gradient formed between Buffer A and Buffer B (20 mM HEPES-KOH, pH 7.5, 1 M KCl, 5% (v/v) Glycerol, 2 mM DTT (or 1 mM TCEP), 0.03% (w/v) DDM (membrane-anchored constructs)) over 60-80 mL. The peak ferlin fractions were pooled and concentrated by ultrafiltration to ∼3-3.5 mg/mL (membrane-anchored constructs), ∼4.0-5.5 mg/mL (soluble dysferlin (1-2017) and myoferlin (1-1997)) or ∼7.5-10 mg/mL (the domain truncation mutants of dysferlin), aliquoted, snap frozen in liquid nitrogen, and stored at - 80°C. The identities of the purified proteins were verified by mass-spectrometry (Proteomics Facility, Max-Planck-Institute for Multidisciplinary Sciences, Göttingen).

### NanoDSF-based characterization of myoferlin and dysferlin

As means of protein quality control, all myoferlin and dysferlin preparations (both soluble and membrane-anchored) were subjected to nanoDSF (nano differential scanning fluorimetry) measurements. In a nanoDSF typical assay, the 5 µL protein sample (∼0.6 µM) was mixed with 5 µL assay buffer (25 mM HEPES-KOH, pH 7.5, 150 mM KCl, 0.03% (w/v) DDM (membrane-anchored constructs)), which was pretreated with Chelex 100 (Bio-Rad, cat#142-1253) and supplemented with increasing concentrations of CaCl_2_ or MgCl_2_ (0-40 mM). Following 10 min incubation at room temperature, the samples were loaded into capillaries and the emission intensity at 350 nm and 330 nm (due to the intrinsic fluorescence of tryptophan residues at 280 nm) was measured as a function of temperature using a Prometheus NT.48 instrument (NanoTemper Technologies). The temperature was increased from 20°C to 95°C by applying an unfolding ramp of 1°C/min and the excitation power was adjusted to yield at least 2000 integrated fluorescence counts at both wavelengths. The melting temperatures (*T*_m_) were estimated by plotting the first derivative of the 350 nm/330 nm ratio as a function of temperature in the PR.ThermControl v2.3.1 software. The [Me^2+^]_1/2_ values were estimated in GraphPad Prism 10 (v10.3.1) by nonlinear regression fitting to a modified Hill function: *T*_m_([Me^2+^])=*T*_m_i+(*T*_m_f-*T*_m_i)x([Me^2+^]*^n^*/([Me^2+^]_1/2_*^n^*+[Me^2+^]*^n^*)), where *T*_m_i and *T*_m_f, represent the initial and final *T*_m_, respectively, and *n* – the Hill coefficient.

### Preparation of liposomes

All lipids (phospholipids and cholesterol), used to prepare liposomes (Large unilamellar vesicles) and lipid nanodiscs, were obtained from Avanti Polar Lipids as powder (or chloroform stocks), further dissolved in chloroform to their working concentrations, and stored at -20°C. Porcine brain PI(4,5)P_2_ (L-α-phosphatidylinositol-4,5-biphosphate) was dissolved in a chloroform:methanol:water (20:9:1) solution. Unlabeled LUVs devoid of anionic phospholipids (referred to as “DOPC-DOPE-only LUVs”) were prepared by mixing DOPC (1,2-dioleoyl-sn-glycero-3-phosphocholine, 18:1 (Δ9-cis) PC), DOPE (1,2-dioleoyl-sn-glycero-3-phosphoethanolamine, 18:1 (Δ9-cis) PE) and Cholesterol (ovine wool) in a 7:2:1 molar ratio (70:20:10 mol%) to a final 8 mM concentration. LUVs comprising in addition DOPS (1,2-dioleoyl-sn-glycero-3-phospho-L-serine, referred to as “15 mol% DOPS LUVs”) and both DOPS and PI(4,5)P_2_ (referred to as, for example, “25 mol% DOPS/5 mol% PI(4,5)P_2_ LUVs”) were prepared by substituting a part of DOPC in the lipid mixture with respective anionic phospholipids to obtain the desired lipids’ ratio. Dansyl-labeled LUVs were prepared by replacing a portion of DOPE in the lipid mixture with 5 mol% 18:1 Dansyl PE (1,2-dioleoyl-sn-glycero-3-phsophoethanolamine-N-(5-dimethylamino-1- naphtalenesulphonyl)). Similarly, Rhodamine B and NBD (Nitro-2,1,3-benzoxadiazole-4-yl)- labelled LUVs were obtained by replacing a part of DOPE in the lipid mixture with 1 mol% 18:1 Lissamine Rhodamine B PE (1,2-dioleoyl-sn-glycero-3-phosphoethanolamine-N- (lissamine rhodamine B sulfonyl)) and 1 mol% 18:1 NBD PE (1,2-dioleoyl-sn-glycero-3- phosphoethanolamine-N-(7-nitro-2-1,3-benzoxadiazol-4-yl). To obtain unilamellar liposomes, the mixed lipids were transferred to a glass vial, the solvent was evaporated under a nitrogen stream and the lipid film was then dried in a vacuum desiccator (∼200 mbar) for at least 3 hours to overnight. The lipid film was hydrated in reconstitution buffer (20 mM HEPES-KOH, pH 7.5, 150 mM KCl) by gently vortexing. The resulting multilamellar vesicles were extruded through a 0.4 µm Nuclepore track-etch membrane (Cytiva, cat#10417104) for at least 21x using a Mini-extruder (Avanti Polar Lipids) and further used in coflotation and Dansyl-based lipid binding assays. Vesicles used for proteoliposome reconstitution and in lipid mixing assays were additionally passed 21x through a 0.1 µm Nuclepore membrane (Cytiva, cat#10419504). For the reconstitution of MSP2N2-based lipid nanodiscs, lipid films were prepared as for liposome reconstitution, except omitting DOPE and cholesterol from the lipid mixture (in the case of the 25 mol% DOPS/5 mol% PI(4,5)P_2_ and 15 mol% DOPS/2 mol% PI(4,5)P_2_ nanodiscs) or using lower ratios of DOPE (10 mol% DOPE and 5 mol% 18:1 Dansyl PE) and Cholesterol (5 mol%) when assembling the 25 mol% DOPS/5 mol% PI(4,5)P_2_/5 mol% Cholesterol and 15 mol% DOPS/5 mol% Cholesterol nanodiscs. For nanodisc reconstitution, the dried lipid films were hydrated in the reconstitution buffer supplemented with 1.7 % (w/v) or 0.6 % (w/v) DDM, resulting in 13.2 mM or 5.6 mM final lipid concentrations, respectively, and briefly sonicated with the microtip (Branson Sonifier 250).

### Reconstitution of ferlins into proteoliposomes

Dysferlin and myoferlin (**Fig EV4D-E**) were reconstituted into 100 nm LUVs by mixing DOPC/DOPE-only liposomes, OG (n-octyl-β-D-glucoside, Glycon Biochemicals, cat#D970001) and the purified ferlin (in DDM micelles) at a 1:3500 protein-to-lipids ratio, an R-value of 1, and a 4 mM final lipid concentration. The samples were then incubated for 20 min at room temperature and then transferred to Slide-A-Lyzer MINI dialysis devices (with a MWCO 10 kDa, Thermo Fisher Scientific, cat#69570) or Slide-A-Lyzer cassettes (MWCO 10 kDa, Thermo Fisher Scientific, cat#66383) for overnight dialysis at 4-8°C against 2 L of reconstitution buffer (20 mM HEPES-KOH, pH 7.5, 150 mM KCl), supplemented with 2.5 g/L Bio-Beads SM-2 (Bio-Rad, cat#1523920). Bio-Beads SM-2 were prepared fresh by sequential washing with methanol, ethanol, ddH_2_O, and reconstitution buffer. The successful reconstitution of ferlins into liposomes was verified by liposome flotation on a Nycodenz step gradient following ultracentrifugation for 90 min at 50,000 r.p.m. in a TLS-55 rotor (Beckman Coulter).

### Liposome binding assays

To assess the ability of soluble myoferlin (1-1997) to interact with model lipid bilayers, 50 µM Dansyl-labeled LUVs (labeled with 5 mol% 18:1 Dansyl PE), of a varying anionic phospholipid composition, were mixed with ∼0.9 µM ferlin sample in the presence of Ca^2+^. The total reaction volume was 15 µL. Following incubation for 5 min at room temperature, the protein-lipid samples were transferred into an ultra-micro cuvette (QS 105.252, 1.5 mm optical path, Hellma, cat#105-252-15-40) and the emission spectra were taken between 450- 560 nm at a 284 nm excitation wavelength (3 nm slits, 0.1 s integration time) using a Fluorolog 3 spectrofluorometer (Horiba Jobin Yvon). The relative protein-to-membrane FRET (Förster Resonance Energy Transfer) efficiency was calculated as follows: rFRET=(*I*- *I*_min_)/(*I*_max_-*I*_min_), where *I* represents the average emission intensity of the sample at 518-520 nm, *I*_min_ – the intensity of the protein-free blank sample, and *I*_max_ – the maximum Dansyl emission of the titration series. To estimate the [Ca^2+^]_1/2_ values, the liposome binding data were fitted to the Hill equation (Brandt, Coffman et al., 2012) in GraphPad Prism 10. All experiments were repeated three times (technical replicates).

### Coflotation assays

In typical liposome coflotation experiment, soluble ferlin constructs were mixed with LUVs in presence of 50 µM or 0.5 mM (**Appendix Fig S12F**) Ca^2+^/Mg^2+^ and added to the bottom layer of a Nycodenz step gradient. The final protein and liposomes assay concentrations were ∼1 µM and 1 mM, respectively. The bottom Nycodenz layer (40% (w/v)) was then overlaid with equal volumes (40 µL) of a 30% (w/v) Nycodenz solution and reconstitution buffer (20 mM HEPES-KOH, pH 7.5, 150 mM KCl), which were supplemented with 50 µM CaCl_2_ or MgCl_2_ (or 0.5 mM CaCl_2_/MgCl_2_). The Nycodenz gradients were centrifuged at 50,000 r.p.m. for 90 min in a TLS-55 rotor (Beckman Coulter), harvested from the top in 20 µL fractions and analyzed by SDS-PAGE (NuPAGE 4-12% Bis-Tris gels, Thermo Fisher Scientific, cat#NP0323BOX). The SDS-PAGE gels were stained with One-Step Blue (Biotium, cat#21003). Under these experimental conditions, liposomes float to the top two Nycodenz fractions and comigration of ferlins to the top fractions is indicative of their stable interaction with the lipid bilayer. All coflotation experiments were conducted at least three times (technical and biological replicates).

### Lipid mixing assays

In a typical “bulk” lipid mixing assay (Hernandez, Stein et al., 2012, Hoekstra & Duzgunes, 1993, Yavuz, Kattan et al., 2018), used to monitor the ability of full-length myoferlin and dysferlin to promote tight vesicle-vesicle aggregation or docking, 20 µL empty dual labelled 100 nm LUVs (comprising 1 mol% NBD PE and 1 mol% Rhodamine B PE) were mixed in a 1:1 ratio with unlabelled proteoliposomes in 1 mL reconstitution buffer (20 mM HEPES-KOH, pH 7.5, 150 mM KCl), supplemented with 0.1-1 mM CaCl_2_. The lipid mixing data (**Fig EV4 D-E**) was acquired at 37°C using a Fluorolog 3 spectrofluorometer (Horiba Jobin Yvon) and corrected for signal intensity variations (the S/R acquisition mode). The extent of NBD (donor) dequenching because of lipid mixing was monitored at 460 nm (3 nm slit) and 538 nm (3 nm slit) excitation and emission wavelengths, respectively. The lipid mixing reactions were stopped upon addition of 5 µL 10 % (v/v) Triton X-100 (in reconstitution buffer) and the maximal NBD dequenching signal, after detergent solubilization of liposomes, was considered as the maximal fluorescence (F_max_). The normalized lipid mixing efficiency was calculated as: (F-F_i_)/(F_max_-F_i_), where F_i_ represents the initial fluorescence of the labelled liposomes and F_max_ – the final fluorescence of the sample (i.e., after detergent addition). The dual labelled LUVs comprised 25 mol % DOPS/5 mol% PI(4,5)P_2_ or 15 mol% DOPS, whereas full-length myoferlin and dysferlin were reconstituted in DOPC/DOPE-only LUVs. The control reactions included only protein-free liposomes. The lipid mixing data was analysed in OriginPro 2020 (v9.7.0.188). The lipid mixing assays were repeated at least three times (technical replicates) and at least two separate proteoliposome reconstitutions (biological replicates).

### Mass photometry characterization of full-length human dysferlin

All measurements (**Fig S1H**) were performed with the OneMP mass photometer (Refeyn). Images were acquired with Refeyn AcquireMP and analysed using Refeyn DiscoverMP software. For mass photometry measurements, twin-StrepII-tagged full-length dysferlin (residues 1-2080), was reconstituted into LMNG (Lauryl maltose neopentyl glycol, Anatrace, cat#NG310) micelles (instead of DDM) and purified by anion-exchange chromatography. The dysferlin sample was concentrated to ∼3 mg/mL in the presence of 0.01% (w/v) LMNG and dialysed before measurements using a Slide-A-Lyzer MINI device (MWCO 20 kDa, Thermo Fisher Scientific, cat#69590) against the dialysis buffer (20 mM HEPES-KOH, pH 7.5, 150 mM KCl, 2 mM DTT). The sample was then 20-fold diluted in the dialysis buffer and centrifuged at 13,000 r.p.m. for 10 min at 4°C before the measurements. Prior to loading, for each measurement, 1 µL of sample was added to a droplet of 12 µL of dialysis buffer. For the experiments carried out in the presence of CaCl_2_, the sample after the dialysis was diluted in a dialysis buffer containing 2 mM or 4 mM CaCl_2_ and incubated for 45 – 60 minutes, before performing the measurement. Mass calibration was achieved by adding 6 µL of NativeMark™ Unstained Protein Standard (Invitrogen, cat#LC0725) diluted 100 times (in the dialysis buffer), to a drop of 12 µL of the dialysis buffer and using the peaks corresponding to bovine serum albumin (66 kDa), lactate dehydrogenase (146 kDa) and apo- ferritin (480 kDa). Each experiment was repeated four times (technical replicates).

### Cryo-EM sample preparation

To preserve the integrity of the samples in vitreous ice, lipid-free soluble myoferlin (1-1997) and dysferlin (1-2017) were stabilized through Glutaraldehyde (GA, Electron Microscopy Sciences, cat#16220) crosslinking upon gradient centrifugation (GraFix) (Kastner et al., 2008). Briefly, prior to vitrification, 80 µL soluble dysferlin (1-2017) at a ∼5.5 mg/mL concentration were applied to a linear 5-40% (w/v) sucrose gradient prepared by mixing equal volumes of the light (20 mM HEPES-KOH, pH 7.5, 150 mM KCl, 1 mM CaCl_2_, 5% (w/v) Sucrose) and Glutaraldehyde (GA)-supplemented heavy gradient solution (20 mM HEPES-KOH, pH 7.5, 150 mM KCl, 1 mM CaCl_2_, 40% (w/v) Sucrose, 0.2% (v/v) GA) using the Gradient Master 108 (Biocomp). The gradient was then centrifuged at 4°C for 15 h at 29,100 r.p.m. in a SW40Ti rotor (Beckman Coulter). The gradient was harvested from the top in 500 μL fractions using the Piston Gradient fractionator (Biocomp) and the crosslinker was immediately quenched by the addition of 50 mM L-Lysine and L-Arginine (final concentration), from a 500 mM stock. Following incubation for 2 hours on ice, the monomeric 6-8 fractions were pooled and concentrated to ∼80 µL, transferred to a Slide-A-Lyzer MINI device (Thermo Fisher Scientific, MWCO 10 kDa), and then dialyzed overnight against 2 L of the minimal buffer (20 mM HEPES-KOH, pH 7.5, 150 mM KCl, 1 mM CaCl_2_, 1 mM DTT, 2.5% (v/v) Glycerol). Following dialysis for an additional 2 hours against 1 L minimal buffer, the concentration was adjusted to A280 ∼1.08 (absorbance at 280 nm) and the sample was used directly for cryo-EM grid preparation. Cryo-EM grids were prepared using a Vitrobot Mark IV plunger (Thermo Fisher Scientific), operated at 4°C and 100% humidity. Soluble dysferlin (1-2017) grids suitable for data collection were obtained through application of ∼3 μL crosslinked sample to one side of UltrAuFoil R1.2/1.3 300 gold mesh grids (Jena Bioscience, cat#X-201-AU300), which were pretreated with the Plasma Cleaner (Harrick Plasma, cat#PDC-32G) for 1 min at medium settings prior to their vitrification in liquid ethane, cooled by liquid nitrogen. For optimal sample vitrification the cryo-EM grids were blotted for 2-3 s using a blot force of 5 and stored in liquid nitrogen prior to screening/data collection.

Like dysferlin, lipid-free myoferlin (1-1997) was stabilized through GraFix. However, in contrast to dysferlin (1-2017), myoferlin (1-1997) samples were assembled in a minimal buffer containing 200 mM KCl (25 mM HEPES-KOH, pH 7.5, 200 mM KCl, 2.5% (v/v) Glycerol, 1 mM TCEP), in the presence 0.5 mM CaCl_2_ and 50 µM WJ460 (GlpBio, cat#GC38203), and then centrifuged for 90 min at 50,000 r.p.m. on 5-40% (w/v) sucrose Grafix gradients using a TLS-55 rotor, mounted in a Optima MAX-XP ultracentrifuge (Beckman Coulter). The lipid-free myoferlin (1-1997) gradients were harvested from the top in 100 µL fractions and the crosslinker was quenched with 50 mM L-Lysine and L-Arginine. The peak gradient fractions of myoferlin (1-1997) were concentrated and the buffer exchanged to the minimal buffer (containing 0.5 mM CaCl_2_) by repeated ultrafiltration using a Vivaspin 500 spin concentrator (MWCO 50 kDa, cat#VS0131). Lipid-free myoferlin (1- 1997) was then vitrified in a mixture of liquid ethane and propane (37%:63%) cooled by liquid nitrogen, following the application of 3 µL sample at A280∼1.15 to a plasma-cleaned Quantifoil R1.2/1.3 200 copper mesh grid (Jena Bioscience, cat#X-101-CU200), which was blotted for 7.5 s using a blot force of 3.

To assemble lipid-bound myoferlin (1-1997) complexes (**Appendix Fig S1B-C**), empty nanodiscs with the desired lipid composition were reconstituted in the presence of the MSP2N2 scaffold, as recently described (Cannon et al., 2023). Purified His-tagged human MSP2N2 (∼2.9 mg/mL, obtained from Cube Biotech, cat# 26172) and DDM-solubilized lipids were mixed in ∼1:80-1:200 protein:lipids molar ratios, incubated at room temperature for 20 min, and dialyzed overnight at 4-8°C against 2L of reconstitution buffer (25 mM HEPES-KOH, pH 7.5, 150-200 mM KCl, 1 mM DTT), supplemented with 2.5 g/L Bio-Beads SM-2. Empty nanodiscs were subsequently purified by size-exclusion chromatography (SEC) on Superdex 200 Increase 10/300 GL (Cytiva, cat#28-9909-44), equilibrated in the SEC buffer (25 mM HEPES-KOH, pH 7.5, 150-200 mM KCl, 1.25%-2.5% (v/v) Glycerol, 1 mM DTT (or 0.5 mM TCEP), 0.5 mM CaCl_2_). Peak MSP2N2 nanodisc fractions were concentrated to ∼2.2 mg/mL and added in ∼2-fold molar excess over purified soluble myoferlin (1-1997) in the SEC buffer, followed by incubation for 30 min at room temperature. The formation of stable myoferlin (1-1997)-nanodisc complexes (**Appendix Fig S1B-C**) was assessed by SEC on a Superose 6 Increase 10/300 GL (Cytiva, cat#29-0915-96) column, equilibrated in the SEC buffer. For cryo-EM grid preparation, the myoferlin-nanodisc complexes were crosslinked on ice in batch with 0.05-0.08% (v/v) GA for 10-30 min and the reaction was stopped by quenching with 50 mM L-Lysine and L-Arginine for 15 min on ice. Following centrifugation at 14,800 r.p.m. at 4°C for 5 min, the crosslinked complexes were applied to a Superose 6 column and myoferlin (1-1997)-nanodisc fractions were pooled and concentrated by ultrafiltration to A280∼0.7-0.8. Lipid-bound myoferlin (1- 1997) complexes were vitrified in liquid ethane-propane (37%:63%) following the application of 3 µL sample to plasma-cleaned Quantifoil R1.2/1.3 200 copper mesh grids, which were blotted for 6.5-8 s at a blot force of 3 (except for the 15% DOPS/2 mol% PI(4,5)P_2_ myoferlin (1-1997)-nanodisc complex, frozen on an UltrAuFoil R1.2/1.3 300 gold mesh grid).

### Cryo-EM data collection and processing of lipid-bound myoferlin complexes

Sample size calculation was not performed, and no randomization or blinding was required. All lipid-bound myoferlin cryo-EM datasets were acquired on a high-end Titan Krios G4 electron microscope (Collaborative Laboratory and User Facility for Electron Microscopy, Georg-August-Universität Göttingen), operated at an accelerating voltage of 300 kV and equipped with a Falcon4i direct electron detector and a Selectris X zero-loss energy filter (**Appendix Fig S2, S4 and S5, Tables S1-S4**). All cryo-EM movies were recorded with EPU (Thermo Fisher Scientific) at a 165,000x nominal magnification at the specimen level (resulting in a 0.72 Å/pixel exposure sampling rate), using a slit width of 10 eV and a 50 μm C2 aperture (the objective aperture was not inserted). Cryo-EM movies were stored as raw camera frames (in EER format) following exposure over ∼3.16-3.46 s, which resulted in a total fluence of ∼38.03-39.89 e^-^/Å^2^ (**Appendix Tables S1-S4**). The EER movies were fractionated in 40 EER fractions during on-the-fly preprocessing (patch motion correction, patch CTF estimation, dose-weighting) with cryoSPARC Live (v4.4-v4.5) and curated (contaminated or low-resolution exposures were removed from subsequent analyses). Myoferlin particles were picked first using the Blob Picker and then with the Template Picker (or with crYOLO (Wagner, Merino et al., 2019), **Appendix Fig S5**), extracted in 360 pixel boxes and 2x binned, before being subjected to 3D and 2D classification in cryoSPARC. The initial particle sets were first cleaned by supervised 3D classification (Heterogeneous refinement) using one “good” *ab initio* generated reference volume and 4-5 “junk” classes. Particles assigned to the nanodisc-bound myoferlin classes were, additionally, cleaned in 2D, refined in 3D, and then re-extracted in a 360 pixel box (0.72 Å/pixel) before Non-uniform (NU) refinement in cryoSPARC. To resolve the peripheric C_2_G and the missing inner DysF domain (from some particles), the cryoSPARC-refined particles were next subjected to 3D classification in RELION-5.0-beta (Kimanius et al., 2024). Briefly, the monomeric myoferlin-nanodisc particles were first classified in 3D without image alignment using 4 classes and soft masks applied to the inner DysF domain or to both C_2_G, the inner, and outer DysF (**Appendix Fig S2, S4, and S5**). The subset of particles exhibiting stronger density for the inner DysF and C_2_G were then re-imported and refined in cryoSPARC (**Appendix Fig S2, S4 and S5**). Prior to focused classification, the myoferlin particles of the 15 mol% DOPS/5 mol% Cholesterol nanodisc dataset were subjected to an additional round of 3D classification in 5 classes with image alignment (**Appendix Fig S5**). For all lipid-bound myoferlin complexes, the final particle sets were subjected to sequential CTF refinement and reference-based motion correction in cryoSPARC to obtain the final maps. To facilitate model building, the lipid-bound myoferlin maps were sharpened using LocScale (Jakobi et al., 2017) and DeepEMhancer (Sanchez-Garcia et al., 2021), and the local resolution of the maps was estimated in cryoSPARC. The continuous flexibility and conformational space of membrane-bound myoferlin particles were evaluated in cryoSPARC using the 3D variability analysis (3DVA) tool.

### Cryo-EM data collection and image analysis of lipid-free myoferlin and dysferlin datasets

As in the case of the membrane-bound samples, the lipid-free myoferlin and dysferlin datasets were acquired using EPU on the same microscope and detector (Falcon4i/Selectris), with energy filter set to slit widths of 10 eV or 15 eV (**Appendix Fig S6 and Fig EV3, Tables S5-S6**). Lipid-free dysferlin cryo-EM movies were recorded at a nominal magnification of 165,000x (pixel size of 0.72 Å per pixel), from the same grid in two sessions over a ∼3.0 s exposure, resulting in a total fluence of ∼40.22 e^-^/Å^2^. Cryo-EM movies of the lipid-free myoferlin particles were collected in two sessions at total fluences of ∼39.91 e^-^/Å^2^ and ∼39.85 e^-^/Å^2^, respectively, and saved as raw EER frames. Lipid-free ferlin datasets were preprocessed and curated on-the-fly in cryoSPARC Live (v4.1 (lipid-free dysferlin) and v4.4- v4.5 (lipid-free myoferlin)), including beam-induced motion correction, dose-weighting, patch CTF estimation, and particle picking (using the Blob Picker and Template Picker). Particles from “good” micrographs were extracted in a 360 pixel box (0.72 Å/pixel), 2x binned, split into subsets of ∼0.7-1 mln particles each (**Appendix Fig S6 and Fig EV3**), before supervised 3D classification in cryoSPARC with 4 or 5 classes, which included an *ab initio* generated ferlin reference volume. Following an additional 2D classification, the lipid-free myoferlin and dysferlin particles were refined in 3D to obtain the consensus maps, which exhibited strong density of the Fer^core^ region. The C_2_F and C_2_E domains could also be resolved in the lipid-free myoferlin consensus maps (**Fig EV3**). As C_2_B and the C-terminal C_2_F-C_2_G appeared dynamic in the imaged lipid-free ferlin particles, multiple rounds of global and focused 3D classifications were employed, with the aim of finding more ordered conformations of the domains. Consequently, focused classification with soft mask applied to the C_2_F and C_2_G domain regions of the map, allowed us to identity more homogenous dysferlin particles, which refined in 3D to 3.5 Å (dysferlin (1-2017) map M2) and 4.8 Å (dysferlin (1-2017) map M3) resolutions. In these maps, C_2_F appears to establish a defined interface with the N-terminal C_2_B, whereas C_2_G occupies a location near C_2_F. By employing a similar 3D classification approach, we could also resolve the C_2_B-C_2_F-C_2_G interface in a subset of lipid-free myoferlin particles (**Fig EV3**). These particles refined in 3D to 3.2 Å (myoferlin (1-1997) map M9) and 8.8 Å (myoferlin (1-1997) map M11) global resolutions and are reminiscent of the lipid-free dysferlin (1-2017) maps. Attempts to locate the N-terminal C_2_A domain by masked 3D classification were not successful, consistent with it not sharing significant contact interfaces with the remaining ferlin C_2_ domains or with it being destabilized in vitreous ice.

### Model building and refinement

Model building was initiated from Alphafold2 predictions of human myoferlin and dysferlin (Jumper et al., 2021). The theoretical models were divided into individual domains (or structural modules, such as the Fer^core^), fitted into the overall ferlin maps in ChimeraX (v1.6- v1.8) and UCSF Chimera (v1.16), and their fitting adjusted with ISOLDE (Croll, 2018). All structural models were corrected and rebuilt in Coot (v0.9.8.5) (Emsley & Cowtan, 2004) and iteratively refined with phenix.real_space_refine (Phenix v1.19.2-4158-v1.21.2-5419) (Afonine et al., 2018) with nonbonded_weight set to 1000 for the final refinements. Modelling of bound Ca^2+^ ions and phospholipid headgroups was enabled by *F_o_*-*F_c_* omit maps, calculated with Servalcat (Yamashita, Palmer et al., 2021). Geometry restraints for phospholipid refinement were generated with Grade2 (https://grade.globalphasing.org). No cryo-EM density was observed for the WJ460 ligand (Zhang et al., 2018), added to the lipid-free myoferlin (1-1997) and the 15 mol% DOPS/2 mol% PI(4,5)P_2_ nanodisc-myoferlin complex. As the proposed binding site residues of C_2_D are not readily accessible in the lipid-free and lipid-bound myoferlin cryo-EM structures, further studies are likely needed to clarify whether myoferlin is indeed the cellular target of the small-molecular compound (Zhang et al., 2018). All structural models were validated with MolProbity in Phenix. Structural figures were prepared with ChimeraX (v1.6-v1.8) and Pymol (v2.5.4, Schrödinger LLC). The contact interface areas were estimated with PISA (v1.5.2) (Krissinel & Henrick, 2007). Data collection and refinement statistics are provided in **Appendix Tables S1-S6**.

### Chemical crosslinking mass spectrometry characterization of human dysferlin

Prior to mass spectrometry analysis (**Appendix S14A-D**), purified soluble dysferlin (1-2017) was complexed with 1.5 mM CaCl_2_ and crosslinked in batch with 0.3 mM BS3 (Thermo Fisher Scientific, cat#A39266) for 30 min at room temperature. The crosslinking reaction was stopped with 50 mM Tris-HCl, pH 8.0 and the sample was further purified by size-exclusion chromatography on Superose 6 Increase 10/300 GL (Cytiva), equilibrated in the sample buffer (20 mM HEPES-KOH, pH 7.5, 150 mM KCl, 5% (v/v) Glycerol, 2 mM DTT, 1.5 mM CaCl_2_). Individual monomeric peak fractions (fraction 14 and 15) were ethanol precipitated and resuspended in 12.5 µl 8 M Urea. The samples were then diluted to 2 M Urea end concentration and reduced by the addition of 10 µl 50 mM DTT for 30 minutes at 37 °C. Alkylation was achieved by addition of 10 µl 200 mM iodoacetamide and incubation for 30 minutes at 25 °C. Unreacted iodoacetamide was quenched with additional 10 µl 50 mM DTT. Protein digestion was performed overnight at 37 °C with the addition of 1 µg of trypsin (Promega) in the presence of 1 M Urea and 50 mM Tris-HCl, pH 7.75. The samples were acidified with formic acid to 0.1% (v/v) end concentration and acetonitrile was added to 5% (v/v) end concentration. Digested peptides were desalted with C18 reversed-phase MicroSpin columns (Harvard Apparatus). Bound peptides were eluted with 50 % (v/v) acetonitrile, 0.1% (v/v) formic acid. The samples were subsequently dried under vacuum and resuspended in 75 µl 2% (v/v) acetonitrile, 0.05% (v/v) TFA and 5 µl were used for LC-MS analysis.

Chromatographic separation was achieved with Dionex Ultimate 3000 UHPLC (Thermo Fischer Scientific) coupled with C18 column packed in-house (ReproSil-Pur 120 C18-AQ, 1.9 µm particle size, 75 µm inner diameter, 33 cm length, Dr. Maisch GmbH) over a 74 minute linear gradient from 8% - 46% mobile phase B (mobile phase A - 0.1% (v/v) FA and B - 80% (v/v) ACN, 0.08% (v/v) FA). Eluting peptides were analyzed with Orbitrap Exploris 480 (Thermo Fischer Scientific) and the following settings for survey scans: Resolution – 120 000; Scan range – 380-1600; AGC target – 300%; Maximum injection time set to “Auto”. Analytes with charge state 3 to 8 were selected for fragmentation with 28% normalized collision energy. Dynamic exclusion was set to 15 s. Fragment spectra were acquired with the following settings: Resolution – 30 000; Isolation window – 1.6 *m/z*; AGC target – 100%; Maximum injection time – 128 ms. Resulting .raw files were analyzed with pLink (Chen et al., 2019) (v 2.3.9) against a database containing the sequence of the protein. Carbamidomethyl on cysteines was selected as fixed modification and oxidation of methionines as variable modifications. BS3 was selected as crosslinker, peptide tolerance was set to 6 ppm and False discovery rate was set to 1%. The identified crosslinked residues (**Dataset EV1**) were mapped onto the Alphafold2 prediction of dysferlin and the lipid-free dysferlin (1-2017) cryo-EM model using xiNet (Combe, Fischer et al., 2015), analysed and visualized in PyMOL (v2.5.4).

**Expanded View** for this article is available online.

## Acknowledgements

We thank Christiane Senger-Freitag, Sandra Gerke, Sina Langer, Andy Schacht, and Patricia Räke-Kügler for excellent technical and administrative support. We are grateful to Dr. Tat Cheung Cheng and Prof. Dr. Ruben Fernandez-Busnadiego for supporting cryo-EM data acquisition and Dr. Eri Sakata for critically reading the manuscript. This work was supported and funded by Deutsche Forschungsgemeinschaft (DFG) through CR 937/2-1 (to CC) and the Cluster of Excellence (EXC2067) Multiscale Bioimaging EXC 2067/1-390729940 (CC, TM and JP). CZKS and VP were supported by an Wellcome Trust Investigator grant (220300Z/20/Z). HU was supported by the Max Planck Society. Cryo-EM instrumentation was jointly funded by the DFG Major Research Instrumentation program (448415290) and the Ministry of Science and Culture of the State of Lower Saxony. TM was also supported by the Leibniz Program of the DFG (MO896/5), the Ernst Jung Prize for Medicine, by Fondation Pour l’Audition (FPA RD-2020-10)

## Author contributions

CC, TM and JP designed the research. CC engineered and cloned constructs, prepared samples for structural studies, collected and processed cryo-EM data. CC built, refined and interpreted the ferlin structural models. CC designed and performed functional assays, under the supervision of JP and TM. CC and AC prepared mass spectrometry samples. AC collected and analysed the proteomics and chemical crosslinking mass spectrometry data under the supervision of HU. CZKS and VP carried out the mass photometry measurements and contributed to data analysis. CC wrote the paper under the supervision of TM and JP and with input from all the authors. CC, TM and JP acquired funding for the project.

## Data availability

The atomic coordinates of the lipid-bound myoferlin were deposited in the Protein Data Bank (PDB) and Electron Microscopy Data Bank (EMDB) with the following accessing codes: PDB 9H6X/EMD-51902 (lipid-bound human myoferlin, 25 mol% DOPS/5 mol% PI(4,5)P_2_ nanodisc).

## Extended View Figure Legends

**Figure EV1.**
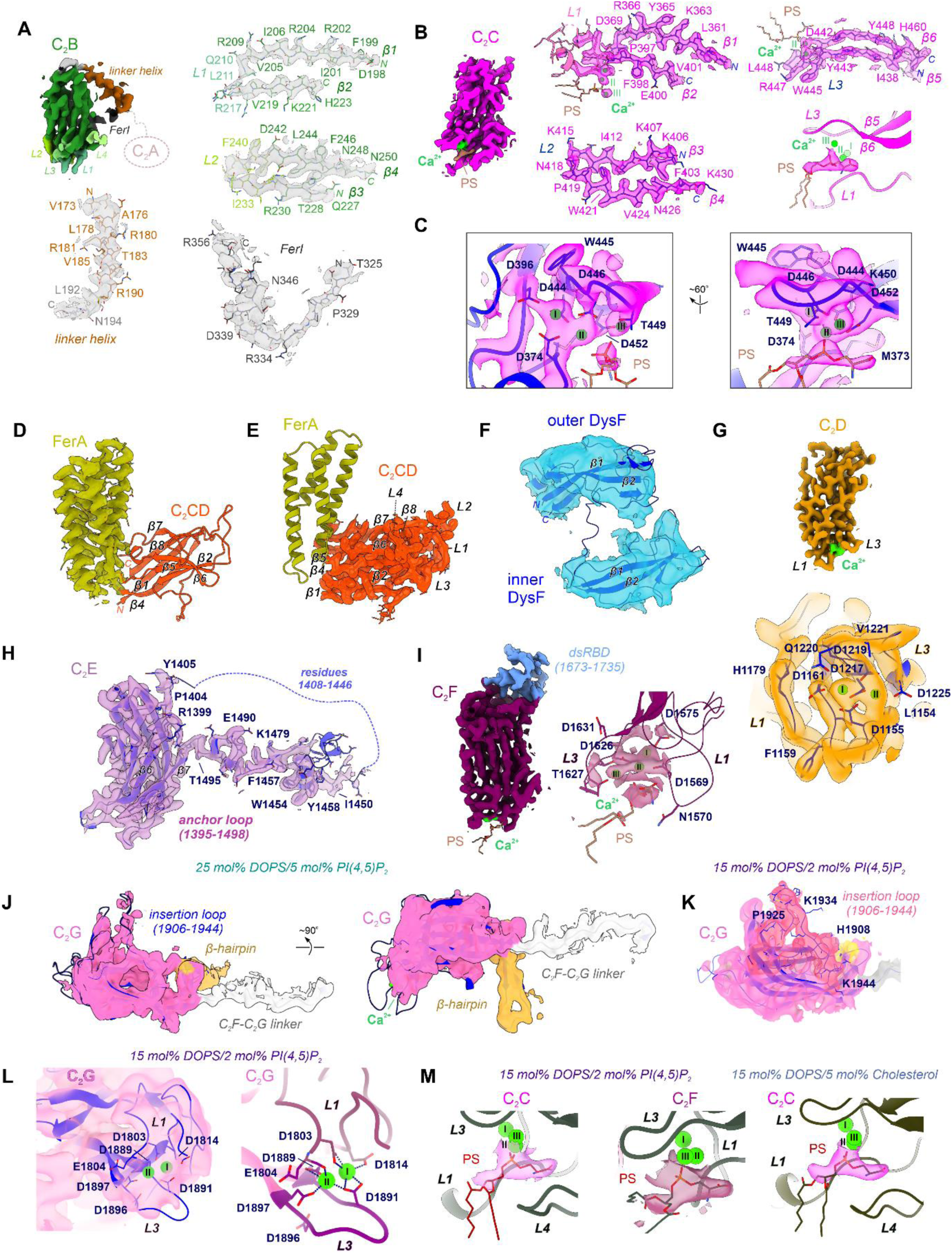
Cryo-EM density snapshots of the lipid-bound soluble myoferlin (1-1997). **A** Selected cryo-EM density snapshots of C_2_B and of its proximal motifs (the linker helix and the FerI motif) cryo-EM density. Myoferlin’s residues are depicted as sticks. **B** Cryo-EM density of myoferlin’s C_2_C domain. The three bound Ca^2+^-ions and the recruited phosphatidylserine (PS) are indicated. **C** Ca^2+^-binding sites of the C_2_C domain of myoferlin. The cryo-EM density is contoured around the Ca^2+^-binding residues. **D** Cryo-EM density of the FerA motif of myoferlin. Note that the four-helix bundle domain inserts between the β4 and β5 strands of the C_2_CD domain (red, shown in a cartoon representation). **E** Cryo-EM density of the C_2_CD domain of myoferlin. The seven β-strands of the domain are indicated. **F** Cryo-EM densities of the modelled inner and outer DysF motifs of myoferlin. **G** Cryo-EM density and the observed Ca^2+^-binding sites of the C_2_D domain of myoferlin. C_2_D’s cryo-EM density is contoured around the Ca^2+^-binding sites (bottom panel). **H** Density of the C_2_E domain. The domain comprises an extended insertion loop (denoted as the anchor loop), inserted between the β6-β7 strands. The loop establishes contacts with the downstream C_2_F domain. **I** Cryo-EM density of C_2_F and of its Ca^2+^-/lipid-binding sites. As in the case of C_2_C, three Ca^2+^ ions and a PS headgroup could be assigned in the cryo-EM density map. **J** Cryo-EM density of the C-terminal C_2_G domain, originating from the 25 mol% DOPS/5 mol% PI(4,5)P_2_ myoferlin-nanodisc complex, shown in two orientations. The lipid-binding β- hairpin motif of C_2_G is coloured in light orange. **K** Density of the C-terminal C_2_G domain, resolved in the 15 mol% DOPS/2 mol% PI(4,5)P_2_ myoferlin-nanodisc complex. Note that the insertion loop of C_2_G (residues 1906-1944) is ordered in this myoferlin complex. **L** Modelled Ca^2+^-binding sites of C_2_G, observed in the lipid-bound myoferlin structure (15 mol% DOPS/2 mol% PI(4,5)P_2_). Because of the lower local resolution, Ca^2+^ modelling was initiated by Alphafold3 predictions (Abramson, Adler et al., 2024), and the optimal sites were refined against the cryo-EM map. Our modelling (right panel) suggests that two Ca^2+^ ions are coordinated the L1 and L3 loops of C_2_G, in a phospholipid-independent manner (Rizo & Sudhof, 1998). **M** Cryo-EM densities of the Ca^2+^-bound phosphatidylserine (PS) moieties, resolved in the myoferlin-nanodisc complexes. The cryo-EM density is contoured around the modelled phospholipid headgroups, and the ligands are depicted as sticks.

**Figure EV2.**
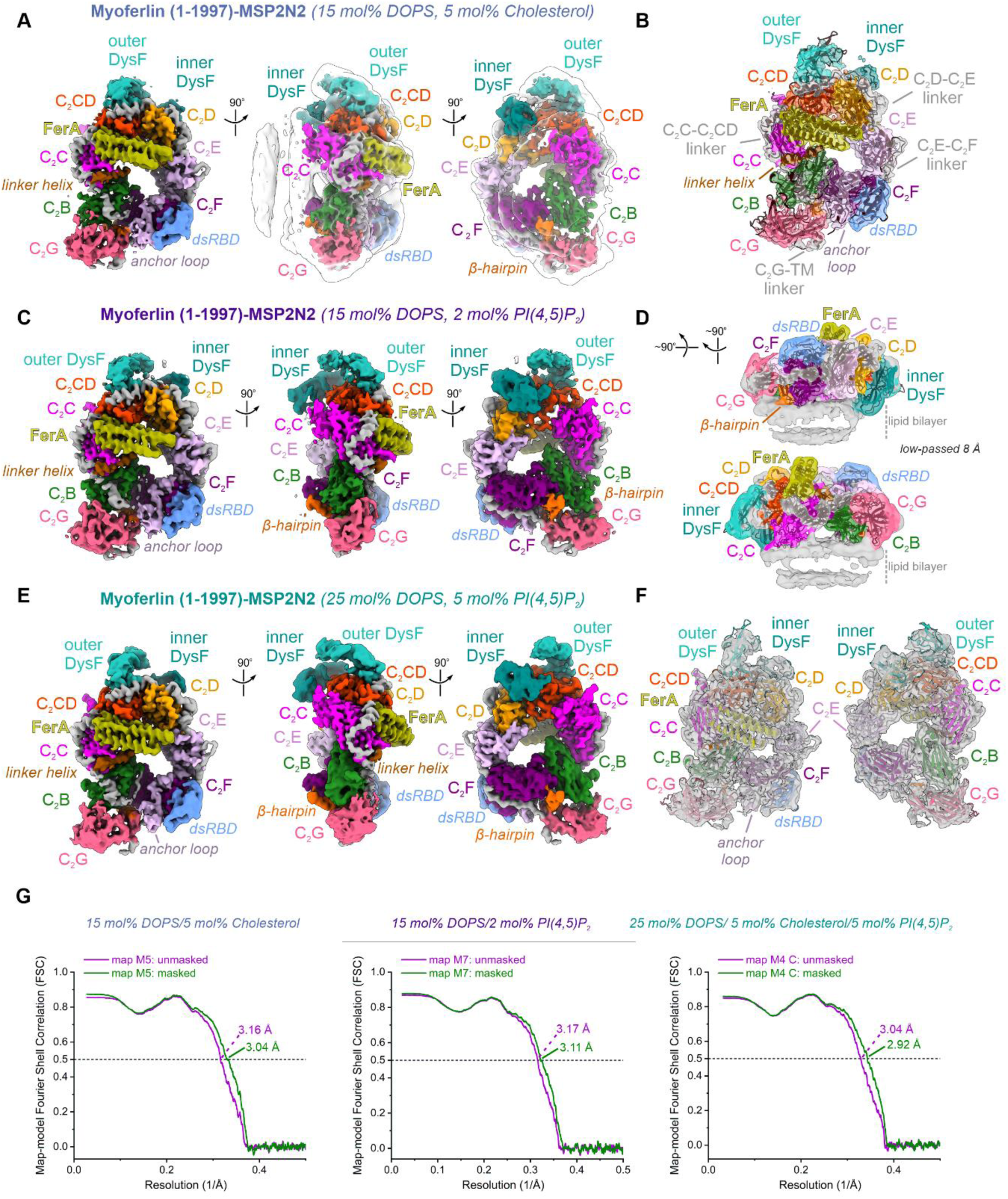
Structural comparison between the different lipid-bound myoferlin complexes. **A** Overall cryo-EM map (map M6, ∼3.3 Å) of soluble myoferlin (1-1997) bound to an MSP2N2 nanodisc comprising 15 mol% DOPS and 5 mol% Cholesterol. The map is depicted in three different orientations and color-coded. A low-passed map (transparent surface) is superimposed to visualize the nanodisc density. **B** The map M6 of the myoferlin (1-1997)-MSP2N2 complex with the final model fitted inside. Note the well-resolved C_2_ domain linker regions, the linker helix, and the anchor loop of C_2_E. **C** Overall cryo-EM map (map M8, ∼3.43 Å) of the soluble myoferlin (1-1997) bound to an MSP2N2 nanodisc comprising 15 mol% DOPS and 2 mol% PI(4,5)P_2_. The map is depicted in three different orientations as in **A**. **D** Side-views of the map M8 (15 mol% DOPS/2 mol% PI(4,5)P_2_) of the lipid-bound myoferlin (1-1997) together with the fitted model. The map has been low-pass filtered to 8 Å and the ordered nanodisc density is indicated. **E** Cryo-EM map of the soluble myoferlin (1-1997)-lipid complex assembled onto a 25 mol% DOPS/5 mol% PI(4,5)P_2_ MSP2N2 nanodisc (∼2.56 Å, map M3). **F** Overall map of lipid-bound myoferlin (1-1997) complex (map M3) with the fitted final model. Except the flexible N-terminal C_2_A, all the ordered myoferlin domains (C_2_B-C_2_G) and accessory motifs (FerA, the linker helix, the anchor loop) could be assigned, resulting in the near-complete structure of ferlin’s cytosolic region. **G** Map versus model Fourier Shell Correlation (FSC) plots for the myoferlin (1-1997)- nanodisc complexes.

**Figure EV3.**
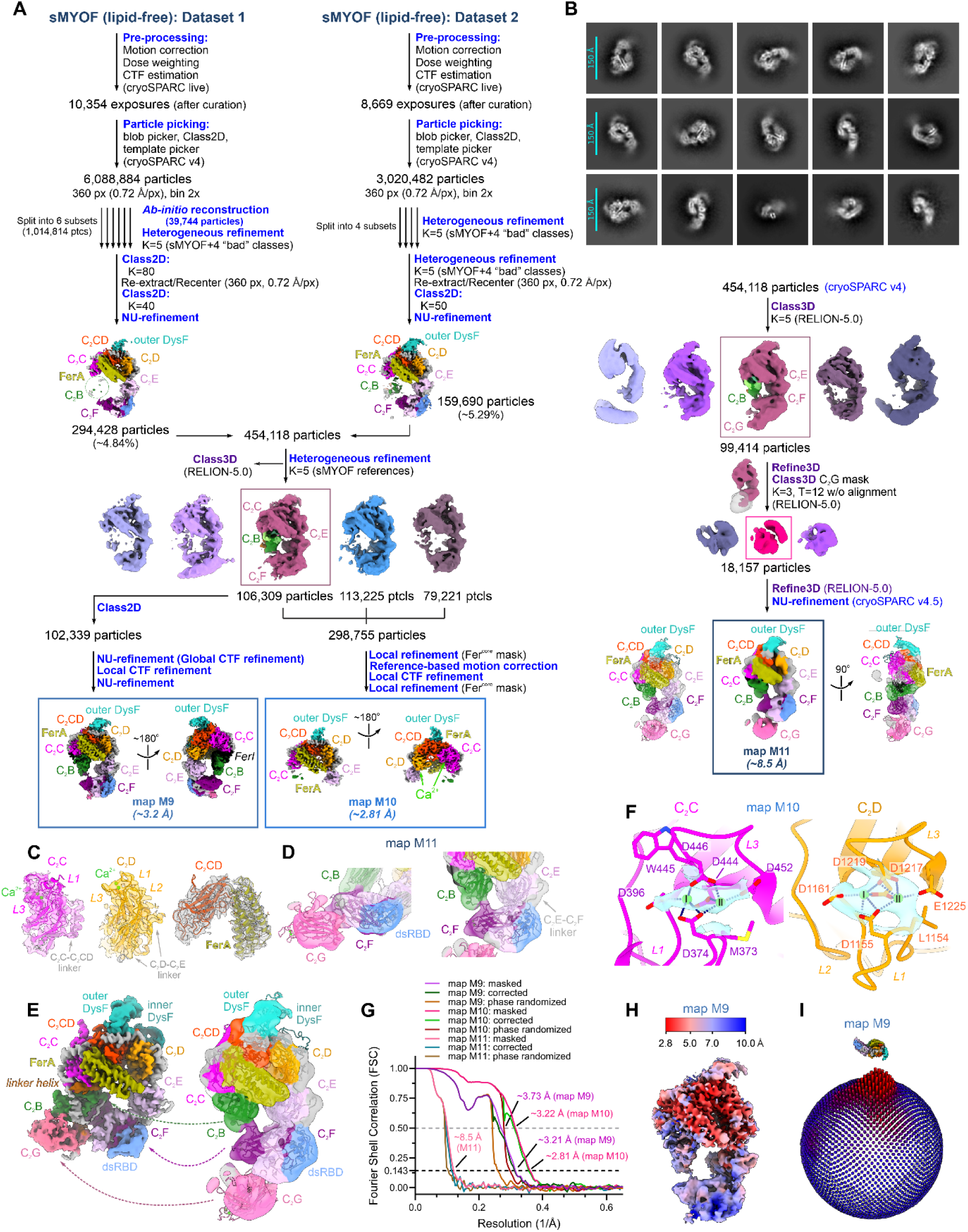
Cryo-EM image analysis of the vitrified lipid-free soluble myoferlin. **A** Cryo-EM image processing schematic for the Ca^2+^-bound, lipid-free myoferlin (1-1997). Resolution of the final maps (M9, M10, and M11 maps) was estimated according to the gold-standard Fourier Shell Correlation (FSC) criterion of 0.143. The structural domains of myoferlin are color-coded as in Fig 1. **B** Reference-free 2D class averages of lipid-free myoferlin (1-1997). Note the similarities between the dysferlin (1-2017) (**Appendix Fig S6C**) and myoferlin (1-1997) 2D class averages in their lipid-free states. **C** Cryo-EM density snapshots of the myoferlin (1-1997) C_2_C, C_2_D and C_2_CD-FerA domains. The final model is fitted inside and the Fer^core^ map of myoferlin (map M10) is depicted as a transparent surface. **D** Cryo-EM density of the C_2_F (map M9) and C_2_G (map M11) domains as modelled in the lipid-free myoferlin (1-1997) structure. Note the presence of tertiary interfaces between the C_2_F-C_2_B and C_2_F-C_2_G, as also observed in the lipid-free dysferlin (1-2017) structure. **E** Side-by-side comparison between the membrane-bound (map M3) and lipid-free (map M11) myoferlin structures. The domains undergoing significant displacement upon membrane binding (C_2_B, C_2_F, and C_2_G) are indicated with dashed arrows. **F** Ca^2+^-binding sites observed in the lipid-free myoferlin structure. The cryo-EM density is coloured in cyan and depicted as a transparent surface. The two modelled Ca^2+^ ions, bound to C_2_C and C_2_D, are coloured green. **G** Global resolution estimates for the lipid-free myoferlin (1-1997) cryo-EM maps using Fourier Shell Correction (FSC) between half maps. The resolution estimates at FSC=0.143 and FSC=0.5 are indicated. **H** Local resolution of the overall map of the lipid-free soluble myoferlin (map M9). The map regions coloured in red indicate a higher resolution. **I** Angular distribution of the myoferlin particles contributing to the M9 overall map. The M9 map is shown above the 3D distribution representation. Relative cylinder height and the red colour indicate a higher number of particle images.

**Figure EV4.**
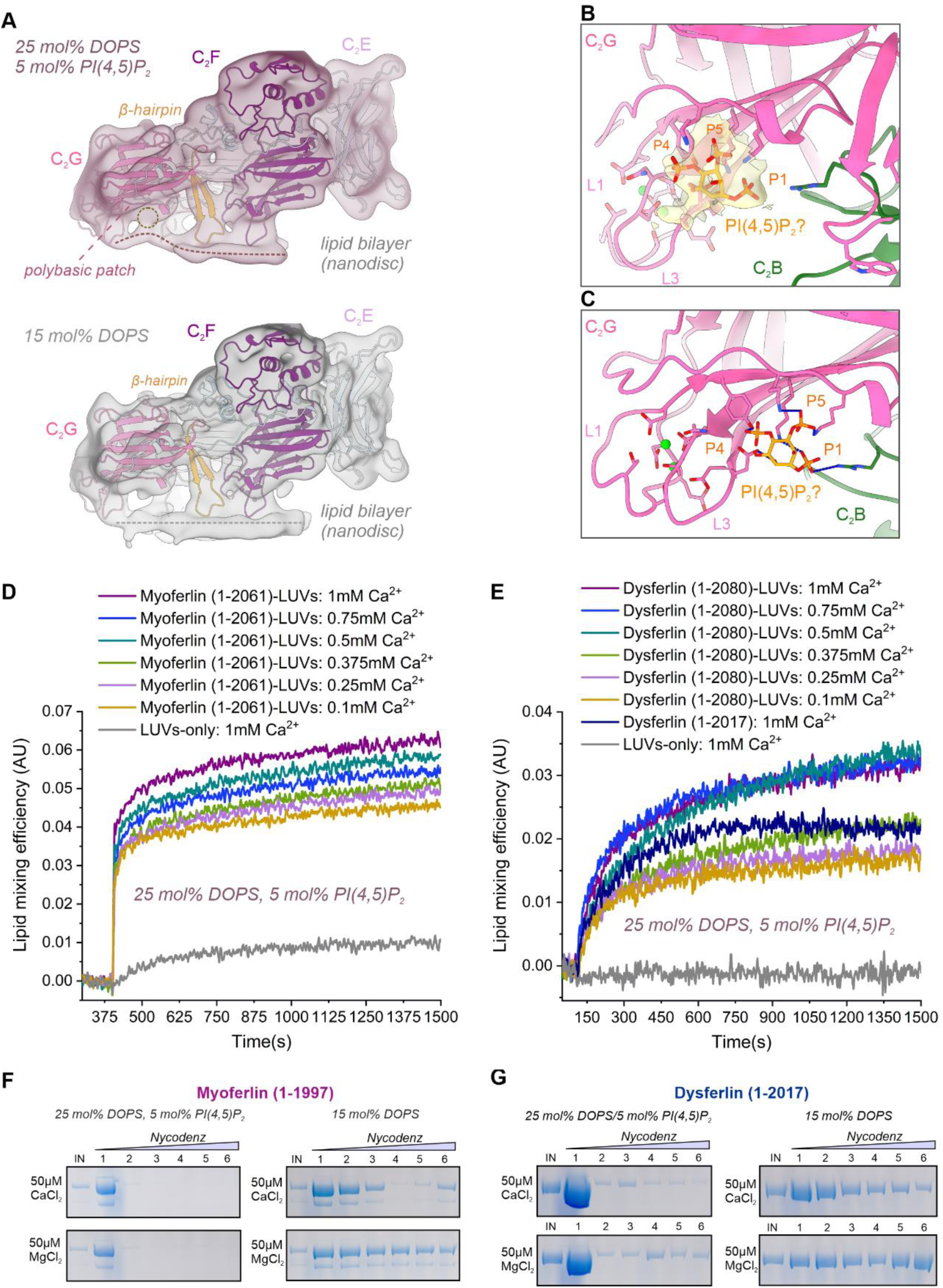
Full-length myoferlin and dysferlin appear to promote tight binding between vesicles in lipid mixing assays. **A** Comparison between the local lipid bilayer structure observed in the myoferlin (1-1997)- nanodisc cryo-EM maps. Note that the lipid bilayer forms close contacts with the concave surface of C_2_G in the presence (top), but not in the absence of PI(4,5)P_2_ (bottom) in the assembled nanodiscs. The cryo-EM maps have been low-passed to 10 Å and the myoferlin model is fitted inside. **B** The C_2_G-proximal cryo-EM density element, with a tentatively modelled PI(4,5)P_2_ headgroup. C_2_G and C_2_B residues, located at the interface, are displayed as sticks. The modelled PI(4,5)P_2_ is located in close proximity to three lysine residues (K1830, K1828 and K1816), projecting from the concave surface of C_2_G. Note the proximal location of the Ca^2+^- binding L3 loop of C_2_G. **C** Polar contacts between C_2_G residues of the polybasic patch and a tentative acidic phospholipid, possibly PI(4,5)P_2_, originating from the nanodisc bilayer (comprising 25 mol% DOPS and 5 mol% PI(4,5)P_2_). **D** Lipid mixing assays between myoferlin proteoliposomes and DOPS/PI(4,5)P_2_-bearing vesicles in the presence of different concentrations of Ca^2+.^ The fluorescent liposomes (100 nm LUVs, 25 mol% DOPS and 5 mol % PI(4,5)P_2_) were labelled with the Lissamine Rhodamine B-NBD (Nitro-2,1,3-benzoxadiazole-4-yl) FRET (Förster Resonance Energy Transfer) pair and the NBD dequenching signal, due to lipid dilution, was used to monitor the extent of tight vesicle-vesicle docking (and, possibly, fusion) to non-fluorescent proteoliposomes. LUVs lacking full-length myoferlin (1-2061) were used as a control. The lipid mixing fluorescent traces were smoothened (using the Savitzky-Golay method with 10 points) and normalized to the maximal dequenching signal to calculate the lipid mixing efficiency. The initial jump in fluorescence is likely due to partial, Ca^2+^-independent myoferlin-induced aggregation of proteoliposomes. The assays were repeated at least three times and representative fluorescent traces are shown. **E** Lipid mixing assays between dysferlin proteoliposomes and DOPS/PI(4,5)P_2_-containing vesicles. The ability of the full-length dysferlin (1-2080) to promote tight vesicle-vesicle docking (or, possibly, vesicle-vesicle fusion) was monitored as a function of Ca^2+^ concentration. The lipid mixing activity of soluble dysferlin (1-2017) was also tested (deep blue). Empty LUVs were used as a control. The assays were repeated at least three times and representative fluorescent traces are shown. **F-G** Ca^2+^-dependent liposome binding by soluble myoferlin (1-1997) and dysferlin (1-2017) using a coflotation assay. The liposomes used in the assay had a similar lipid composition as in **D-E**. The Nycodenz gradients were harvested from the top and analysed by SDS-PAGE. Note the increased liposome binding by both myoferlin (1-1997) and dysferlin (1-2017) in the presence of PI(4,5)P_2_-containing LUVs.

## Appendix

**Figure S1.**
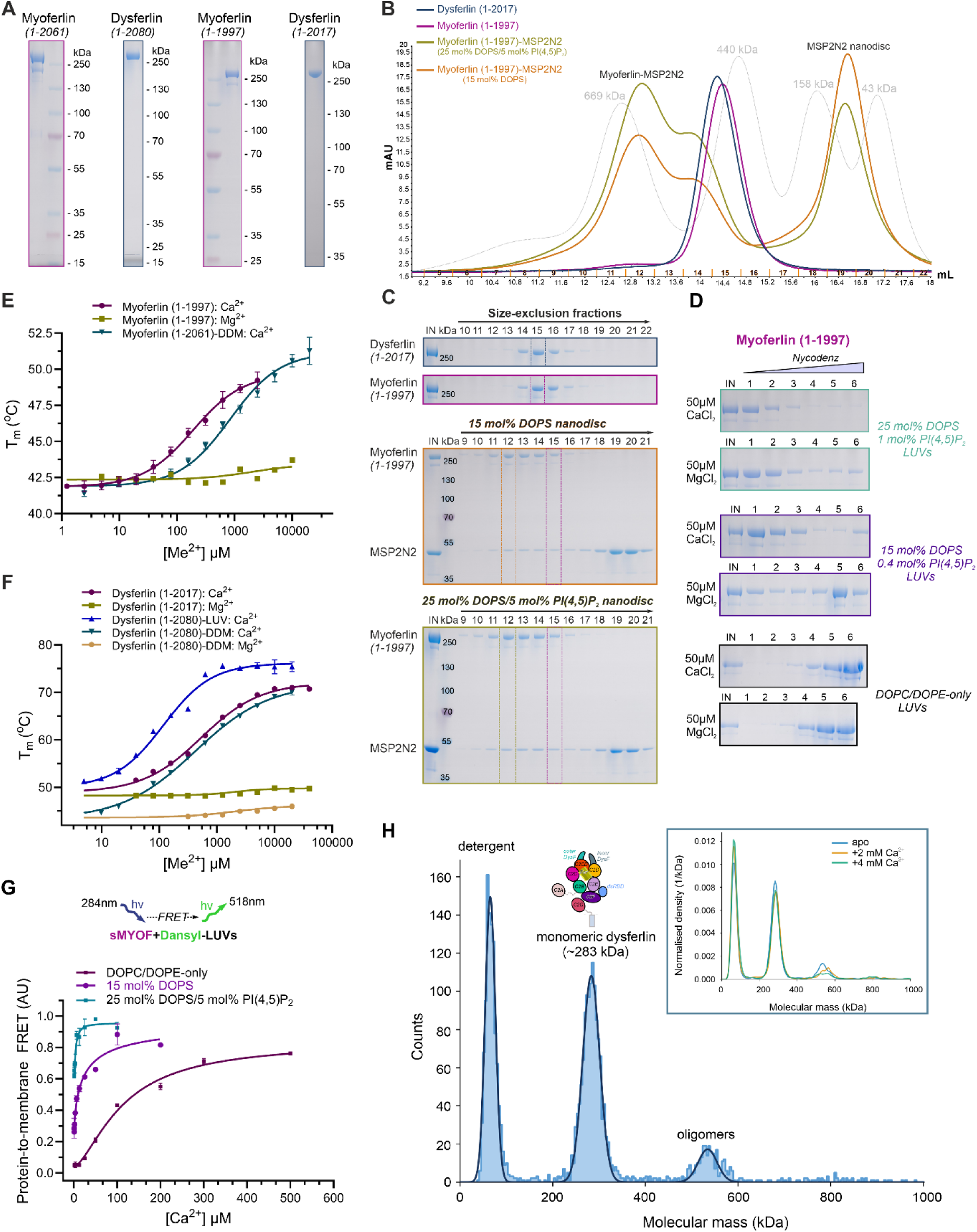
Biochemical characterization of human myoferlin and dysferlin. **A** SDS-PAGE analysis of the purified, insect cell expressed myoferlin and dysferlin constructs used in structural and functional analyses. **B** Size-exclusion chromatography (SEC) elution profile of the soluble myoferlin (residues 1-1997, sMYOF), soluble dysferlin (residues 1-2017, sDYSF), and the reconstituted myoferlin (1-1997)-MSP2N2 complexes. The MSP2N2-based nanodiscs, comprising 15 mol% DOPS or 25 mol% DOPS/5 mol% PI(4,5)P_2_ anionic phospholipids, were purified through SEC prior to the assembly of their complexes with sMYOF. Gel filtration standards (dashed lines) were applied to the Superose 6 column in the same buffer and their molecular masses are indicated. **C** SDS-PAGE analysis of the size-exclusion chromatography peak fractions from **B.** To allow the compact visualization of the myoferlin (1-1997) and dysferlin (1-2017) fractions, the top two gels were cropped (see also **S1A**). **D** Ca^2+^-sensitive liposome binding activity of soluble myoferlin (1-1997) using a coflotation assay. The myoferlin-liposome (LUVs) complexes were assembled *in vitro* in the presence of 50 µM Ca^2+^ or Mg^2+^ and subjected to ultracentrifugation on a Nycodenz step gradient, which was harvested from the top. The anionic phospholipid compositions of the tested LUVs are indicated. LUVs lacking anionic phospholipids (DOPC/DOPE-only LUVs) were used as a control. **E** Ca^2+^-binding activity of membrane-anchored, full-length myoferlin (residues 1-2061, in detergent micelles) and soluble myoferlin (residues 1-1997) using a nanoDSF-based assay. Error bars represent the standard error of the mean (s.e.m., soluble myoferlin (1-1997): n=4, full-length myoferlin (1-2061)-DDM: n=7). [Ca^2+^]_1/2_ was estimated by nonlinear regression curve fit and the Mg^2+^ titration was used as a control. **F** Ca^2+^-binding activity of full-length dysferlin (residues 1-2080, in detergent micelles, DDM), of soluble dysferlin (residues 1-2017) and of dysferlin proteoliposomes (dysferlin (1-2080)-LUVs) using a nanoDSF-based assay. The dysferlin proteoliposomes were prepared by dialysis of the detergent reconstituted full-length sample in the presence of extruded LUVs (assembled from 70 mol% DOPC, 20 mol% DOPE and 10 mol% Cholesterol). The error bars represent the s.e.m. (dysferlin (1-2017): n=4, dysferlin (1-2080)-DDM: n=6, dysferlin (1-2080)-LUV: n=6). **G** Ca^2+^-sensitive liposome binding activity of soluble myoferlin (residues 1-1997, sMYOF) using a FRET-based lipid-binding assay. Dansyl-labelled LUVs, prepared with DOPS (15 mol% DOPS), DOPS and PI(4,5)P_2_ (25 mol% DOPS and 5 mol% PI(4,5)P_2_) or without anionic phospholipids (DOPC/DOPE-only), were used at a 50 µM concentration. Error bars represent s.e.m. (n=3). **H** Mass-photometry characterization of full-length dysferlin (residues 1-2080) reconstituted in detergent micelles (LMNG), showing a major population of monomeric dysferlin particles (molar mass of ∼283 kDa). The inset depicts the mass distributions of dysferlin’s major species at two Ca^2+^ concentrations (2 mM CaCl_2_ and 4 mM CaCl_2_, orange and green curves, respectively), compared to the apo state (blue). In all cases, the mass photometry measurements were carried out at a ∼50 nM dysferlin concentration and in a sample buffer devoid of detergent. A dysferlin domain composition schematic is shown above the mass photometry profile of the ferlin. The mass photometry measurements were repeated four times (n=4).

**Figure S2.**
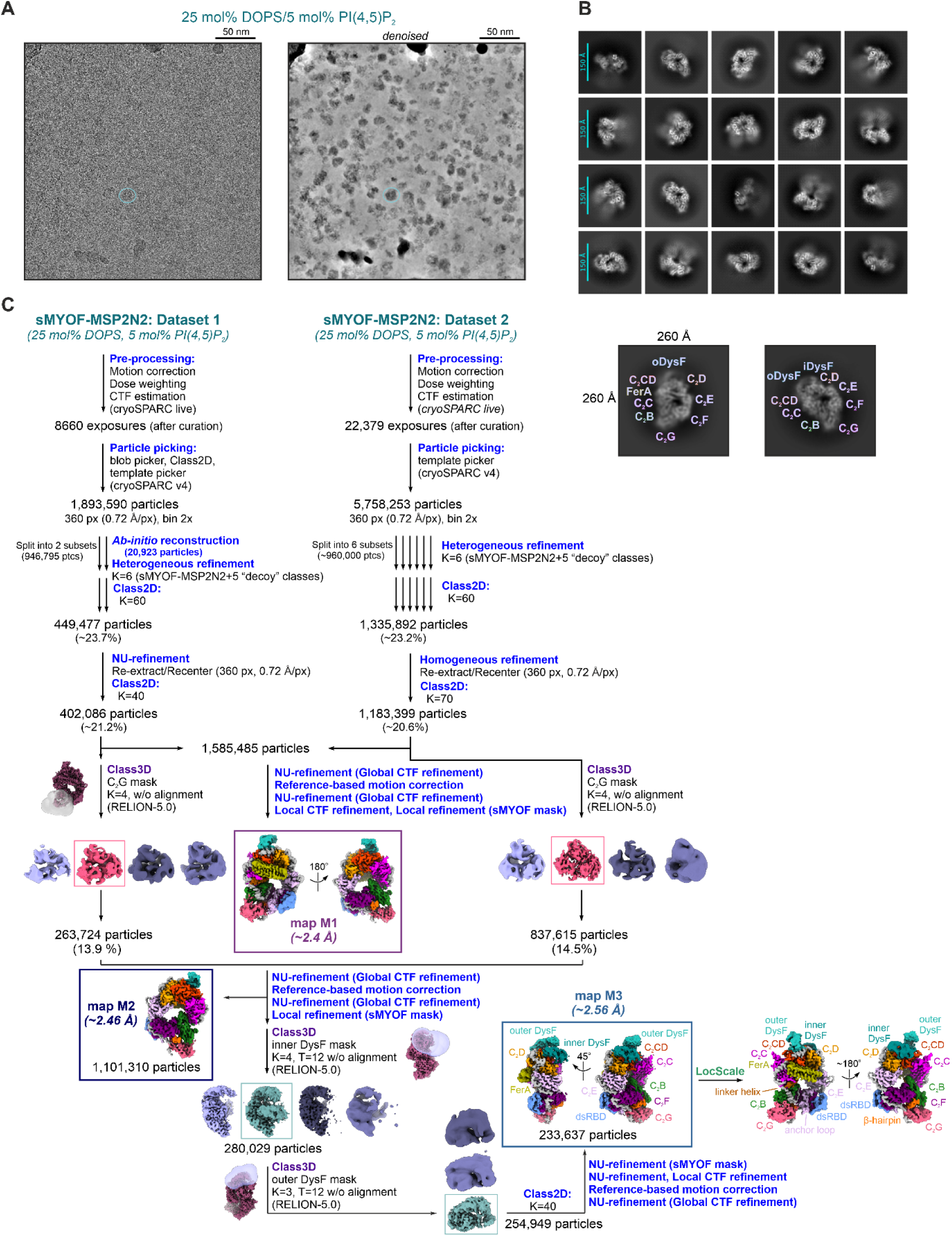
Computational image analysis of the vitrified myoferlin-MSP2N2 nanodisc complex (25 mol% DOPS/5 mol% PI(4,5)P_2_ cryo-EM datasets). **A** Cryo-EM micrograph of the vitrified myoferlin (1-1997)-MSP2N2 complex (25 mol% DOPS and 5 mol% PI(4,5)P_2_ nanodisc). A characteristic, monomeric particle is circled in cyan, in both the raw (left) and denoised (right) exposure. **B** Reference-free 2D class averages of the myoferlin (1-1997)-MSP2N2 complex (comprising 25 mol% DOPS and 5 mol% PI(4,5)P_2_). The visible myoferlin domains are indicated on a characteristic top view of the complex (bottom panel). **C** Cryo-EM data processing routine of the lipid-bound myoferlin, assembled on a 25 mol% DOPS/5 mol% PI(4,5)P_2_ MSP2N2 nanodisc (see also **Fig S1B-C**). The final maps are boxed and colored-coded after the modeled domains (*e.g.*, as in Fig 1).

**Figure S3.**
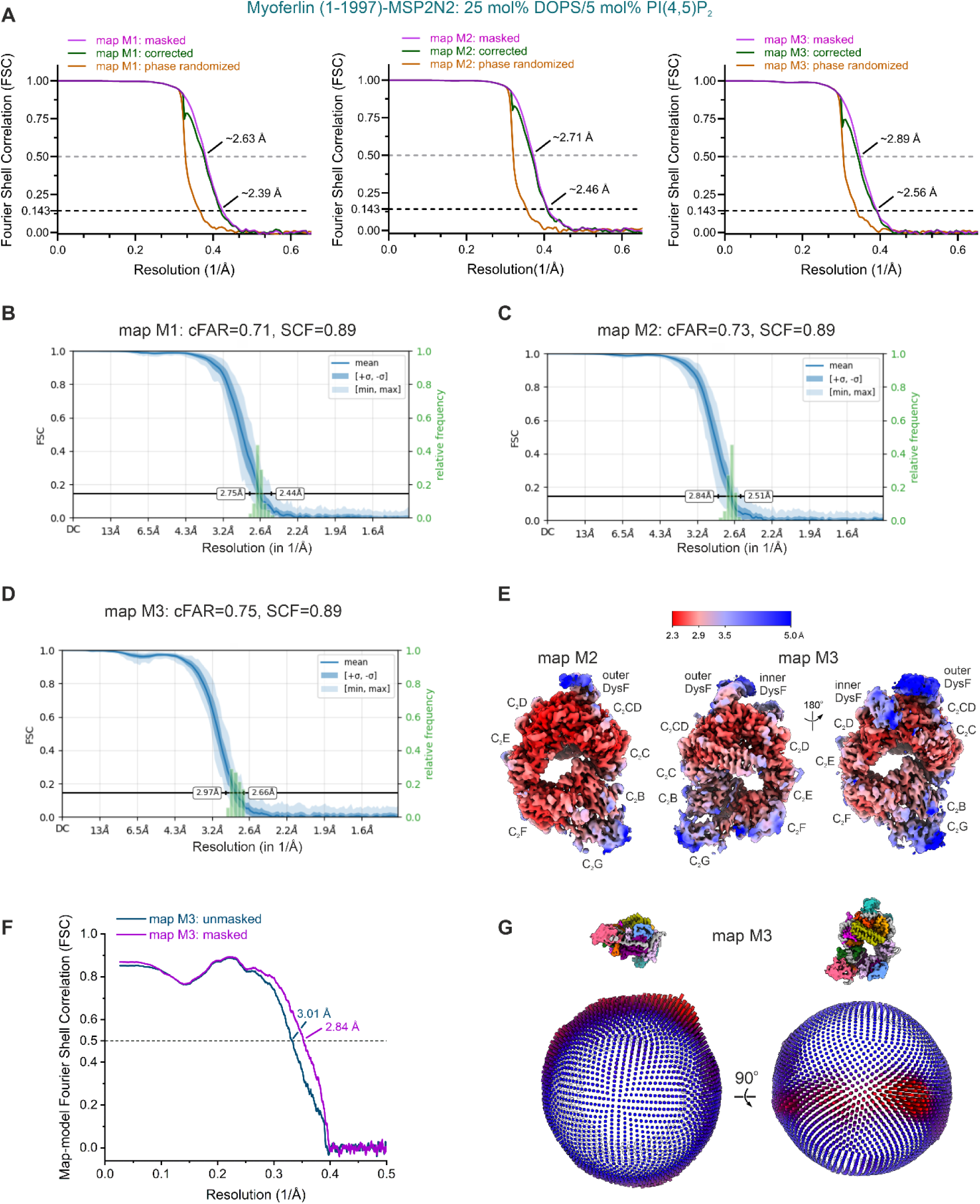
Resolution and quality of the lipid-bound myoferlin cryo-EM maps (25 mol% DOPS/5 mol% PI(4,5)P_2_ nanodisc). **A** Fourier shell correlation (FSC) between the final myoferlin (1-1997)-MSP2N2 cryo-EM half-maps (25 mol% DOPS/5 mol% PI(4,5)P_2_) datasets). The gold-standard FSC criterion (FSC=0.143) indicates global resolution estimates of ∼2.39 Å (consensus map), ∼2.46 Å (after masked C_2_G classification), and ∼2.56 Å (after masked DysF classification) for the M1, M2, and M3 maps, respectively. **B-D** Directional resolution and Fourier sampling of the myoferlin (1-1997)-MSP2N2 cryo-EM maps. The conical FSC, the conical FSC area ratio (cFAR), and the sampling compensation factor (SCF) were calculated and plotted with cryoSPARC v4.5. **E** Local resolution of the myoferlin (1-1997)-MSP2N2 cryo-EM maps (25 mol% DOPS/5 mol% PI(4,5)P_2_ nanodisc). Red color indicates a higher resolution. **F** Map versus model FSC plot for the myoferlin (1-1997)-MSP2N2 complex (map M3, 25 mol% DOPS/5 mol% PI(4,5)P_2_ datasets). **G** 3D angular distribution of the myoferlin particles contributing to the overall (M3) map (25 mol% DOPS/5 mol% PI(4,5)P_2_) datasets). The similarly oriented and color-coded maps are shown above the angular distribution plot. Red color and relative cylinder height indicates a higher number of particle images.

**Figure S4.**
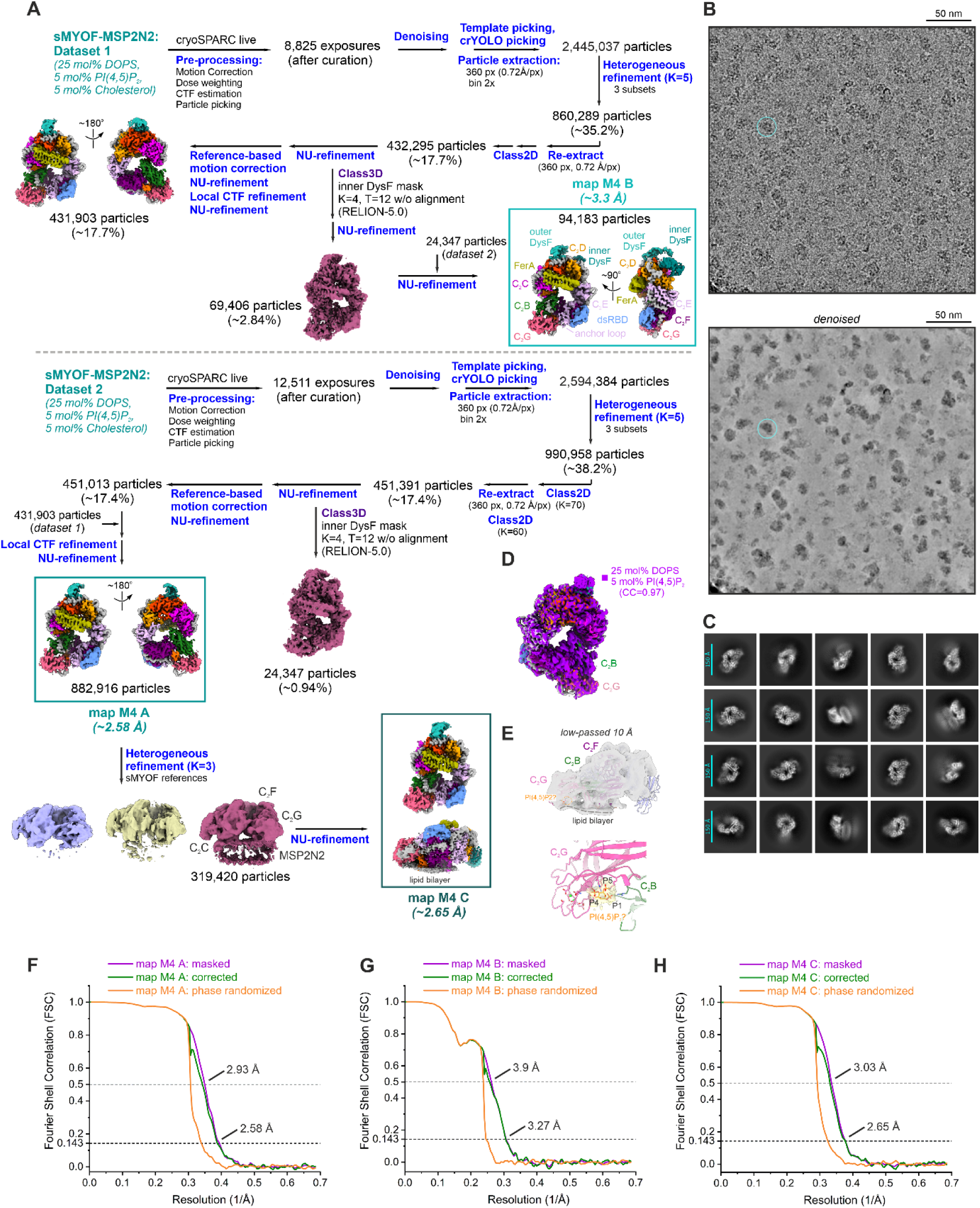
Cryo-EM image analysis of the vitrified myoferlin (1-1997)-MSP2N2 nanodisc complex (25 mol% DOPS/5 mol% PI(4,5)P_2_/5 mol% Cholesterol cryo-EM datasets). **A** Cryo-EM processing schematic for the soluble myoferlin (1-1997), bound to a 25 mol% DOPS/5 mol% PI(4,5)P_2_/5 mol% Cholesterol MSP2N2 nanodisc. The resolution estimates (FSC=0.143 criterion) of the final maps are indicated and the final maps are colored-coded. **B** Cryo-EM micrograph (raw and denoised) of the imaged myoferlin (1-1997)-MSP2N2 complex (comprising 25 mol% DOPS, 5 mol% PI(4,5)P_2_ and 5 mol% Cholesterol). A single vitrified particle is circled in cyan. **C** Reference-free 2D class averages of the myoferlin (1-1997)-MSP2N2 complex (25 mol% DOPS, 5 mol% PI(4,5)P_2_, and 5 mol% Cholesterol datasets). **D** Superposition between the consensus cryo-EM maps of nanodisc-bound myoferlin (1-1997), obtained in the absence (purple, 25 mol% DOPS/5 mol% PI(4,5)P_2_) or presence of Cholesterol (color-coded, 25 mol% DOPS/5 mol% PI(4,5)P_2_/5 mol% Cholesterol). **E** Low-passed cryo-EM map of the lipid-bound myoferlin (25 mol% DOPS/5 mol% PI(4,5)P_2_/5 mol% Cholesterol nanodisc, map M4 C) showing ordered lipid headgroups, including a tentative PI(4,5)P_2_, located close to the interface between C_2_F, C_2_G and C_2_B. **F-H** Fourier shell correlation (FSC) plots for the myoferlin (1-1997)-nanodisc maps (25 mol% DOPS/5 mol% PI(4,5)P_2_/5 mol% Cholesterol datasets).

**Figure S5.**
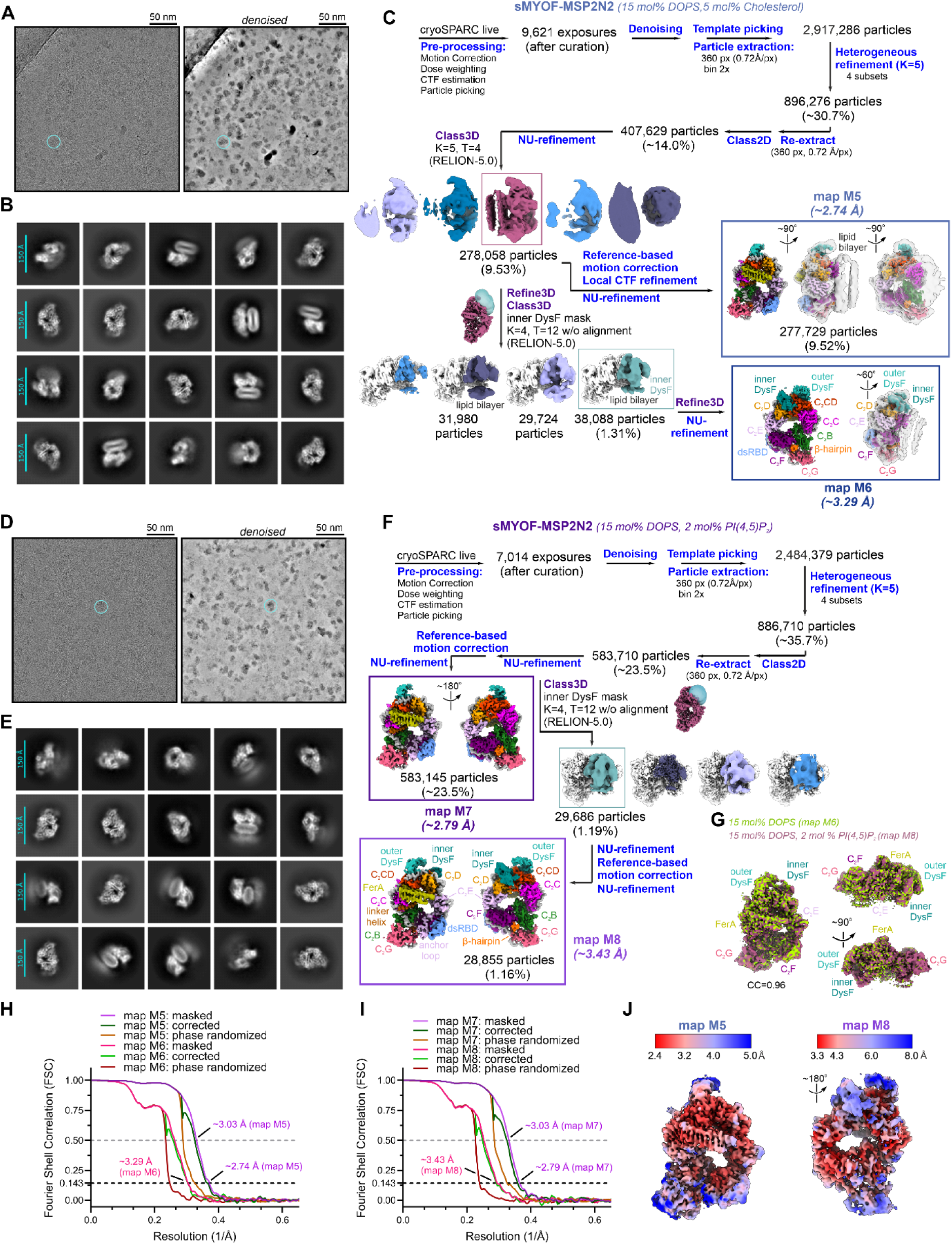
Cryo-EM analysis of the myoferlin (1-1997)-lipid complexes formed on 15 mol% DOPS/5 mol% Cholesterol and 15 mol%/2 mol% PI(4,5)P_2_ MSP2N2 nanodiscs. **A-B** Cryo-EM (raw and denoised) micrograph and 2D class averages of the myoferlin (1-1997)-MSP2N2 complex (comprising 15 mol% DOPS and 5 mol% Cholesterol). **C** Cryo-EM data processing flow-chart for the vitrified myoferlin (1-1997), bound to a 15 mol% DOPS/5 mol% Cholesterol nanodisc. The final maps are color-coded after the modeled domains and the resolution estimates (after the FSC=0.143 criterion) are indicated. **D-F** Cryo-EM image analysis of the soluble myoferlin (1-1997)-lipid complex (assembled onto a 15 mol% DOPS/2 mol% PI(4,5)P_2_ nanodisc). **G** Superposition between the M6 (15 mol% DOPS/5 mol% Cholesterol) and M8 (15 mol% DOPS/2 mol% PI(4,5)P_2_) nanodisc-bound myoferlin (1-1997) maps. The two cryo-EM maps of the complex are nearly identical (CC=0.96). **H-I** Fourier shell correlation (FSC) plots for the lipid-bound myoferlin (1-1997) maps (comprising 15 mol% DOPS/2 mol% PI(4,5)P2 and 15 mol% DOPS/5 mol% Cholesterol). **J** Local resolution of the myoferlin (1-1997)-MSP2N2 cryo-EM maps depicted in **C** and **F**. The map regions colored in red color indicate a higher resolution.

**Figure S6.**
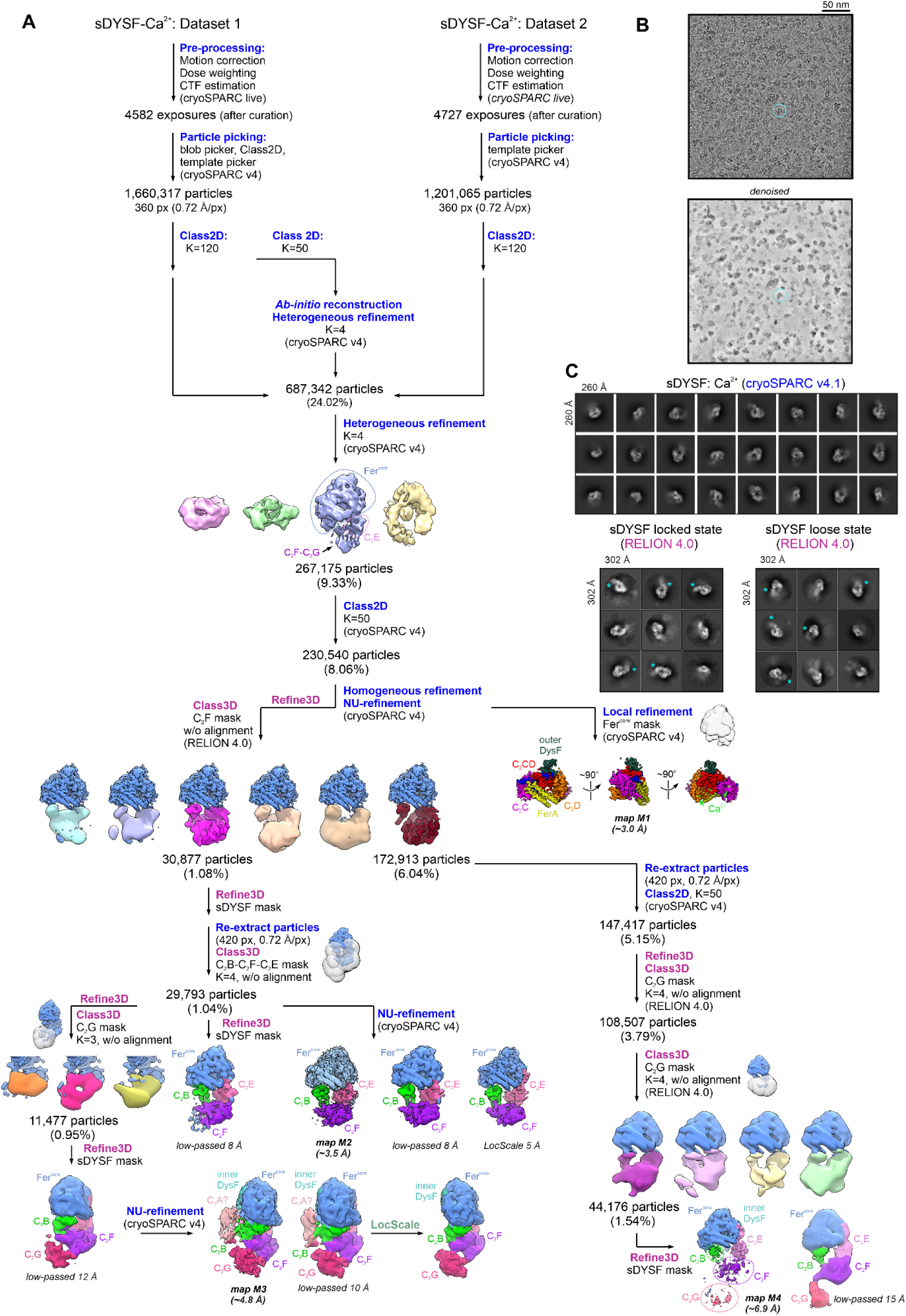
Computational image analysis of the lipid-free soluble dysferlin (1-2017) cryo-EM datasets. **A** Cryo-EM data processing routine of the Ca^2+^-bound soluble dysferlin (sDYSF, residues 1-2017). Resolution of the final maps (M1-M4) was estimated according to the gold-standard Fourier Shell Correlation (FSC) cutoff of 0.143. The key structural domains of dysferlin (1-2017) are color-coded and indicated in the final maps. **B** Typical cryo-EM micrograph (raw and denoised) of the vitrified lipid-free soluble dysferlin (1-2017). An imaged dysferlin particle is indicated in cyan. **C** Reference-free 2D class averages of lipid-free dysferlin (1-2017). 2D class averages calculated from the locked and loose dysferlin particle images are shown in the bottom panel. The C_2_F domain is indicated with an asterisk. The C-terminal domain is more ordered in the locked (closed) dysferlin 2D classes (left panel) compared to the loose state (right panel).

**Figure S7.**
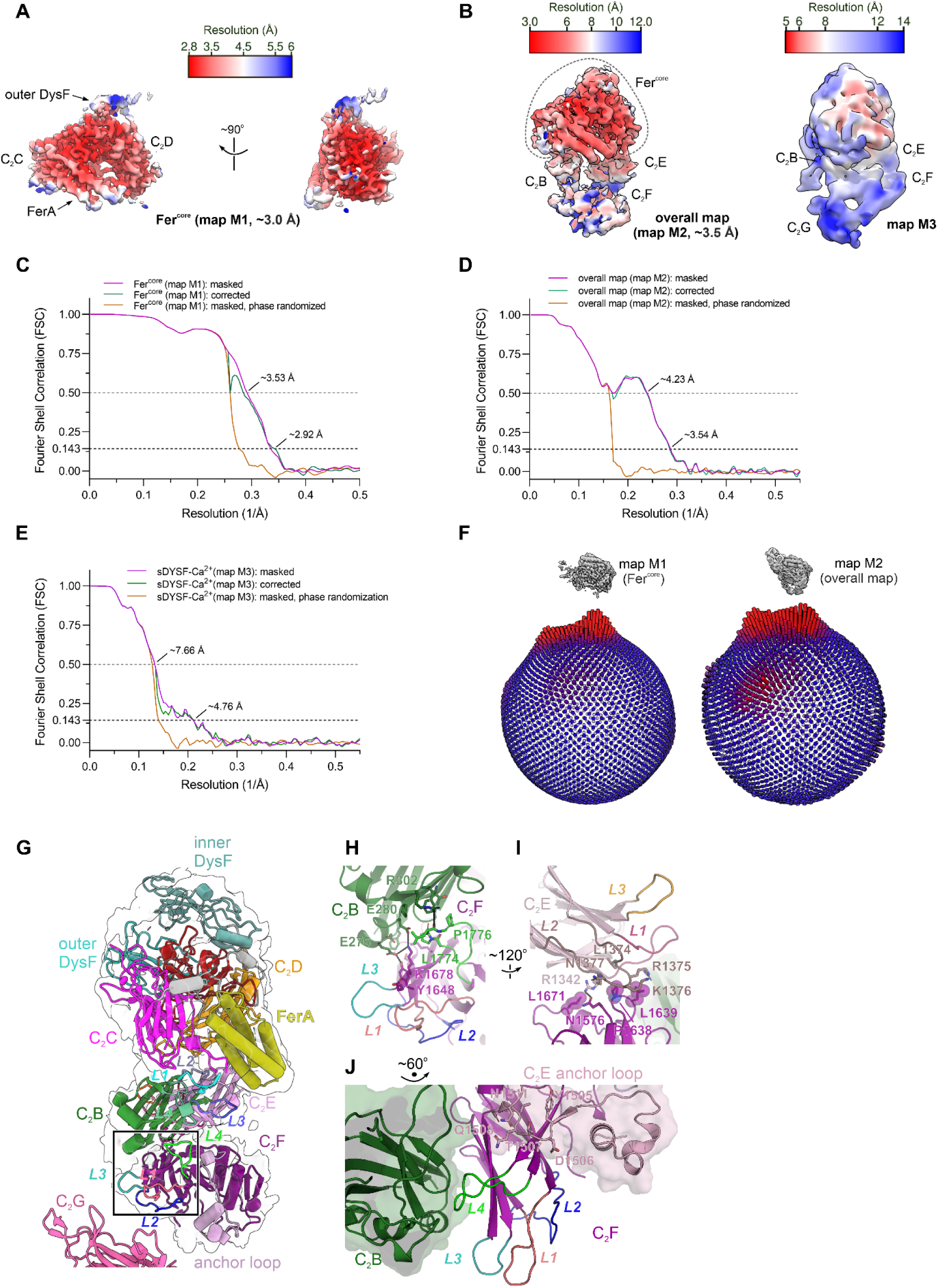
Resolution and quality of the lipid-free dysferlin (1-2017) cryo-EM maps. **A** Local resolution of the lipid-free Fer^core^ map (map M1) of dysferlin (1-2017). Red colored map regions indicate a higher resolution. **B** Overall maps of Ca^2+^-bound soluble dysferlin (map M2 and M3), colored after their local resolution. Note the higher local resolution of the Fer^core^ (C_2_C-C_2_CD-FerA-C_2_D) compared to the more dynamic C_2_B-C_2_F-C_2_E. **C-E.** Fourier shell correlation (FSC) between the final lipid-free dysferlin (1-2017) half maps. The gold-standard FSC criterion (FSC=0.143) indicates global resolution estimates of ∼2.9 Å, ∼3.5 Å, and ∼4.8 Å for the M1, M2 and M3 maps, respectively. **F** 3D angular distribution of the dysferlin particles contributing to the core (M1) and overall (M2) maps. The similarly oriented maps are shown above the angular distribution representation. Red color and relative cylinder height indicate a higher number of particle images. **G** Tertiary interfaces between C_2_F, C_2_B, and C_2_E observed in the dysferlin (1-2017) cryo-EM model. In the lipid-free state, the C_2_F domain is positioned at the base of the dysferlin ring by multiple tertiary interactions with the N-terminal C_2_B and C-terminal C_2_E. The final model of the soluble dysferlin (1-2017) is represented as cartoons and fitted inside the overall cryo-EM map (transparent surface). The L1-L4 loops of C_2_F are colored pink, blue, teal, and green, respectively. **H** Selected polar interactions between C_2_F’s L3-L4 loops and C_2_B. The polar contacts between interacting residues are depicted as dashed lines. **I** Polar and hydrophobic interactions between the bottom loops of C_2_F and the L2 loop of C_2_E. Residues involved in hydrophobic interactions are shown as transparent spheres. **J** Contact interfaces between C_2_F and the anchor loop of C_2_E (residues 1475-1514). C_2_B and the anchor loop are shown as transparent surfaces.

**Figure S8.**
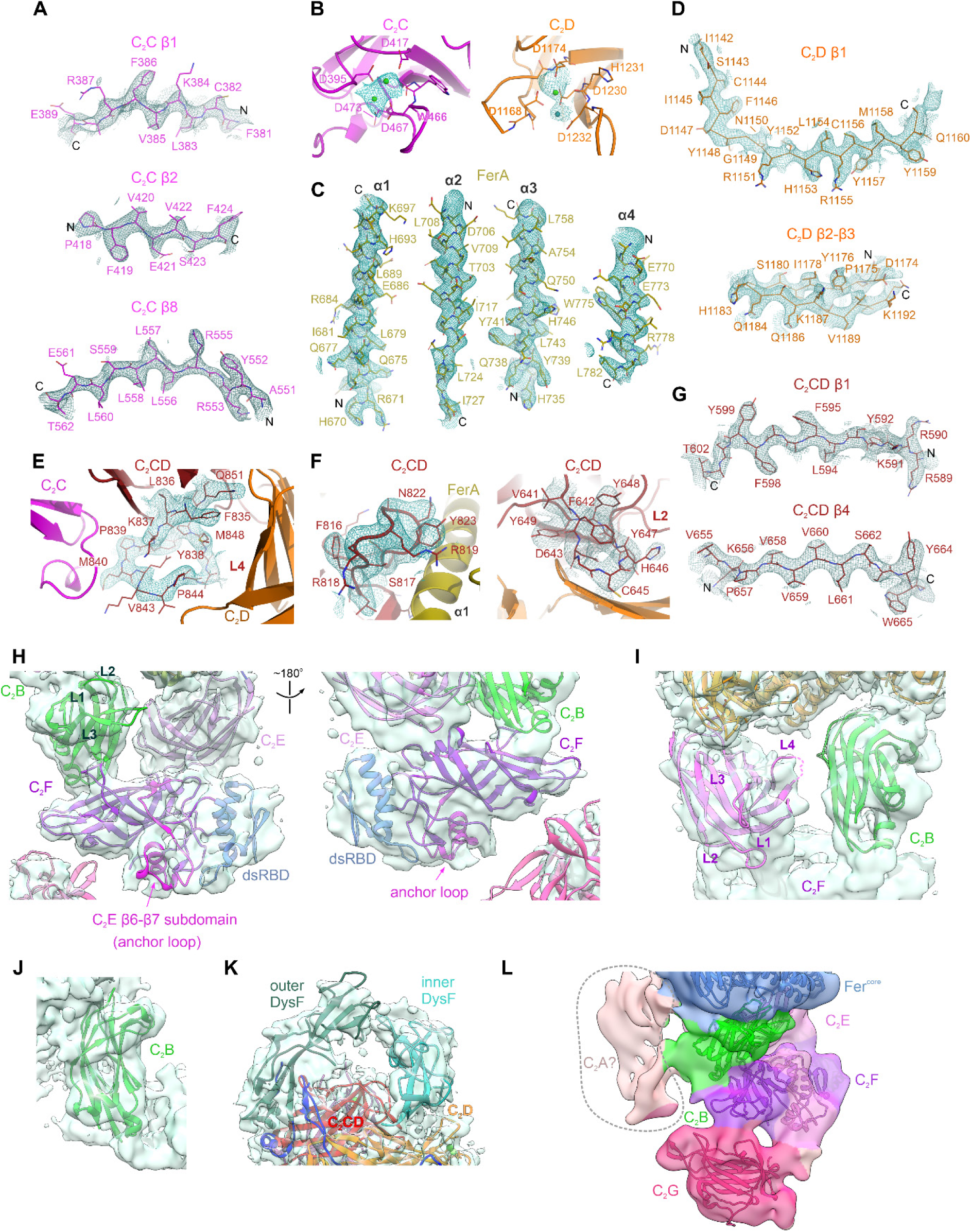
Snapshots of the lipid-free dysferlin (1-2017) cryo-EM density. **A** Selected snapshots of C_2_C’s cryo-EM density. C_2_C’s β-strands as shown as sticks. **B** Ca^2+^-binding sites of dysferlin’s C_2_C and C_2_D domains. The cryo-EM density map is colored in cyan and contoured around each of the two Ca^2+^-binding sites. **C** Cryo-EM density of the FerA domain helices. The FerA α-helices are amphipathic and form a typical, up-down-up-down four-helix bundle. **D** Cryo-EM densities of the C_2_D’s β-strands. **E** Cryo-EM density of C_2_CD’s L4 loop that bridges the C_2_C and C_2_D domains. **F** Cryo-EM densities of C_2_CD’s β6-β7 and L2 loops, contacting the FerA and C_2_D, domains, respectively. **G** Densities of selected C_2_CD β-strands. **H-I** Modeling of C_2_B, C_2_E and C_2_F in the locked (closed) state cryo-EM map (map M2). Note that the top loops of C_2_B and C_2_E face opposite surfaces of the dysferlin plane, whereas the top loops of C_2_F adopt a lateral orientation and are located close to C_2_B. **J** Side-view of C_2_B’s density with the modeled domain fitted inside the cryo-EM map. **K** Cryo-EM densities of the two, outer and inner DysF motifs of dysferlin. The outer and inner DysF occupy a peripheric location, between C_2_CD and C_2_D, and share a dynamic contact interface (*i.e.*, not present in all imaged dysferlin particles). **L** Densities of the C-terminal C_2_F and C_2_G domains. The cryo-EM map (map M3) has been low-passed to 10 Å. The additional density element, connected to C_2_B, might correspond to the flexible, N-terminal C_2_A domain, which likely samples many conformational states in solution. The long linker between C_2_A and C_2_B is, likely, largely unstructured as it is susceptible to protease digestion in cells. Note that cryo-EM density for the linker helix is absent in the lipid-free dysferlin state.

**Figure S9.**
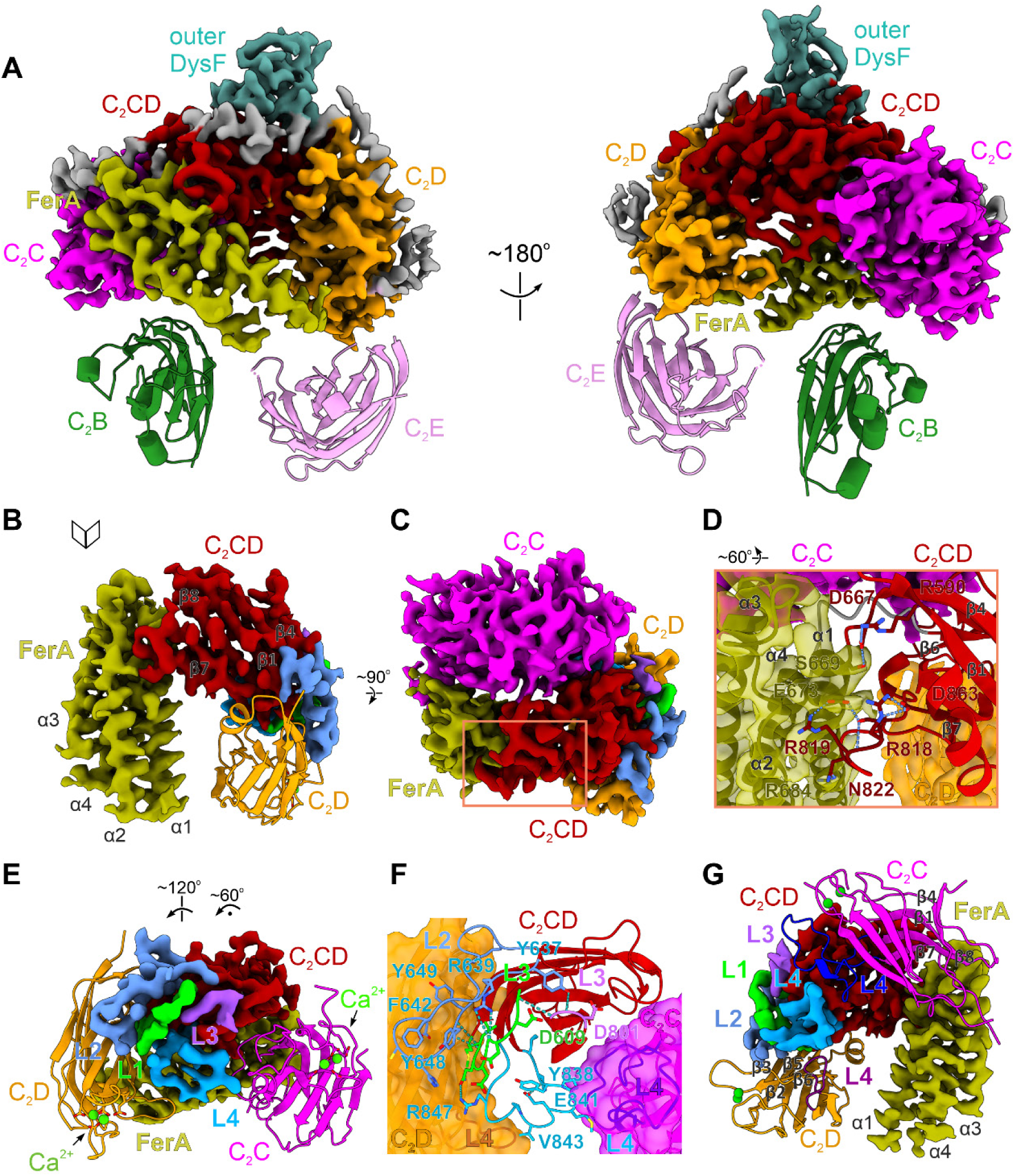
Structural organization of dysferlin’s Fer^core^ module. **A** Structure of dysferlin’s core module (Fer^core^) resolved at 2.9 Å resolution. The focused cryo-EM map of the C_2_C-C_2_D region (map M1), shown in two different orientations, has been sharpened and colored after the modelled subunits. **B-C** Interaction interfaces between C_2_CD and its neighboring FerA, C_2_C and C_2_D domains. The cryo-EM maps of the dysferlin domains have been sharpened. **D** An insertion in C_2_CD’s β6-β7 appears to stabilize the pose of the FerA domain. Dysferlin’s domains are depicted in cartoon representation and the sharpened cryo-EM map is contoured around FerA and C_2_D. The key polar interactions between C_2_CD’s β6-β7 loop and FerA are shows as dashed blue lines. **E** C_2_CD’s top loops are engaged in multiple contacts and bridge the C_2_C and C_2_D domains. The L1-L4 loops of C_2_CD are colored green, light blue, purple and blue, respectively. C_2_CD and FerA are depicted as cryo-EM maps. **F** Selected polar interactions between C_2_CD’s top loops (L1-L4) and the bridged C_2_C and C_2_D domains. The polar interactions are shown as dashed blue lines. C_2_C and C_2_D are depicted as surfaces. **G** The extensive contact interfaces between C_2_CD-FerA and the Ca^2+^-binding C_2_C and C_2_D. C_2_CD and FerA are shown as cryo-EM maps, whereas C_2_C and C_2_D are depicted as cartoons.

**Figure S10.**
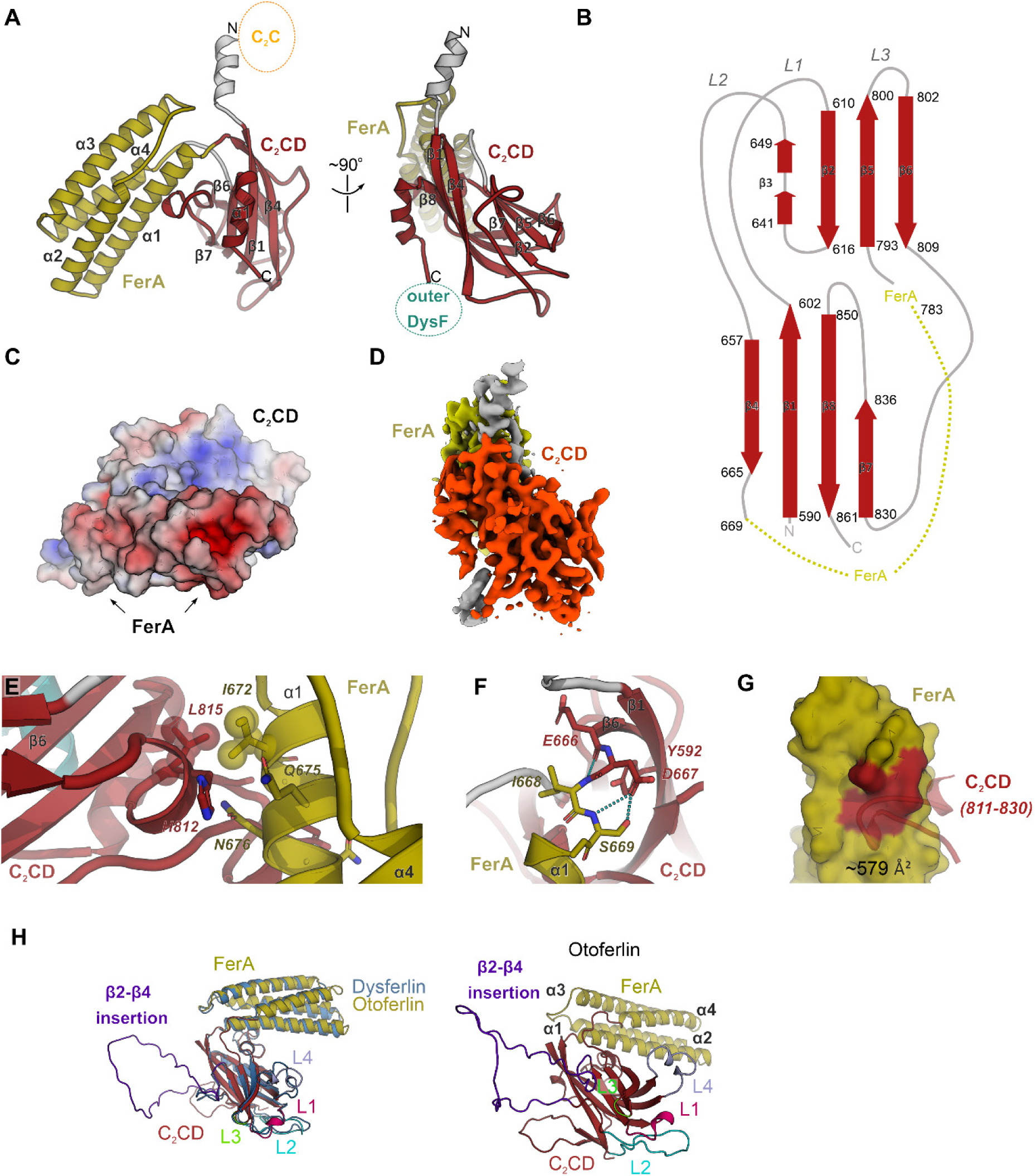
C_2_CD represents a new C_2_-like ferlin structural domain. **A** Structure of the C_2_CD-FerA module, using dysferlin as an example. Note that the FerA helical bundle inserts between C_2_CD’s β4-β5 strands and projects from the bottom side of the C_2_-like domain. **B** Topology diagram of the C_2_CD domain. As all the other ferlin C_2_ domains, C_2_CD has a type-II topology (*i.e.*, the two β-sheets are formed by β1-β4-β7-β8 and β2-β3-β5-β6, respectively). **C** Surface-charge distribution of the C_2_CD-FerA module of dysferlin. The electrostatic potentials were colored from red (negative, -3 kT/e) to blue (positive, +3 kT/e) and mapped onto the solvent excluded surface. **D** Cryo-EM density of dysferlin’s C_2_CD-FerA module oriented and colored as in **A**. **E-F** Selected intramolecular contacts between C_2_CD and FerA. The polar interactions are represented as dashed lines. **G** C_2_CD’s β6-β7 loop interacts with FerA’s bottom side. FerA’s α1 helix is shown as surface representation and the C_2_CD interacting residues are colored red. **H** A similarly organized C_2_CD-FerA module is likely present in all human ferlins. Alphafold2 prediction of otoferlin’s C_2_CD-FerA region (right) is depicted side-by-side with its structural superposition to the cryo-EM model of dysferlin’s C_2_CD-FerA (left). Note the predicted large insertion between C_2_CD’s β2-β4 strands observed in otoferlin.

**Figure S11.**
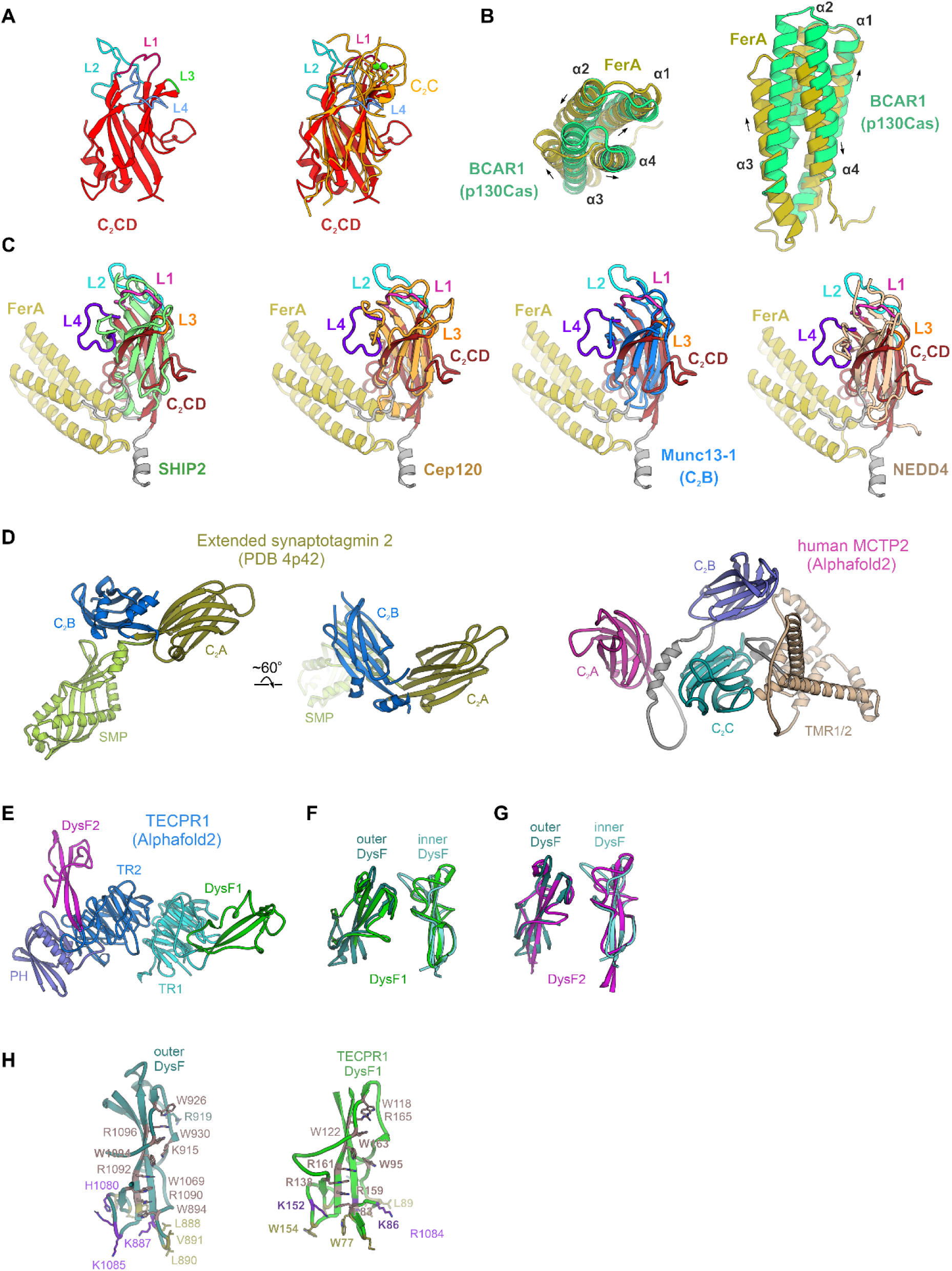
Structural conservation of ferlins’ C_2_CD, FerA and DysF motifs. **A** Structural superposition between C_2_CD and the Ca^2+^-bound C_2_C domain of dysferlin, as an exemplary type-I ferlin. C_2_C’s Ca^2+^ ions are shown as green spheres. **B** Structural similarity between dysferlin’s FerA motif and the four-helix bundle of BCAR1 (p130Cas, PDB 3t6g). The direction of the four α-helices is indicated by arrows. **C** Alignments between the C_2_CD of dysferlin and the structurally similar C_2_ domains of the SHIP2 phosphatase (PDB 5okm), Cep120 (PDB 6flj), Munc13-1 (PDB 5ue8) and NEDD4 (PDB 3m7f). The C_2_ domains were aligned using Foldseek (van Kempen, Kim et al., 2024). **D** Structural organization of extended synaptotagmin 2 (ESYT2) and human MCTP2 (Alphafold2 prediction). SMP – synaptotagmin*-*like mitochondrial lipid-binding protein; TMR1/2 – transmembrane region 1 and 2. **E** Alphafold2 prediction of TECPR1 (Tectonin Beta-Propeller Repeat-containing Protein 1)’s structure. The tectonin repeat β-propellers are colored blue and cyan respectively (TR1, TR2). The two spatially separated DysF motifs of TECPR1 are colored in green (DysF1) and magenta (DysF2), respectively. **F** Superposition between dysferlin’s outer and inner DysF motifs, as modelled in the cryo-EM map, and the related DysF1 domain of TECPR1, predicted by Alphafold2 (Jumper et al., 2021). **G** Structural superposition between the DysF motifs of dysferlin and TECPR1’s DysF1. **H** Stabilizing arginine/tryptophan (R/W) stacking interactions are present in both dysferlin/myoferlin’s DysF and TECPR1’s DysF motifs. In both cases, the R/W stacks (shown as sticks) are mainly observed on one side of the DysF β-sheet (Patel, Harris et al., 2008, Sula, Cole et al., 2014).

**Figure S12.**
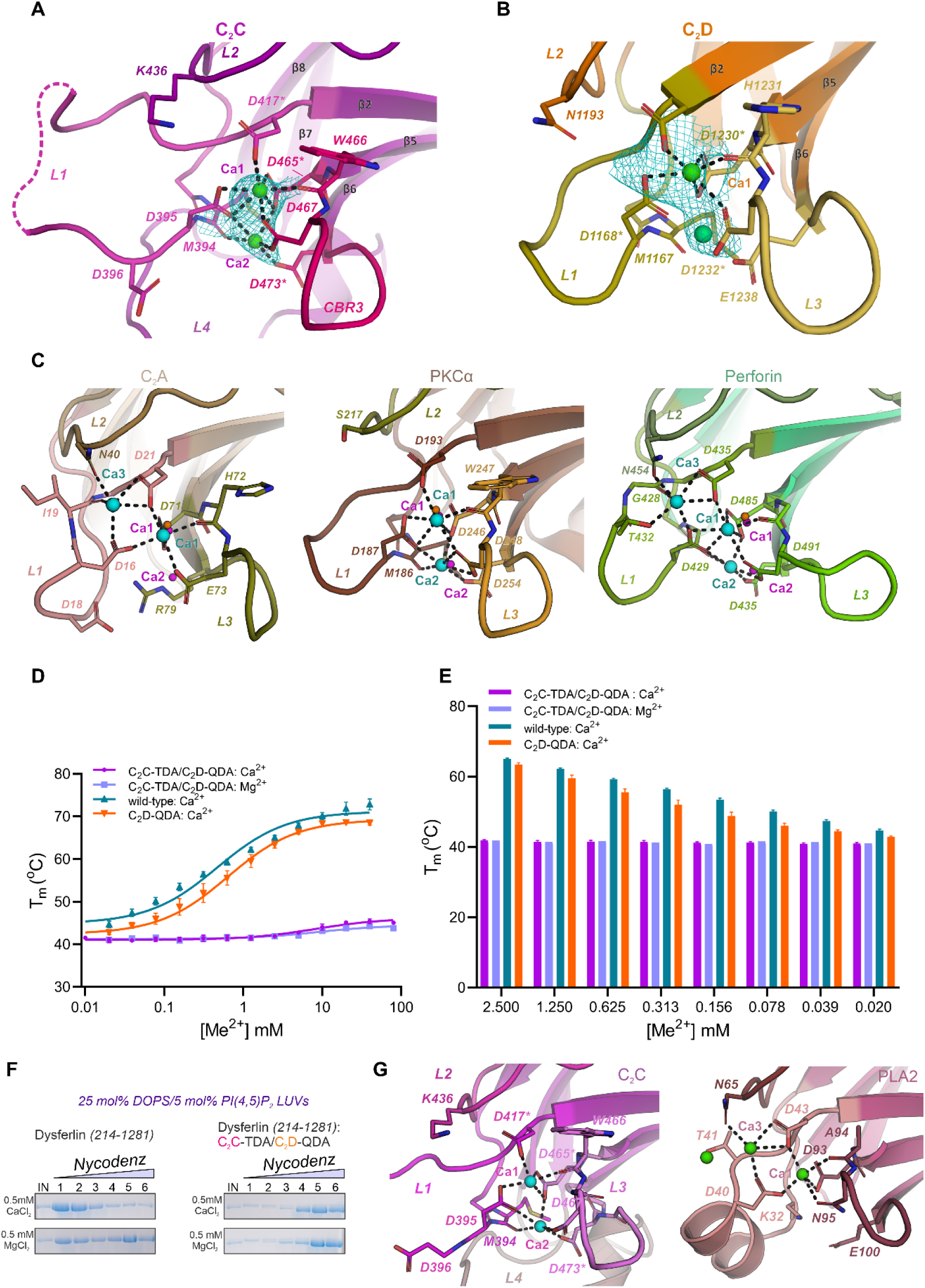
Ca^2+^-binding sites of the Fer^core^ module of dysferlin, modeled in the lipid-free state. **A** In dysferlin, two Ca^2+^ ions bind C_2_C’s Ca^2+^-binding pocket. The cryo-EM map (cyan mesh) is contoured around the Ca^2+^ ions (green spheres). Residues substituted to alanine (in **D-E**) are marked with asterisks. **B** The Ca^2+^-binding sites of the C_2_D domain. Residues substituted for alanine in **D-E** are indicated with asterisks. **C** Comparison between Ca^2+^-coordination by dysferlin’s C_2_C and C_2_A (PDB 7jof), and the C_2_-domains of PKCα (PDB 1dsy) and perforin (PDB 4y1t). C_2_C’s Ca^2+^ ions are shown as magenta spheres, whereas the Ca^2+^ sites of C_2_A, PKCα and perforin are colored in cyan. **D** Ca^2+^-binding activities of the Fer^core^ of dysferlin (residues 214-1281) and of its C_2_D-QDA (“quadruple D (aspartate) to A (alanine)”) and C_2_C-TDA (“triple D (aspartate) to A (alanine)”)/C_2_D-QDA mutants measured using a nanoDSF assay. Four Ca^2+^-binding aspartates were substituted in C_2_D-QDA (D1168A, D1174A, D1230A, D1232A). Three additional residues were substituted to alanine (D417A, D465A, D473A) in the C_2_C-TDA/C_2_D-QDA variant (see also **Appendix Fig S13A-C**). The Ca^2+^-activity measurements were performed in triplicates (n=3) for the C_2_D-QDA and C_2_C-TDA/C_2_D-QDA samples and repeated five times for the wild-type Fer^core^ sample (n=5). Error bars represent the standard deviation (s.d.). The melting temperatures (T_m_) of the analyzed samples were plotted as a function of Ca^2+^/Mg^2+^ concentration and the [Ca^2+^]_1/2_ values were estimated by nonlinear regression curve fitting. **E** Bar plot of the Ca^2+^-activity data shown in **D**. Error bars represent the s.d.. **F** Ca^2+^-sensitive liposome binding activity of the wild-type and C_2_C-TDA/C_2_D-QDA Fer^core^ of dysferlin (residues 214-1281), assessed using coflotation assays. The liposomes (LUVs) comprised 25 mol% DOPS and 5 mol% PI(4,5)P_2_. The Nycodenz step gradients were harvested from the top and analyzed by SDS-PAGE. Note that substitution of C_2_C’s and C_2_D’s Ca^2+^-binding sites reduced significantly the ability of the Fer^core^ module to interact with lipid membranes comprising acidic phospholipids. **G** Modeled Ca^2+^-binding sites in dysferlin’s C_2_C and the C_2_ domain of phospholipase A2 (PLA2, PDB 6eij). The Ca^2+^ ions are depicted as spheres and the coordinating side-chains are shown as sticks.

**Figure S13.**
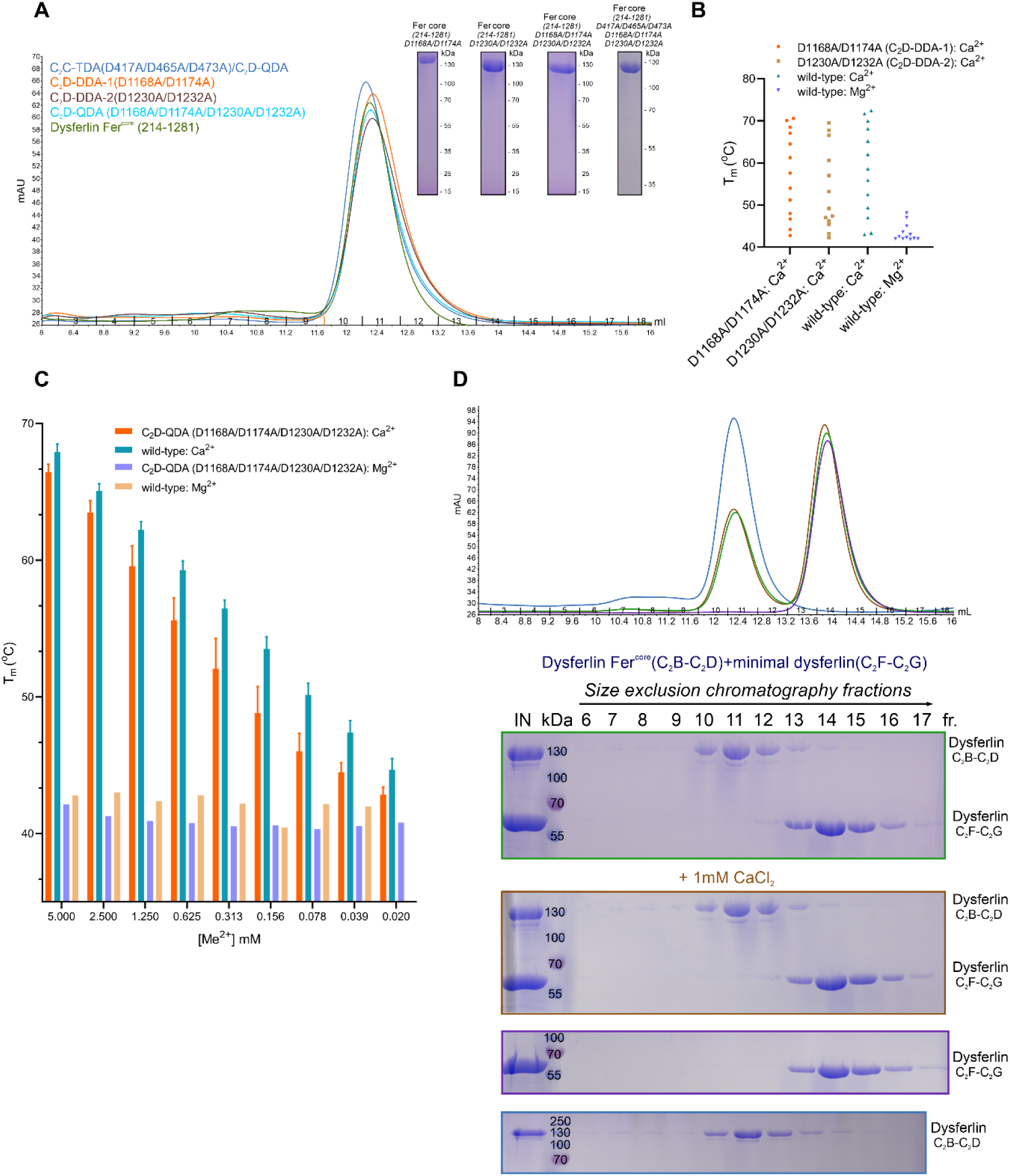
Biochemical validation of dysferlin’s Ca^2+^-binding sites. **A** Size-exclusion chromatography profiles of C_2_C and C_2_D Ca^2+^-binding mutants. The purified mutants have similar retention volumes on Superdex 200, comparable to the wild-type Fer^core^ sample (dysferlin (214-1281)). Therefore, substitution of two (C_2_D-DDA-1: D1168A/D1174A and C_2_D-DDA2:D1230A/D1232A) or four Ca^2+^-binding aspartates (C_2_D-QDA: D1168A/D1174A/D1230A/D1232A) in C_2_D does not have a significant impact on Fer^core^’s structure (*i.e.*, does not destabilize its overall fold). Consistently, combined substitutions in C_2_C (C_2_C-TDA: D417A/D465A/D473A) and C_2_D (C_2_D-QDA) preserve the structural integrity of the Fer^core^ of dysferlin while abolishing its Ca^2+^-binding activity (see also **Appendix Fig S12D-E**). SDS-PAGE gels of the Fer^core^ dysferlin mutants carrying substitutions in C_2_C’s and C_2_D’s Ca^2+^-binding loops are shown as an inset. Three aspartate to alanine substitutions were introduced in C_2_C (D417A, D465A, D473A) and two or four substitutions were introduced in C_2_D (D1168A/D1174A, D1230A/D1232A and D1168A/D1174A/D1230A/D1232A). **B** Ca^2+^-binding activity of C_2_D’s DDA (“double D (aspartate) to A (alanine)”) substitutions measured using a nanoDSF assay. The melting temperatures (T_m_) of C_2_D-DDA-1 (D1168A/D1174A) and C_2_D-DDA-2 (D1230A/D1232A) were measured as a function of [Ca^2+^] or [Mg^2+^]. Single representative nanoDSF T_m_ estimates are shown (n=1). **C** Ca^2+^-activity measurements of the C_2_D-QDA (D1168A/D1174A/D1230A/D1232A) Fer^core^ mutant. The estimated T_m_ values were plotted as a function of [Ca^2+^] or [Mg^2+^]. Error bars represent the s.d. (C_2_D-QDA: n=3, wild-type: n=5). **D** Probing the interaction between C_2_B-C_2_D and C_2_F-C_2_G dysferlin fragments by size-exclusion chromatography (SEC). The purified samples were mixed in *trans*, in the presence or absence of 1 mM Ca^2+^, subjected to SEC on the Superdex 200 column and the peak fractions analyzed by SDS-PAGE. In these experiments, no complex formation between C_2_B-C_2_D (Fer^core^, residues 214-1281) and C_2_F-C_2_G (*min*DYSF, residues 1503-1560 and 1613-2056) was observed (C_2_B-C_2_D peaked in fraction 11, whereas the C_2_F-C_2_G fragment in fraction 14). The chromatograms of the reconstitution experiments are shown on the top.

**Figure S14.**
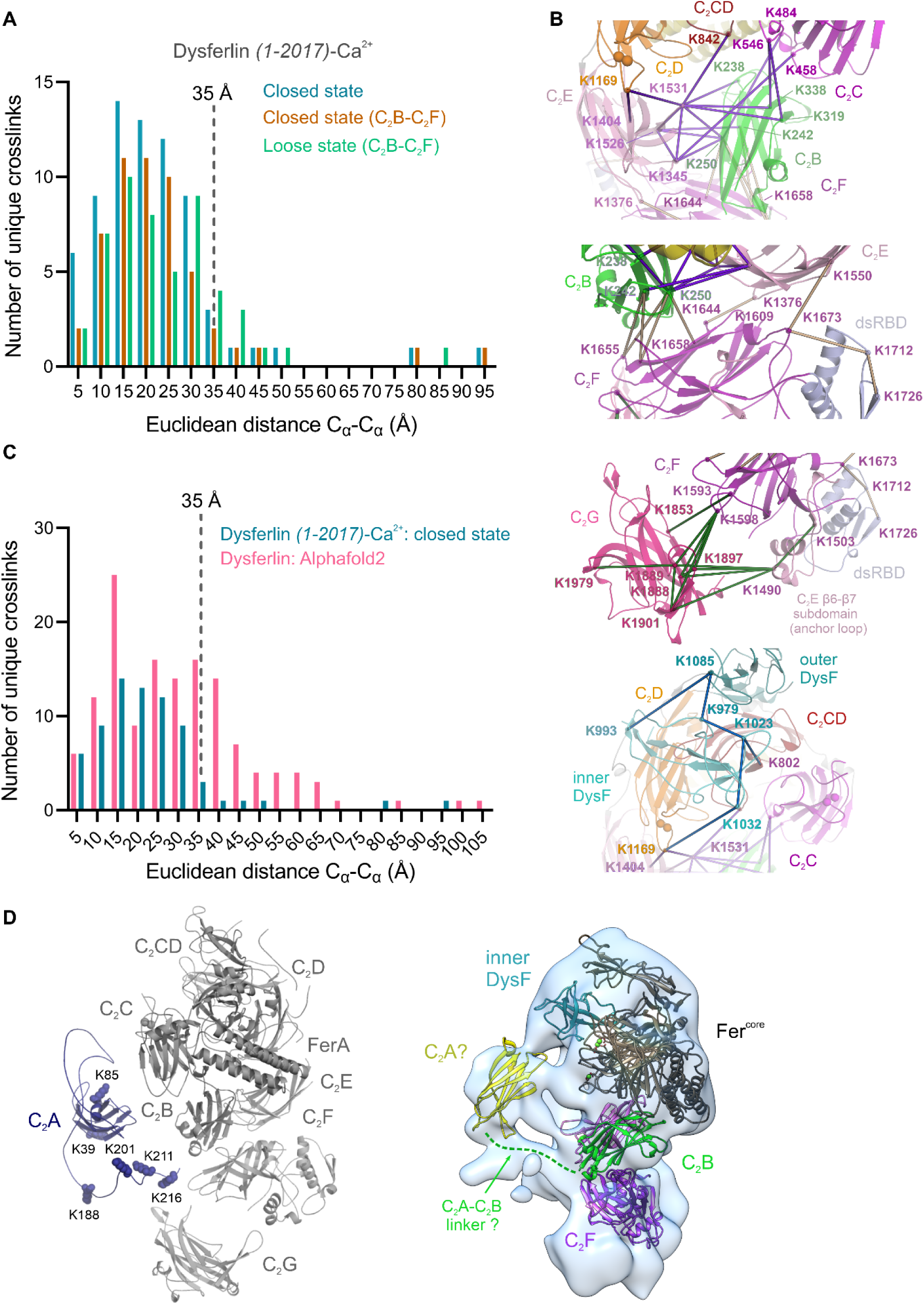
Chemical-crosslinking mass spectrometry analysis of the Ca^2+^-bound soluble dysferlin (1-2017). **A** Distribution of the Cα-Cα Euclidean distances between crosslinked lysine residues in the locked and loose dysferlin states. Dysferlin’s chemical crosslinking data indicate that, when mapped onto the locked/closed state model, ∼93% of unique crosslinks are observed between lysine residues less than 35 Å apart. Mapping to a loose state model lacking the more dynamic C_2_G, results in ∼85% crosslinks satisfying the distance threshold, showing that the dysferlin conformations could co-exist in the presence of Ca^2+^. **B** Selected chemical crosslinks between DYSF’s structural motifs. The Euclidean distances were calculated between the Cα atoms of the crosslinked lysines in the locked/closed dysferlin state and depicted as colored lines. **C** Distribution of the crosslink distances in the locked/closed state of soluble dysferlin (residues 1-2017, sDYSF) and a full-length dysferlin Alpahfold2 prediction (model 1, **Appendix Fig S15B**). Note that compared to the locked/closed state model, only ∼64.9% (37/57) unique crosslinks occur between residues less than 35 Å apart. The crosslink outliers map to the C_2_B-C_2_F, C_2_F-C_2_G and C_2_A-C_2_B interfaces, indicating that the relative orientation of these domains is not accurately predicted in the Alphafold2 model. **D** BS3 crosslinks involving the flexible C_2_A and the C_2_A-C_2_B linker. The crosslinking data indicate that, although generally flexible, C_2_A, likely, samples a restricted number of conformational states in the proximity of the inner DysF and C_2_B. Some of the poses of C_2_A could be accommodated by the unassigned density region which continues from C_2_B towards the center of dysferlin’s (and myoferlin’s) membrane-binding surface.

**Fig. S15.**
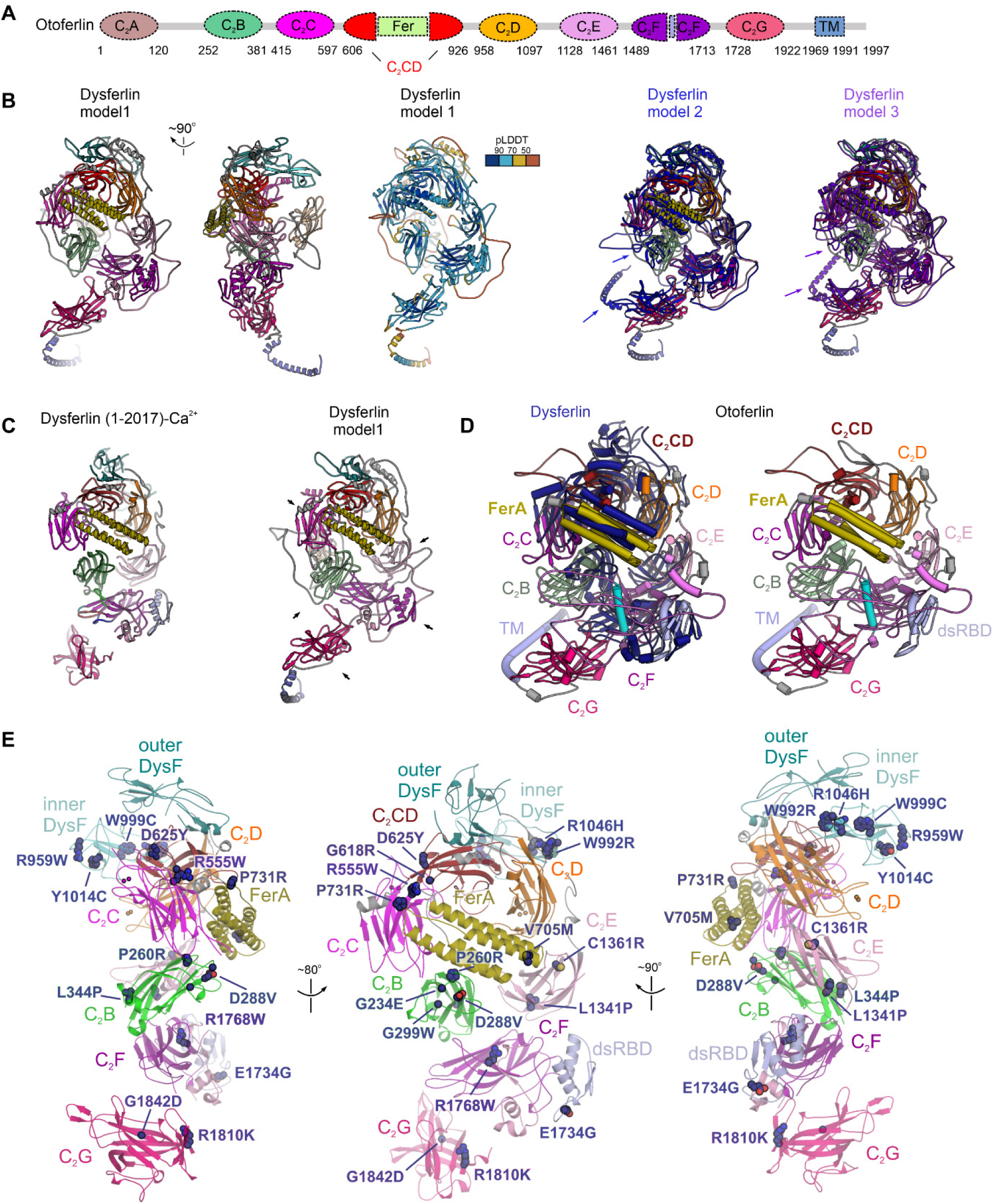
Common organization principles of human ferlins. **A** Alphafold2-based domain composition schematic of human otoferlin (Jumper, Evans et al., 2021). **B** Alphafold2 models of human dysferlin. While the Fer^core^ module is similar in these predictions, C_2_B, C_2_E, C_2_F, C_2_G and TM adopt alternative poses (indicated with arrows), consistent with them being structurally dynamic. The long linkers connecting the C_2_A-C_2_B and C_2_E-C_2_F are likely unstructured, having pLDDT (predicted local distance difference test) scores <50. **C** Comparison between soluble dysferlin’s locked (closed) state and an Alphafold2 prediction (model 1) of full-length dysferlin. Note the significant different poses of the C_2_B, C_2_E, C_2_F, and C_2_G domains (marked with arrows, r.m.s.d.=4.85 over 1156 atoms). Minor differences are observed for the FerA domain. The Fer^core^ module (except the inner DysF motif) is similarly organized in the soluble dysferlin (residues 1-2017, sDYSF) structure and Alphafold2 models (r.m.s.d.=1.79 over 618 atoms). **D** Comparison between soluble dysferlin’s cryo-EM model and the Alphafold2 prediction of human otoferlin. Note that otoferlin lacks the DysF motifs and the C_2_E anchor loop (magenta) is more extended. In addition, C_2_B, C_2_E and C_2_F are arranged in a near circular conformation, compared to the more elliptical appearance of dysferlin’s ring – a configuration which appears reminiscent of the lipid-bound myoferlin state. **E** Highly pathogenic patient mutations mapped onto dysferlin’s structure in the lipid-free state. Pathogenic and likely pathogenic dysferlin substitutions were extracted from the ClinVar database (Landrum, Chitipiralla et al., 2020) and manually curated. Note that some dysferlin substitutions appear to cluster at tertiary C_2_ domain interfaces, such as between C_2_C and C_2_CD (*e.g*., R555W and D625Y).

References

---

**Table S1.**
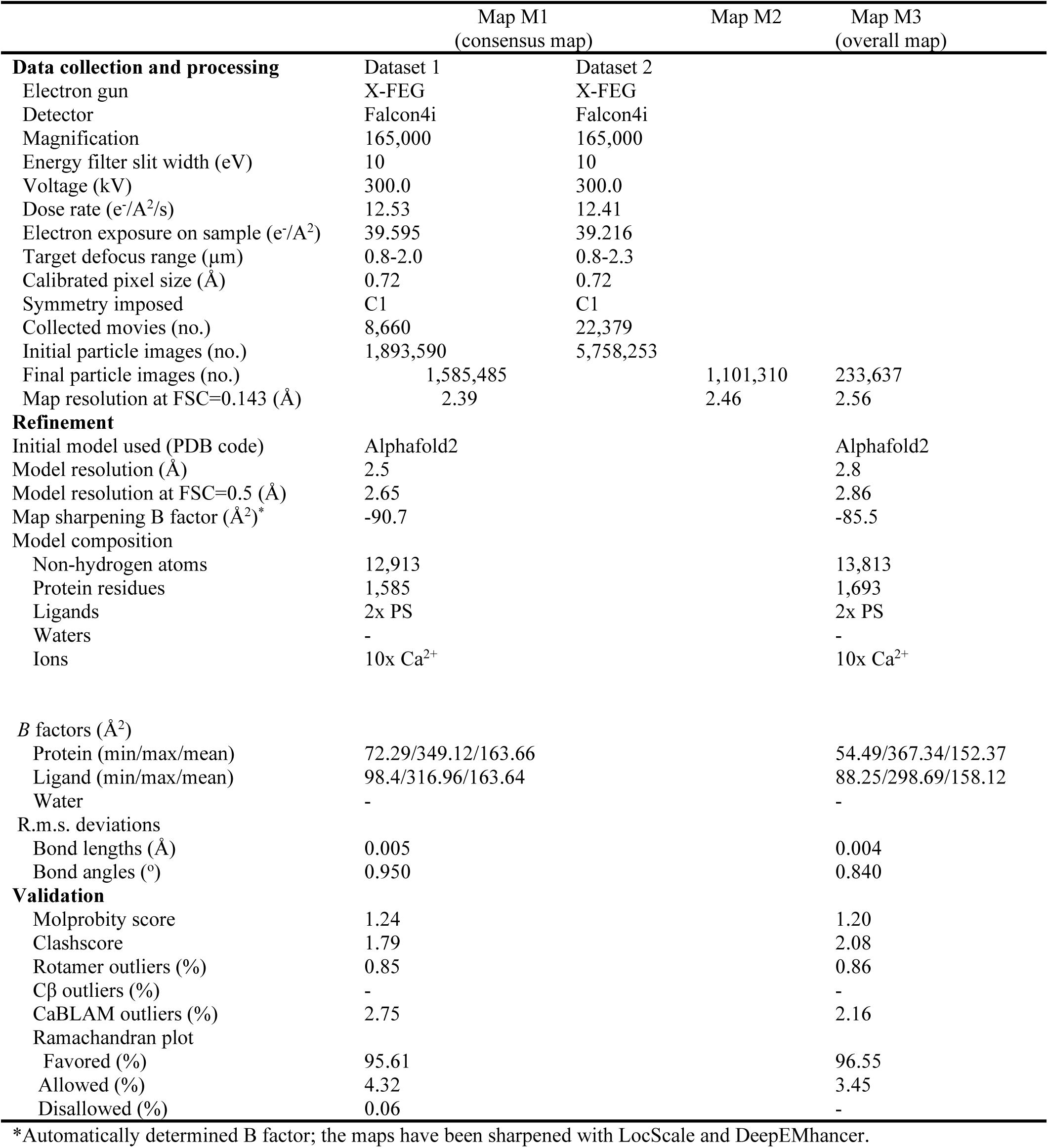
Data collection, refinement, and validation statistics for the lipid-bound soluble myoferlin (1-1997): 25 mol% DOPS/5 mol% PI(4,5)P_2_ lipid nanodisc complex.

|  | Map M1<br>(consensus map) | Map M2 | Map M3<br>(overall map) |
| --- | --- | --- | --- |
| **Data collection and processing** | Dataset 1 | Dataset 2 |  |
| Electron gun | X-FEG | X-FEG |  |
| Detector | Falcon4i | Falcon4i |  |
| Magnification | 165,000 | 165,000 |  |
| Energy filter slit width (eV) | 10 | 10 |  |
| Voltage (kV) | 300.0 | 300.0 |  |
| Dose rate (e<sup>-</sup>/Å<sup>2</sup>/s) | 12.53 | 12.41 |  |
| Electron exposure on sample (e<sup>-</sup>/Å<sup>2</sup>) | 39.595 | 39.216 |  |
| Target defocus range (μm) | 0.8-2.0 | 0.8-2.3 |  |
| Calibrated pixel size (Å) | 0.72 | 0.72 |  |
| Symmetry imposed | C1 | C1 |  |
| Collected movies (no.) | 8,660 | 22,379 |  |
| Initial particle images (no.) | 1,893,590 | 5,758,253 |  |
| Final particle images (no.) | 1,585,485 | 1,101,310 | 233,637 |
| Map resolution at FSC=0.143 (Å) | 2.39 | 2.46 | 2.56 |
| **Refinement** |  |  |  |
| Initial model used (PDB code) | AlphaFold2 |  | AlphaFold2 |
| Model resolution (Å) | 2.5 |  | 2.8 |
| Model resolution at FSC=0.5 (Å) | 2.65 |  | 2.86 |
| Map sharpening B factor (Å<sup>2</sup>)* | -90.7 |  | -85.5 |
| Model composition |  |  |  |
| Non-hydrogen atoms | 12,913 |  | 13,813 |
| Protein residues | 1,585 |  | 1,693 |
| Ligands | 2x PS |  | 2x PS |
| Waters | - |  | - |
| Ions | 10x Ca<sup>2+</sup> |  | 10x Ca<sup>2+</sup> |
| *B* factors (Å<sup>2</sup>) |  |  |  |
| Protein (min/max/mean) | 72.29/349.12/163.66 |  | 54.49/367.34/152.37 |
| Ligand (min/max/mean) | 98.4/316.96/163.64 |  | 88.25/298.69/158.12 |
| Water | - |  | - |
| R.m.s. deviations |  |  |  |
| Bond lengths (Å) | 0.005 |  | 0.004 |
| Bond angles (°) | 0.950 |  | 0.840 |
| **Validation** |  |  |  |
| Molprobity score | 1.24 |  | 1.20 |
| Clashscore | 1.79 |  | 2.08 |
| Rotamer outliers (%) | 0.85 |  | 0.86 |
| Cβ outliers (%) | - |  | - |
| CaBLAM outliers (%) | 2.75 |  | 2.16 |
| Ramachandran plot |  |  |  |
| Favored (%) | 95.61 |  | 96.55 |
| Allowed (%) | 4.32 |  | 3.45 |
| Disallowed (%) | 0.06 |  | - |

\*Automatically determined B factor; the maps have been sharpened with LocScale and DeepEMhancer.

**Table S2.** Data collection, refinement, and validation statistics for the lipid-bound soluble myoferlin (1-1997): 25 mol% DOPS/5 mol% Cholesterol/5 mol% PI(4,5)P_2_ lipid nanodisc complex.

|  | Map M4 A<br>(consensus map) | Map M4 C | Map M4 B<br>(overall map) |
| --- | --- | --- | --- |
| **Data collection and processing** | Dataset 1 | Dataset 2 |  |
| Electron gun | X-FEG | X-FEG |  |
| Detector | Falcon4i | Falcon4i |  |
| Magnification | 165,000 | 165,000 |  |
| Energy filter slit width (eV) | 10 | 10 |  |
| Voltage (kV) | 300.0 | 300.0 |  |
| Dose rate (e<sup>-</sup>/Å<sup>2</sup>/s) | 10.99 | 15.82 |  |
| Electron exposure on sample (e<sup>-</sup>/Å<sup>2</sup>) | 38.025 | 38.917 |  |
| Target defocus range (μm) | 0.8-2.0 | 0.8-2.0 |  |
| Calibrated pixel size (Å) | 0.72 | 0.72 |  |
| Symmetry imposed | C1 | C1 |  |
| Collected movies (no.) | 8,825 | 12,511 |  |
| Initial particle images (no.) | 2,445,037 | 2,594,384 |  |
| Final particle images (no.) | 882,916 | 319,420 | 94,183 |
| Map resolution at FSC=0.143 (Å) | 2.58 | 2.65 | 3.27 |
| **Refinement** |  |  |  |
| Initial model used (PDB code) | AlphaFold2 | AlphaFold2 | AlphaFold2 |
| Model resolution (Å) | 2.8 | 2.8 | 3.4 |
| Model resolution at FSC=0.5 (Å) | 2.91 | 2.92 | 3.56 |
| Map sharpening B factor (Å<sup>2</sup>)* | -100.7 | -95.4 | -82.0 |
| **Model composition** |  |  |  |
| Non-hydrogen atoms | 13,844 | 13,844 | 13,814 |
| Protein residues | 1,698 | 1,698 | 1,698 |
| Ligands | 2x PS | 2x PS | 1x PS |
| Waters | - | - | - |
| Ions | 10x Ca<sup>2+</sup> | 10x Ca<sup>2+</sup> | 10x Ca<sup>2+</sup> |
| **B factors (Å<sup>2</sup>)** |  |  |  |
| Protein (min/max/mean) | 50.51/380.99/150.8 | 62.73/343.1/159.57 | 53.66/419.51/156.01 |
| Ligand (min/max/mean) | 91.64/253.59/149.89 | 92.36/265.58/148.07 | 102.77/290.86/149.04 |
| Water | - | - | - |
| **R.m.s. deviations** |  |  |  |
| Bond lengths (Å) | 0.006 | 0.005 | 0.005 |
| Bond angles (°) | 0.952 | 1.080 | 0.910 |
| **Validation** |  |  |  |
| Molprobity score | 1.61 | 1.48 | 1.58 |
| Clashscore | 2.84 | 2.11 | 2.92 |
| Rotamer outliers (%) | 1.79 | 1.52 | 1.59 |
| Cβ outliers (%) | - | - | - |
| CaBLAM outliers (%) | 2.99 | 2.75 | 2.99 |
| **Ramachandran plot** |  |  |  |
| Favored (%) | 94.96 | 94.78 | 95.02 |
| Allowed (%) | 4.86 | 5.16 | 4.92 |
| Disallowed (%) | 0.18 | 0.06 | 0.06 |

\*Automatically determined B factor; the maps have been sharpened with LocScale and DeepEMhancer.

**Table S3.** Data collection, refinement, and validation statistics for the lipid-bound soluble myoferlin (1-1997): 15 mol% DOPS/5 mol% Cholesterol lipid nanodisc complex.

|  | Map M5<br>(consensus map) | Map M6<br>(overall map) |
| --- | --- | --- |
| **Data collection and processing** |  |  |
| Electron gun | X-FEG |  |
| Detector | Falcon4i |  |
| Magnification | 165,000 |  |
| Energy filter slit width (eV) | 10 |  |
| Voltage (kV) | 300.0 |  |
| Dose rate (e<sup>-</sup>/Å<sup>2</sup>/s) | 11.55 |  |
| Electron exposure on sample (e<sup>-</sup>/Å<sup>2</sup>) | 39.848 |  |
| Target defocus range (μm) | 0.8-2.0 |  |
| Calibrated pixel size (Å) | 0.72 |  |
| Symmetry imposed | C1 |  |
| Collected movies (no.) | 9,621 |  |
| Initial particle images (no.) | 2,917,286 |  |
| Final particle images (no.) | 277,729 | 38,088 |
| Map resolution at FSC=0.143 (Å) | 2.74 | 3.29 |
| **Refinement** |  |  |
| Initial model used (PDB code) | AlphaFold2 | AlphaFold2 |
| Model resolution (Å) | 2.9 | 3.4 |
| Model resolution at FSC=0.5 (Å) | 3.04 | 3.71 |
| Map sharpening B factor (Å<sup>2</sup>)* | -97.7 | -63.9 |
| **Model composition** |  |  |
| Non-hydrogen atoms | 13,844 | 13,844 |
| Protein residues | 1,698 | 1,698 |
| Ligands | 2x PS | 2x PS |
| Waters | - | - |
| Ions | 10x Ca<sup>2+</sup> | 10x Ca<sup>2+</sup> |
| **B factors (Å<sup>2</sup>)** |  |  |
| Protein (min/max/mean) | 46.64/388.86/135.43 | 62.29/407.28/166.62 |
| Ligand (min/max/mean) | 72.37/240.52/123.64 | 115.2/307.52/177.37 |
| Water | - | - |
| **R.m.s. deviations** |  |  |
| Bond lengths (Å) | 0.005 | 0.005 |
| Bond angles (°) | 0.953 | 1.058 |
| **Validation** |  |  |
| Molprobity score | 1.48 | 1.54 |
| Clashscore | 2.18 | 3.42 |
| Rotamer outliers (%) | 1.59 | 1.26 |
| Cβ outliers (%) | - | - |
| CaBLAM outliers (%) | 2.63 | 2.57 |
| **Ramachandran plot** |  |  |
| Favored (%) | 95.2 | 95.26 |
| Allowed (%) | 4.63 | 4.63 |
| Disallowed (%) | 0.18 | 0.12 |

\*Automatically determined B factor; the maps have been sharpened with LocScale and DeepEMhancer.

**Table S4.** Data collection, refinement, and validation statistics for the lipid-bound soluble myoferlin (1-1997): 15 mol% DOPS/2 mol% PI(4,5)P_2_ lipid nanodisc complex.

|  | Map M7<br>(consensus map) | Map M8<br>(overall map) |
| --- | --- | --- |
| **Data collection and processing** |  |  |
| Electron gun | X-FEG |  |
| Detector | Falcon4i |  |
| Magnification | 165,000 |  |
| Energy filter slit width (eV) | 10 |  |
| Voltage (kV) | 300.0 |  |
| Dose rate (e<sup>-</sup>/Å<sup>2</sup>/s) | 11.53 |  |
| Electron exposure on sample (e<sup>-</sup>/Å<sup>2</sup>) | 39.894 |  |
| Target defocus range (μm) | 1.1-2.0 |  |
| Calibrated pixel size (Å) | 0.72 |  |
| Symmetry imposed | C1 |  |
| Collected movies (no.) | 7,014 |  |
| Initial particle images (no.) | 2,484,379 |  |
| Final particle images (no.) | 583,145 | 28,855 |
| Map resolution at FSC=0.143 (Å) | 2.79 | 3.43 |
| **Refinement** |  |  |
| Initial model used (PDB code) | AlphaFold2 | AlphaFold2 |
| Model resolution (Å) | 2.9 | 3.6 |
| Model resolution at FSC=0.5 (Å) | 3.11 | 3.91 |
| Map sharpening B factor (Å<sup>2</sup>)* | -107.9 | -66.9 |
| **Model composition** |  |  |
| Non-hydrogen atoms | 13,844 | 13,844 |
| Protein residues | 1,698 | 1,698 |
| Ligands | 2x PS | 2x PS |
| Waters | - | - |
| Ions | 10x Ca<sup>2+</sup> | 10x Ca<sup>2+</sup> |
| **B factors (Å<sup>2</sup>)** |  |  |
| Protein (min/max/mean) | 63.62/441.07/154.78 | 66.21/342.59/127.67 |
| Ligand (min/max/mean) | 106.11/238.96/152.08 | 93.97/210.68/131.32 |
| Water | - | - |
| **R.m.s. deviations** |  |  |
| Bond lengths (Å) | 0.005 | 0.006 |
| Bond angles (°) | 1.028 | 1.187 |
| **Validation** |  |  |
| Molprobity score | 1.37 | 1.63 |
| Clashscore | 2.07 | 2.62 |
| Rotamer outliers (%) | 1.32 | 2.05 |
| Cβ outliers (%) | - | 0.06 |
| CaBLAM outliers (%) | 2.57 | 2.96 |
| **Ramachandran plot** |  |  |
| Favored (%) | 95.67 | 94.96 |
| Allowed (%) | 4.09 | 4.74 |
| Disallowed (%) | 0.24 | 0.3 |

\*Automatically determined B factor; the maps have been sharpened with LocScale and DeepEMhancer.

**Table S5.** Data collection, refinement, and validation statistics for the lipid-free soluble myoferlin.

|  | Map M9<br>(consensus map) | Map M10<br>(Fer<sup>core</sup>) | Map M11<br>(overall map) |
| --- | --- | --- | --- |
| **Data collection and processing** | Dataset 1 | Dataset 2 |  |
| Electron gun | X-FEG | X-FEG |  |
| Detector | Falcon4i | Falcon4i |  |
| Magnification | 165,000 | 165,000 |  |
| Energy filter slit width (eV) | 10 | 10 |  |
| Voltage (kV) | 300.0 | 300.0 |  |
| Dose rate (e<sup>-</sup>/Å<sup>2</sup>/s) | 11.67 | 10.8 |  |
| Electron exposure on sample (e<sup>-</sup>/Å<sup>2</sup>) | 39.911 | 39.852 |  |
| Target defocus range (μm) | 1.1-2.3 | 0.8-2.0 |  |
| Calibrated pixel size (Å) | 0.72 | 0.72 |  |
| Symmetry imposed | C1 | C1 |  |
| Collected movies (no.) | 10,354 | 8,669 |  |
| Initial particle images (no.) | 6,088,884 | 3,020,482 |  |
| Final particle images (no.) | 102,339 | 298,755 | 18,157 |
| Map resolution at FSC=0.143 (Å) | 3.21 | 2.81 | 8.5 |
| **Refinement** |  |  |  |
| Initial model used (PDB code) | AlphaFold2 |  |  |
| Model resolution (Å) | 3.4 |  |  |
| Model resolution at FSC=0.5 (Å) | 3.73 |  |  |
| Map sharpening B factor (Å<sup>2</sup>)* | -91.9 |  |  |
| Model composition |  |  |  |
| Non-hydrogen atoms | 10,833 |  |  |
| Protein residues | 1,340 |  |  |
| Ligands | - |  |  |
| Waters | - |  |  |
| Ions | 6x Ca<sup>2+</sup> |  |  |
| *B* factors (Å<sup>2</sup>) |  |  |  |
| Protein (min/max/mean) | 29.97/419.31/167.9 |  |  |
| Ligand (min/max/mean) | 95.77/336.41/180.97 |  |  |
| Water | - |  | - |
| R.m.s. deviations |  |  |  |
| Bond lengths (Å) | 0.005 |  |  |
| Bond angles (°) | 1.002 |  |  |
| **Validation** |  |  |  |
| Molprobity score | 1.51 |  |  |
| Clashscore | 3.06 |  |  |
| Rotamer outliers (%) | 1.34 |  |  |
| Cβ outliers (%) | - |  |  |
| CaBLAM outliers (%) | 2.74 |  |  |
| Ramachandran plot |  |  |  |
| Favored (%) | 95.48 |  |  |
| Allowed (%) | 4.45 |  |  |
| Disallowed (%) | 0.08 |  |  |

\*Automatically determined B factor; the maps have been sharpened with LocScale and DeepEMhancer.

**Table S6.** Data collection, refinement, and validation statistics for the lipid-free soluble dysferlin.

|  | Map M1<br>(Fer<sup>core</sup> map) | Map M2<br>(closed state) | Map M3<br>(overall map) |
| --- | --- | --- | --- |
| **Data collection and processing** | Dataset 1 | Dataset 2 |  |
| Electron gun | X-FEG | XFEG |  |
| Detector | Falcon4i | Falcon4i |  |
| Magnification | 165,000 | 165,000 |  |
| Energy filter slit width (eV) | 15 | 10 |  |
| Voltage (kV) | 300.0 | 300.0 |  |
| Dose rate (e<sup>-</sup>/Å<sup>2</sup>/s) | 13.35 | 13.59 |  |
| Electron exposure on sample (e<sup>-</sup>/Å<sup>2</sup>) | 40.184 | 40.227 |  |
| Target defocus range (μm) | 0.8-2.3 | 0.8-2.3 |  |
| Calibrated pixel size (Å) | 0.72 | 0.72 |  |
| Symmetry imposed | C1 | C1 |  |
| Collected movies (no.) | 4,582 | 4,727 |  |
| Initial particle images (no.) | 1,660,317 | 1,201,065 |  |
| Final particle images (no.) | 230,540 | 29,793 | 11,477 |
| Map resolution at FSC=0.143 (Å) | 2.92 | 3.54 | 4.76 |
| **Refinement** |  |  |  |
| Initial model used (PDB code) |  | AlphaFold2 |  |
| Model resolution (Å) |  | 3.6 |  |
| Model resolution at FSC=0.5 (Å) |  | 3.81 |  |
| Map sharpening B factor (Å<sup>2</sup>)* |  | -56.9 |  |
| Model composition |  |  |  |
| Non-hydrogen atoms |  | 11,100 |  |
| Protein residues |  | 1,370 |  |
| Ligands |  | - |  |
| Waters |  | - |  |
| Ions |  | 6x Ca<sup>2+</sup> |  |
| *B* factors (Å<sup>2</sup>) |  |  |  |
| Protein (min/max/mean) |  | 45.60/370.52/173.98 |  |
| Ligand (min/max/mean) |  | 75.34/297.09/150.49 |  |
| Water |  | - |  |
| R.m.s. deviations |  |  |  |
| Bond lengths (Å) |  | 0.004 |  |
| Bond angles (°) |  | 0.982 |  |
| **Validation** |  |  |  |
| Molprobit score |  | 1.54 |  |
| Clashscore |  | 2.68 |  |
| Rotamer outliers (%) |  | 1.57 |  |
| Cβ outliers (%) |  | - |  |
| CaBLAM outliers (%) |  | 2.19 |  |
| Ramachandran plot |  |  |  |
| Favored (%) |  | 95.17 |  |
| Allowed (%) |  | 4.75 |  |
| Disallowed (%) |  | 0.07 |  |

\*Automatically determined B factor; the maps have been sharpened with LocScale and DeepEMhancer.

**Table S7.**
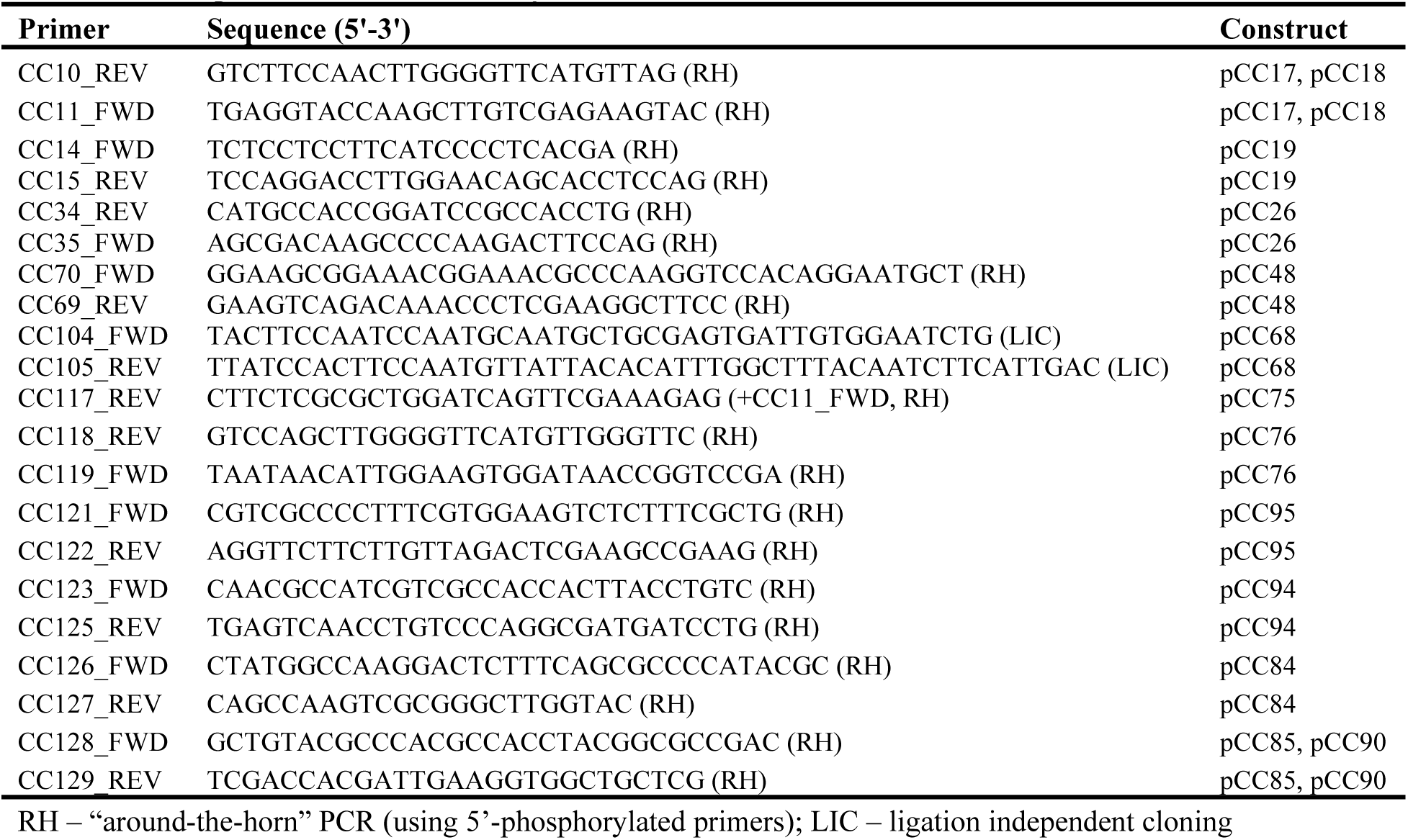
PCR primers used in this study.

